# Pubertal hormones and the early adolescent female brain: a multimodality brain MRI study

**DOI:** 10.1101/2024.11.27.625122

**Authors:** Muskan Khetan, Nandita Vijayakumar, Ye Ella Tian, Megan M. Herting, Michele O’ Connell, Marc Seal, Sarah Whittle

## Abstract

Puberty is a critical developmental process that is associated with changes in steroid hormone levels, which are believed to influence adolescent behaviour via their effects on the developing brain. So far, there are limited and inconsistent findings regarding the relationship between steroid hormones and brain structure and function in adolescent females, with many existing studies employing small sample sizes. Thus, in this study, we explored the association between oestradiol (E2), testosterone (Tes) and dehydroepiandrosterone (DHEA) and brain structure (gray matter volume, sulcal depth, cortical thickness and white matter microstructure) and function (resting-state connectivity, emotional n-back task-related function) in 3024 adolescent females (age 8.92 - 13.33 years, mean age (SD) = 10.37 (0.94) years) from the Adolescent Brain Cognitive Development^SM^ (ABCD^®^) Study. We used elastic-net regression with cross-validation to investigate associations between hormones and brain phenotypes derived from multiple imaging modalities. We found that structural brain features, including cortical thickness, sulcal depth, and white matter microstructure, were among the most important features associated with hormones. E2 was most strongly associated with prefrontal and premotor regions involved in working memory and emotion processing, while Tes and DHEA were most strongly associated with parietal and occipital regions involved in visuospatial functioning. All three hormones were also associated with prefrontal, temporoparietal junction and insula cortices. Thus, using an advanced methodological approach, this study suggests both unique and overlapping neural correlates of pubertal hormones in adolescent females and sheds light on the mechanisms by which puberty influences adolescent development and behaviour.

## 1. INTRODUCTION

Puberty is considered one of the most significant developmental processes during adolescence, influencing social, emotional and behavioural changes via effects on the developing brain (Dai and Scherf, 2019; Vijayakumar et al., 2018a). Pubertal influences on the developing brain also differ between males and females, thought to contribute to sex differences in socioemotional functioning and mental health (Colleen S. et al., 2012; Rose et al., 2006). In particular, female adolescents are approximately 2-3 times more likely to experience emotion dysregulation, anxiety and depression (Campbell et al., 2021; Breslau et al., 2017; Hulvershorn et al., 2011; Lewinsohn, Gotlib, et al., 1998; Kessler et al., 1994; 1995; Nolen-Hoeksema & Girgus, 1994), with sex differences emerging around puberty (Hayword & Sanborn, 2002; Alloy et al., 2016). Pubertal hormones are thought to impact the structure and function of the developing brain via steroid receptors (Laube et al., 2020; Goddings et al., 2019; Schultz et al., 2009), particularly in parts of the brain that contribute to emotion processing and regulation such as prefrontal cortex (PFC) and limbic system (Gruber et al., 2002; Brinton et al., 2008). As such, understanding how pubertal hormones impact the female brain during puberty may shed light on risk mechanisms for emotion dysregulation and associated poor mental health outcomes.

‘True puberty’ begins with gonadarche, when the activation of the hypothalamic-pituitary-gonadal (HPG-axis) leads to an increase in sex hormone levels that ultimately support reproductive maturity (Schulz et al., 2009; Sisk and Foster, 2004). Oestrogens, including oestradiol (E2), are key female sex hormones. E2 binds with E2 receptors (ERα and Erβ: ERs) that are highly expressed across brain regions responsible for emotional regulation, such as the prefrontal cortex (PFC), limbic system and hypothalamus (Gruber et al., 2002; Brinton et al., 2008). Both animal studies and post-mortem human research confirm E2’s presence in these regions, highlighting its impact on brain structure and function (Sisk & Zehr, 2005; Koss et al., 2015; Bixo et al., 1986, 1995, 1997).

In addition, neuroimaging studies have shown the widespread effects of E2 on brain structure and function during adolescence, although the regions and direction of effects have been inconsistent (Vijayakumar et al., 2018).

For example, existing studies have found higher E2 in adolescent females to be associated with a reduced whole brain gray matter (GM) volume and increased amygdala volume over time (Herting et al., 2014), lower anterior cingulate cortex volume (ACC; Koolschijn et al., 2014), higher parahippocampal and uncus gray matter (Neufang et al., 2009), increased gray matter density in the middle frontal, inferior temporal, and middle occipital gyri but decreased GM density and volume in left frontal and parietal regions (Peper et al., 2009; Brouwer et al., 2015). In addition, E2 has been associated with white matter (WM) volume and microstructure. For example, studies have reported higher E2 levels to be associated with higher fractional anisotropy (FA; reflecting stronger structural integrity) in the lateral, middle occipital cortex and left uncinate fasciculus (Herting et al., 2012; Ho et al., 2020), increased whole brain WM volume over time and lower FA in angular gyrus and superior longitudinal fasciculus (Herting et al., 2014; 2012).

Functionally, E2 levels have been associated with brain function during emotion-processing tasks. For example, higher E2 levels were associated with an increased anterior temporal cortex (Goddings et al., 2012; Klapwijk et al., 2013) and insula activity (Macks et al., 2017) during social-emotional processing, as well as higher and lower activity in the occipital cortex and cerebellum, respectively, during an emotion-cognition task (Cservenka et al., 2016). E2 levels were also associated with dorsolateral prefrontal cortex (DLPFC) activity during emotional regulation (Chung et al., 2019). In summary, despite some evidence connecting E2 levels to brain structure and function in adolescent females, findings remain inconsistent, and large-scale studies are lacking.

In addition to E2, there are other pubertal hormones that likely influence behavioural outcomes through their influence on the developing brain (Schulz et al., 2016; Vigil et al., 2016; Peper and Dahl., 2013). These hormones specifically increase in levels during adrenarche and continue during gonadarche and beyond (Witchel et al., 2020; Havelock et al., 2004). Adrenarche involves the activation of the hypothalamus-pituitary-adrenal (HPA) axis, leading to the release of adrenal hormones such as dehydroepiandrosterone (DHEA) from adrenal glands (<u>Rosenfield.,</u> <u>2021</u>). DHEA subsequently acts as a precursor for testosterone (Tes), which is later released from the ovaries and other target tissues (Lebbe and Woodruff., 2013). Molecularly, in females, DHEA binds with androgen receptors (ARs) and ERs after conversion to Tes and oestrogens via aromatase enzyme within cells (Ubuka and Tsutsui, 2014; Kawata, 1995), as well as act directly on ARs in the brain (Lu et al., 2003). Both ERs and ARs are widely available across the whole brain (Lloyd-Evans et al., 2020; Sisk et al., 2005; Nunez et al., 2003).

Similarly to E2, prior findings of associations between Tes and DHEA and brain structure and function in adolescent females have been mixed (Barendse et al., 2018; Bramen et al., 2012; Cedric et al., 2014; Herting et al., 2012; Ho et al., 2020; Goddings et al., 2019; Klauser et al., 2015; Nguyen et al., 2013; Peper et al., 2015; Vijayakumar et al., 2019; Wierenga et al., 2018). For example, prior neuroimaging studies have shown that higher Tes levels have been linked to smaller hippocampal volumes (Wierenga et al., 2018), and higher Tes and DHEA levels have also been linked with thinner cortex in limbic, occipital, and superior temporal regions (Vijayakumar et al., 2019; Cedric et al., 2014; Bramen et al., 2012) among females. In addition, Tes showed a shift from positive to negative association with cortical thickness in the somatosensory cortex and posterior cingulate with advanced puberty in females (Nguyen et al., 2013). Findings on androgens’ association with WM are similarly inconsistent. In one study, Tes was positively related to FA in only a few WM regions (e.g., precentral gyrus; Herting et al., 2012) while another study has shown Tes to be positively associated with FA in diverse WM tracts (including the corpus callosum, bilateral cingulum bundle, and bilateral corticospinal tracts; Ho et al., 2020) among females. Additionally, DHEA was also related to lower frontal WM volume (Klauser et al., 2015) and higher mean diffusivity (MD, reflecting weaker structural integrity) in diverse WM tracts (Barendse et al., 2018) in both males and females.

DHEA and Tes have also shown associations with brain activity in response to emotional stimuli in adolescents. For example, higher DHEA levels were found to relate to decreased activation in the cingulate cortex, insula, striatum, and DLPFC in one study (Whittle et al., 2015), while higher Tes levels were associated with nonlinear increases of activity in amygdala and hippocampus in another study (Vijayakumar et al., 2019). In both studies, associations were apparent in females but not males. In summary, current research suggests that androgens are associated with both brain structure and function in females, however, findings are still limited and largely inconsistent.

To gain a comprehensive understanding of the influence of pubertal process (both adrenarche and gonadarche) on the developing female brain, it is important to investigate these three pubertal hormones. So far, a systematic understanding of how steroid hormones influence the brain during adolescence, particularly in early adolescent female, is still lacking. In addition, research to date has not yet looked at the influence of steroid hormones on brain structure and function simultaneously within the same cohort using the same analytical approach. Utilising different imaging modalities, analytic approaches and samples could all be contributing to inconsistent findings in the literature. This indicates a need to investigate hormone-brain associations across different brain imaging modalities (structural and functional) within the same sample.

### 1.1 Current study

In the current study, we aimed to comprehensively investigate the associations between steroid hormone levels and brain structure and function in a large sample of early adolescent females, when E2 levels begin to increase (Sizonenko., 1978). We aimed to build a multimodal model using cross-validation (CV) to assess the relative contributions of different brain imaging modalities. The use of a CV framework was intended to ensure robust results, minimizing the addition of inconsistent findings to the existing literature. We hypothesised that all three hormones (E2, Testosterone, DHEA) would be associated with brain gray and white matter morphology, functional activity (with a focus on emotion-related brain function), and resting-state connectivity. However, given that there are inconsistent findings in the literature, we did not make any regional or directional hypotheses.

## 2. <u>Methodology</u>

### 2.1 Study design

We utilised data from the ABCD® Study (open-science dataset version 4.0; released 2022), a 10- year longitudinal study of adolescent development that includes data from 22 sites across the United States (www.ABCDStudy.org). Each site obtained consent from the children and their parent(s)/guardians following local Institutional Review Boards (for further details related to study, refer Garavan et al., 2018; Jerniga et al., 2018; Volkow et al., 2018). For the current study, we used data from baseline (aged 9-11 years) and second-year follow-up (aged 11-13 years) for female participants alone.

### 2.2 Exclusion/Inclusion criteria

Full exclusion and inclusion criteria for the main ABCD study, refer to ABCD release notes 4.0 (DOI: 10.15154/1523041). Participants were included if they had data for all hormones (i.e., E2, Tes, DHEA) at either baseline or second-year follow-up. Participants with one or more poor-quality MRI modality data (N ∼ 2332) were excluded (following ABCD’s quality recommendations, using the variable “abcd_imgincl01”; details are in ABCD release notes 4.0).

For the current analyses, a cross-sectional design was utilised, whereby for any one participant, only hormone and imaging data from one-time point (baseline or second-year follow-up) was randomly selected. The random selection ensured equal percentages of participants were drawn from the total sample available at each wave. The final sample comprised 3024 females from 2789 families (number of siblings = 235), aged 9-13 years old. Of these, 2366 were from baseline, and 658 were from the second-year follow-up, with the sub-samples showing demographic similarity (refer to Supplementary Table 2.5).

### 2.3 Hormone measures

E2, Tes, and DHEA were measured from salivary samples (Uban et al., 2018). Participants were asked to provide saliva samples via passive drool with the help of trained research assistants in the laboratory. 30-min prior to the saliva collection, participants were instructed not to have any food or drink, and 60-min prior to collection, they were asked not to have any major meals. The protocol for saliva collection was adapted from Granger and colleagues (2012). All samples were placed into a Nalgene Labtop Cooler and kept inside a small lunchbox cooler immediately after collection. All saliva samples were transferred to an on-site freezer (-80 to -20 degree Celsius) or within a cooler placed inside a refrigerator (4 degrees Celsius) according to the location of the collection site for 2-6 months, and then were shipped to Salimetrics (Carlsbad, CA) on dry ice. To assay the saliva samples, a commercially available immunoassay (following the protocol provided by the Salimetrics manufacturer) specifically designed for the saliva samples was used. Range of sensitivity for each hormone is as follows: E2 = 1 – 32 pg/ml, Tes = 6.1 - 600 pg/ml , DHEA = 10.2 - 1000 pg/ml (refer Herting et al., 2013 for further details) In addition, to avoid multiple freeze-thaw cycles of the saliva samples, all hormones were assayed in duplicates within a single day.

#### 2.3.1 Quality check

A decision tree protocol (Herting et al., 2021) was followed to ensure quality control (e.g., removing samples with any quality concerns based on research assistant notes or mismatched reports of biological sex). To obtain a single value for each hormone, hormone levels were averaged from each participant’s hormone replicate. To ensure robustness, intra-assay variance was also calculated, with the averaged intra-assay variance for all participants reported in Supplementary Section 2.4. In addition, correlations with confounding factors that may affect salivary hormone levels such as caffeine intake (yes/no), physical activity (yes/no), collection duration (time from starting and ending saliva collection), collection time (from midnight), time from collection to freeze, and age were also examined. There were significant correlations of the hormones with age (Tes r=0.44, DHEA r=0.29, E2=0.15, p’s<0.001; Figure 1) and caffeine intake (Tes r=0.1, DHEA r=0.055, E2 r=0.13, p’s<0.002; Supplementary Figure 2). However, the correlations with caffeine intake were not significant for E2 and DHEA after controlling for age (see Supplementary section 2.2). In addition, E2 had a significant correlation with physical activity (r = 0.055, p = 0.002) and collection time (r = 0.136, p < 0.001), however, only the correlation with collection time remained significant after controlling for age (r = 0.145, p < 0.001).

**Figure 1:**
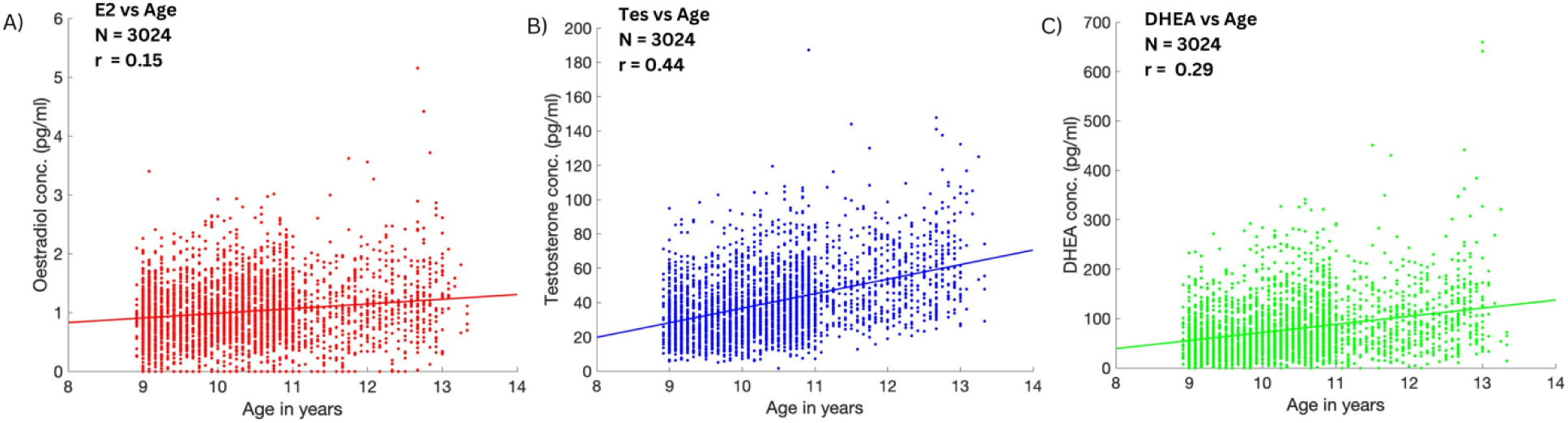
A relationship between steroid hormone levels and age A) E2 conc in pg/ml B) Tes in pg/ml C) DHEA in pg/ml distribution across age (in years); fitted line represents r or pearson’s correlation between each of the hormones and age

### 2.4 Neuroimaging measures

#### 2.4.1 Image acquisition and preprocessing

For all brain imaging measures, the current study has used the available pre-processed data by the ABCD team. Details related to scanning parameters, imaging processing and task paradigms have been previously described (Casey et al., 2018; Hagler et al., 2019). ABCD brain imaging data were collected using 3T scanners from three different manufacturers (*Siemens Prisma, General Electric (GE) 750 and Philips*) across 21 data collection sites. All participants were scanned using T1- and T2-weighted MRI, diffusion-weighted imaging (DWI), resting-state functional (rsfMRI), and task-based fMRI sequences.

#### 2.4.2 sMRI

Cortical regions were parcellated from the T1-weighted data using FreeSurfer v7.1.1, and the Destrieux atlas (Dale et al., 1999; Fischl et al., 1999b; Destrieux et al., 2010), which contained 148 bilateral cortical regions of interest (ROIs). Fourteen bilateral subcortical ROIs were parcellated using FreeSurfer’s subcortical segmentation (Fischl et al., 2002). Cortical structural measures included thickness (CTh), surface area (CAr), volume (CVol), and sulcal depth (SDep) and subcortical structural measures include volume (SCVol).

#### 2.4.3 DWI

In the ABCD study, all diffusion images were pre-processed for head movements, eddy current distortions, B0 distortion, and gradient nonlinearity distortions (refer Hagler et al., 2019 for details on each of these steps). Then, diffusion measures such as fractional anisotropy (FA) and mean diffusivity (MD) were computed using a DTI model of diffusion. For the current study, only b values equal to 500 & 1000 were included for the tensor fitting models (Hagler et al., 2019; Casey et al., 2018).

Thirty-seven bilateral white matter (WM) fibre tracts were parcellated using AtlasTrack atlas, a probabilistic atlas-based method for automated segmentation of white matter fibre tracts (Hagler et al., 2009; 2019). Details on the selected networks from AtlasTrack are shown in Supplementary Section 2.3.

#### 2.4.4 rsfMRI

Estimated resting state activity was collected while the participant remained at rest (looking at the fixation cross with eyes open) across four runs (5 minutes per run; Casey et al., 2018). Then, resting state connectivity was calculated between and within cortical networks and between subcortical regions and cortical networks. Twelve cortical networks were defined based on the Gordon parcellation (Gordon et al., 2016). Average pairwise correlations were then calculated within (intra) and between (inter) each cortical network. Average pairwise correlations were also calculated between each cortical network and each FreeSurfer-defined subcortical region. This resulted in 126 inter-cortical networks, 12 intra-cortical networks, and 199 cortical networks to subcortical region correlations. The correlation values were transformed to z-statistics for the final analysis. Finally, the Fisher Z transformation of all the correlation values was examined (Hagler et al., 2019).

#### 2.4.5 EN-back task

The EN-back task assesses working memory and emotional processing (Casey et al., 2018). This task required participants to observe a sequence of pictures (i.e., faces and places) and then determine if the current image matches one shown in previous blocks, either immediately prior (0-back) or two images back (2-back). Given the focus of this study on emotional processing, only relevant contrasts related to emotional processing were selected: faces vs places (FvP), emotional faces vs neutral faces (EvN), positive faces vs neutral faces (PvN), and negative faces vs neutral faces (NvN). Average activity across each of the FreeSurfer segmented Destrieux cortical and subcortical ROIs was extracted for each contrast and used in subsequent analyses (Hagler et al., 2019).

### 2.5 Analytic approach 2.5.1 Elastic-net regression

To understand the association of each of the hormones with brain structure and function features, Elastic-Net (ENet) regression from the GLMnet package from MATLAB (Qian et al., 2013) was utilised, with hormone levels as the outcome variable and brain measures as predictors. ENet regression is a combination of lasso (L1) and ridge (L2) linear regression, specified by the hyperparameter alpha and lambda values whereby alpha = 0 is ridge while alpha = 1 is lasso. ENet regression is considered to be preferred over other multivariate methods (such as random forest, and supervision learning) when a large number of correlated features predict the outcome (Caunca et al., 2021).

#### 2.5.2 Feature Selection Method

Although we aimed to understand structural and functional (i.e., multi-modal) correlates of hormones, given the large number of structural and functional features, a single-modality approach was first adopted to select features to then feed into the multi-modal model. This approach served to improve the multi-modal model performance by removing sources of noise and reducing data dimensionality. Thus, ENet regression within a machine learning algorithm was applied to each MRI modality separately (i.e., four separate unimodal models for each hormone). From each model, features (regional measures of structural or functional modality) that explained variance (non-zero weights) in each hormone (E2, Tes, and DHEA) were selected for the multi-modality model.

#### 2.5.3 Machine Learning Framework

First, the sample was randomly split using 10-fold cross-validation (CV), each fold including 10% of the data, ensuring that in the case of siblings, each sibling from a given family remained in the same fold to avoid data leakage. Nine out of ten folds were randomly selected for training while the held-out fold was used for testing and evaluation of the model performance.

Before model training, all the structural and functional MRI features were harmonised with age as a covariate across multi-scanner sites using ComBat Harmonisation (Orlhac et al., 2022). Harmonisation was performed separately for training and test data to avoid any data leakage.

While fitting the models, hyperparameters were selected using a grid search for the alpha value within the 10-nested folds (script adapted from https://github.com/owensmax/ADHD). The best model was selected as that with the minimum mean square error (MSE) and used for predicting the outcome for the remaining (test) fold. The complete pipeline was repeated across 10 iterations to increase the robustness of the selected features. Thus, only features with non-zero values across the 10-iterations were selected. Average performance metrics (R-square and correlation) from all 10 iterations were also calculated and reported only for multi- modality models (this is explained in a detailed flowchart in Figure 2).

**Figure 2:**
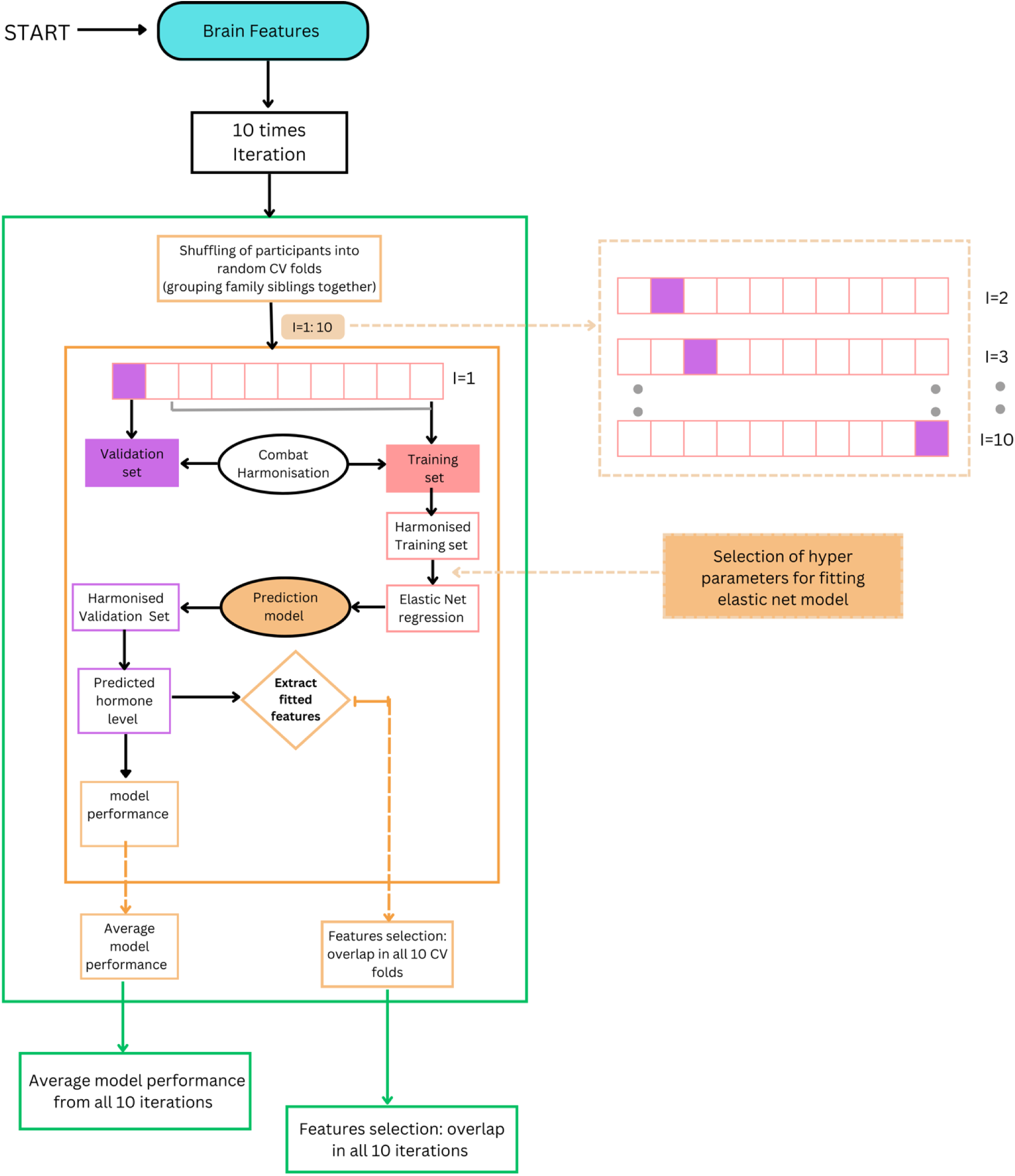
Schematic of machine learning and feature selection framework. The final selection of features from each modality involved repeating the same process within the green box 10- times. Each iteration involved fitting the ENet regression model following the sample cross-validation method to select out hyper-parameters, computing model performance using a validation set, and extracting non-zero weighted features Feature selection was based on the overlap of the same non-zero weighted feature across all 10-iterations. Finally, average model performance and average beta weight for selected features were computed for 10-iterations.

#### 2.5.4 Multi-modality MRI model

Using features input from uni-modal models, three separate multi-modality models, one for each hormone, were run. Similar to the uni-modality models, an ENet regression within the machine-learning framework was implemented. Average performance matrices and fitted features were extracted for brain features that were present across the 10 iterations (Figures 2 & 3).

#### 2.5.5 Sensitivity analyses

In primary analyses, participant age was not controlled, as this could remove meaningful variance in hormone levels. In addition, there were some hormone outliers (outside of the mean +/- 3*SD) and significant correlation between Tes and caffeine intake. Thus, we ran multiple sensitivity analyses, which included a) removing the effects of age from each of the hormone levels, b) removing caffeine consumption from Tes levels, c) removing collection time from E2 levels, and d) removing hormone outliers. Results from these analyses are in the Supplementary Material.

## 3. Results

### 3.1 Descriptive statistics

Demographics: Demographic details collected at baseline and second-year follow-up are presented in Table 2.

**Table 2:** Demographic and pubertal characteristics of participants

|  |  |
| --- | --- |
| **Age in years** |  |
| Range | 8.92 - 13.33 years |
| Mean (SD) | 10.37 (0.94) |
| **Salivary hormone levels (pg/ml)** | **Mean (SD)** |
| E2 | 0.99 (0.49) |
| Tes | 38.87 (18.77) |
| DHEA | 74.17(49.89) |
| **BMI (percentile)<sup>b</sup>** | **n (%)** |
| Underweight (<5<sup>th</sup> percentile) | 102 (3.4) |
| Normal weight (>5<sup>th</sup> to <85<sup>th</sup> ) | 1711 (56.6) |
| Overweight (>85<sup>th</sup> to <95<sup>th</sup> ) | 407 (13.45) |
| Obese (>=95<sup>th</sup> ) | 364 (12.04) |
| Missing | 407 (13.46) |
| **Race/Ethnicity<sup>a</sup>** | **n (%)** |
| Non-Hispanic White | 1693 (55.98) |
| Non-Hispanic Black | 320 (10.58) |
| Hispanic | 586 (19.38) |
| Non-Hispanic Asian | 84 (2.77) |
| Other<sup>c</sup> | 341 (11.28) |
| **Parent Education<sup>a</sup>** | **n (%)** |
| <HS | 153 (5.06) |
| HS/GED | 249 (8.23) |
| Some college | 461 (15.24) |
| Associate degree (occupational/academic program) | 338 (11.18) |
| Bachelor's degree | 948 (31.35) |
| Postgraduate degree | 874 (28.90) |
| Missing | 2 (0.07) |
| **Menarche<sup>a</sup>** | **n (%)** |
| Pre-menarchal | 2715 (89.78) |
| Post-menarchal | 309 (10.22) |
| Age at menarche in years; mean (SD) | 11.905 (0.94) |
| **Reported medication/steroid use<sup>a</sup>** | **n (%)** |
| Yes | 3 (0.09) |
| No | 306 (10.11) |
| Missing | 2716 (89.78) |
| **Tanner stage<sup>a</sup>** | **n (%)** |
| 1: pre-puberty | 371 (12.27) |
| 2: early puberty | 1128 (37.3) |
| 3: mid puberty | 757 (25.03) |
| 4: late puberty | 366 (12.10) |
| 5: post-puberty | 208 (6.88) |

- a. Indicates parent-reported measure - b. Calculated using height in metres and weight in kgs based on the individual's age and biological sex provided by the 2000 CDC growth charts. - c. Other includes those who have indicated more than one race, and those that indicated American Indian, Native American, Alaska Native, Native Hawaiian, Guamanian, Samoan, other Pacific Islander or other race.

<u>Puberty</u>: To describe the pubertal status of participants, we included youth-reported measures of their child’s physical development using the Pubertal Development Scale (PDS; Petersen et al., 1988). For interpretability, we converted PDS scores to Tanner stages (following Shirtcliff et al., 2009). General Tanner stages (average of adrenal and gonadal score) are included in Table 2. For the ranges of each of the hormones in each of these categories, refer to Supplementary Table 2.6. The mean and standard deviation for each hormone were also calculated for the final sample and are presented in Table 2. For minimum, maximum levels and IQR for each of the hormones; refer to Supplementary sectional 2.4 and for the hormone distribution across the study sample; refer to Supplementary Figure 3. In addition, for the correlation among the hormones; refer Supplementary Section 2.1 and Supplementary Figure 1.

### 3.2. Unimodality Analysis

Results from unimodality analyses are presented in Table 3 and detailed results on the selected features and effect sizes are in Supplementary Section 3.1.1.

**Table 3:**
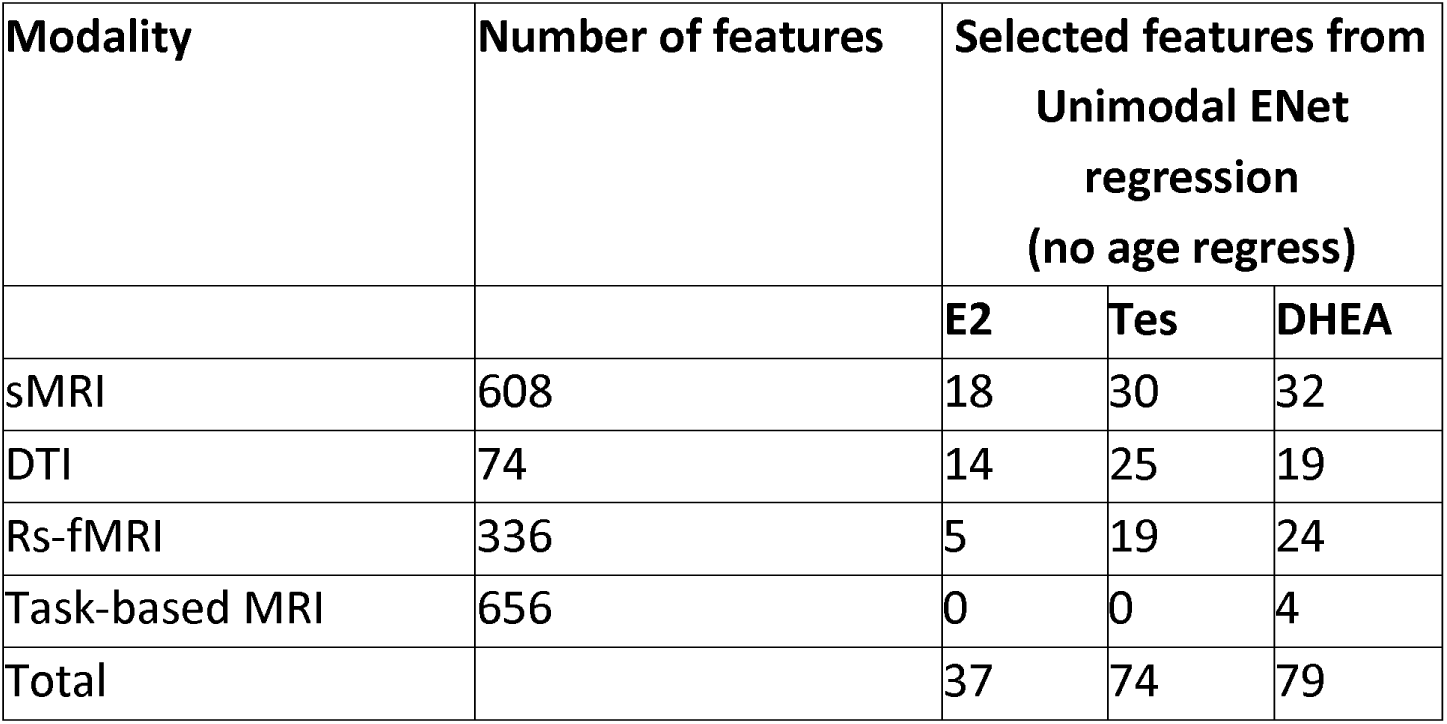
number of the selected features from unimodality models.

| Modality | Number of features | Selected features from Unimodal ENet regression (no age regress) |  |  |
| --- | --- | --- | --- | --- |
|  |  | E2 | Tes | DHEA |
| sMRI | 608 | 18 | 30 | 32 |
| DTI | 74 | 14 | 25 | 19 |
| Rs-fMRI | 336 | 5 | 19 | 24 |
| Task-based MRI | 656 | 0 | 0 | 4 |
| Total |  | 37 | 74 | 79 |

#### 3.2.1 sMRI

There were 18 structural features (1 CAr, 10 CTh, 1 CVol, 5 SDEP and 1 SCVol) associated with E2, 32 (2 CAr, 14 CTh, 6 CVol, 8 SDep and 2 SCVol) with DHEA and 30 (16 CTh, 2 CVol, 8 SDep and 4 SCVol) with Tes. E2 showed associations across the frontal, occipital, and temporal lobe while Tes and DHEA showed associations across the whole brain. Of subcortical regions, all three hormones showed positive associations, except for a negative association between Tes and accumbens area. See Supplementary Materials for the complete list.

#### 3.2.2 dMRI

Out of 74 diffusion metrics features, there were 14 (7 FA and 7 MD) associated with E2, 25 (17 FA and 8 MD) with Tes and 19 (13 FA and 6 MD) with DHEA. The associations were both positive and negative with FA and MD within the WM tracts across the entire brain for both Tes and DHEA. However, E2 showed a majority of the associations within WM tracts connecting the frontal lobe and other brain regions. See Supplementary Materials for the complete list.

#### 3.2.3 rsfMRI

For rsfMRI, there were 5 features associated with E2, 19 with Tes, and 24 with DHEA. For E2, all 5 features were subcortical to cortical network correlations; for Tes, 13 features were subcortical to cortical network correlations, 5 features were inter-cortical network correlations while 1 feature was an intra-cortical network correlation; for DHEA, 17 features were subcortical to cortical network correlations, 5 features were inter-cortical network correlations while 2 features were intra-cortical network correlations.

#### 3.2.4 EN-back task-based fMRI

Out of 656 functional features, only 4 features were associated with DHEA, while no features were associated with E2 and Tes. For DHEA, 2 features reflected activity associated with positive emotion processing (PvN contrast), 1 feature reflected activity associated with face processing (FvP contrast), and 1 was activity associated with negative emotion processing (NvN contrast).

### 3.3 Multimodality Analyses

Taking all the selected features from unimodality models, a multimodal model was computed for each hormone separately. Computational statistics are presented in Table 4 and the list of all the selected features is presented in Table 5.

**Table 4:** R2 and r values from the final fitted multimodal ENet for each hormone

| Hormones | No Age-regress |  |  |
| --- | --- | --- | --- |
|  | Selected Features | R2 value | r |
| E2 | 31 | 0.03 | 0.24 |
| Tes | 71 | 0.14 | 0.41 |
| DHEA | 62 | 0.07 | 0.32 |

**Table 5:** Selected Features from multimodal ENet regression for E2

| Regions | hemisphere | Features | Beta Weights |
| --- | --- | --- | --- |
| Intercept |  |  | 1.015 |
| FMIN | - | MD | 0.044 |
| Default mode network and left thalamus proper | left | Resting state connectivity between cortical network and subcortical region | -0.035 |
| G and S paracentral | left | SDep | -0.033 |
| G and S frontomarginal | left | CTh | -0.027 |
| SCSLH | left | FA | 0.027 |
| ILFLH | left | MD | -0.027 |
| CSTLH | left | MD | 0.027 |
| Pole occipital | left | CTh | 0.026 |
| vertical ramus of the anterior segment of the lateral sulcus | left | CTh | 0.026 |
| Retrosplenial Temporal network and left putamen | left | Resting state connectivity between cortical network and subcortical region | 0.025 |
| TSLFRH | right | FA | -0.025 |
| ATRLH | left | MD | -0.025 |
| horizontal ramus of the anterior segment of the lateral sulcus | left | SDep | -0.025 |
| G planum temporale | left | SDep | -0.023 |
| S occipital anterior | left | CVol | -0.023 |
| cinguloopercular and right pallidum | right | Resting state connectivity between cortical network and subcortical region | -0.022 |
| Retrosplenial temporal network and right caudate | right | Resting state connectivity between cortical network and subcortical region | 0.022 |
| G and S transverse frontopolar | left | CTh | -0.021 |
| CGCLH | left | MD | -0.020 |
| S circular insula anterior | right | CTh | -0.019 |
| Sensory motor hand network and left caudate | left | Resting state connectivity between cortical network and subcortical region | -0.019 |
| S collat transv post | right | CAR | -0.019 |
| ATRRH | right | FA | 0.017 |
| posterior ramus of the lateral sulcus | right | CTh | -0.016 |
| pallidum | right | SCVol | 0.015 |
| S middle occipital and Lunatus | left | CTh | -0.015 |
| G temporal inferior | left | SDep | -0.014 |
| G temporal middle | right | SDep | 0.014 |
| UNCRH | right | FA | 0.009 |
| FXCUTLH | left | MD | 0.008 |
| G and S frontomargin | right | CTh | -0.007 |

*Abbreviations: MD = mean diffusivity, FA = fractional anisotropy, CTh = cortical thickness, CAr = cortical area, SDep = sulcal depth, CVol = cortical volume, SCVol = subcortical volume; G = gyrus, S = sulcus, +ve beta weights represent a positive association of E2 with the specific regional feature while –ve beta weights represent a negative association of E2.*

#### 3.3.1 E2

Of the 37 features selected in the unimodality models, 31 were found to explain variance in E2 in the multimodal model (R = 0.03, r = 0.24). E2 showed associations with temporal (middle, superior, and inferior), occipital (lateral, anterior, polar), superior frontal, insula, and orbitofrontal gray matter structure (Table 5; Figure 4). The highest weighted associated cortical features were reduced sulcal depth in the left paracentral region, reduced cortical thickness in left frontomarginal (lateral orbitofrontal) and right insula regions, and greater cortical thickness in the left occipital lobe and left temporoparietal junction. Additionally, higher E2 levels were associated with WM tracts connecting frontal lobe with other regions. The highest weighted associated WM features were increased MD in the white matter connecting bilateral prefrontal regions (FMIN) and increased FA in the white matter connecting the superior cortex and striatum in the left hemisphere.

**Figure 3:**
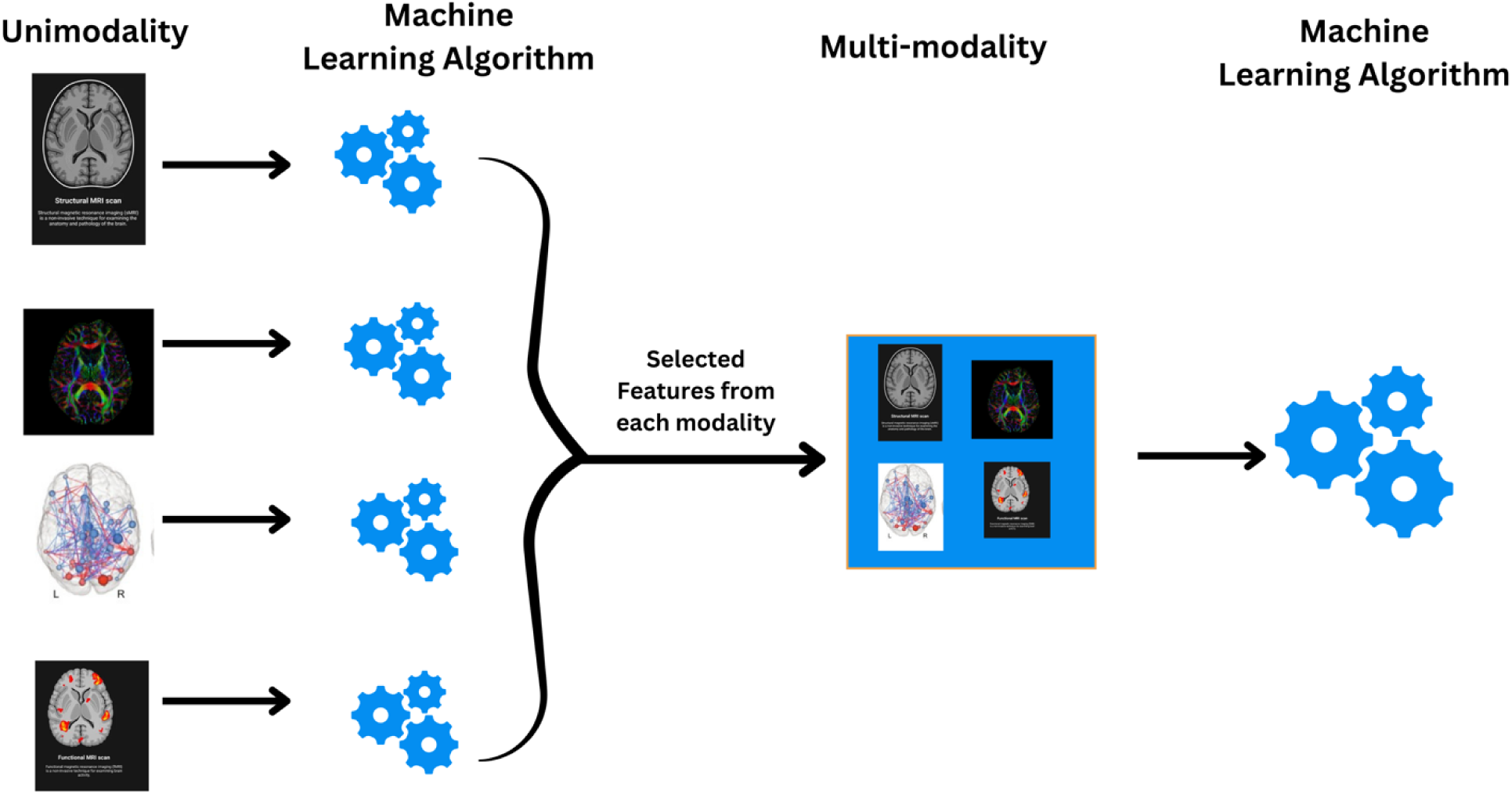
Summary of the analysis pipeline - applying machine algorithm on the features from each of the unimodality and then taking all the selected features into a single multi-modality model. This process was repeated for each hormone. [Blue cogs show machine learning algorithm in Figure 2]

**Figure 4:**
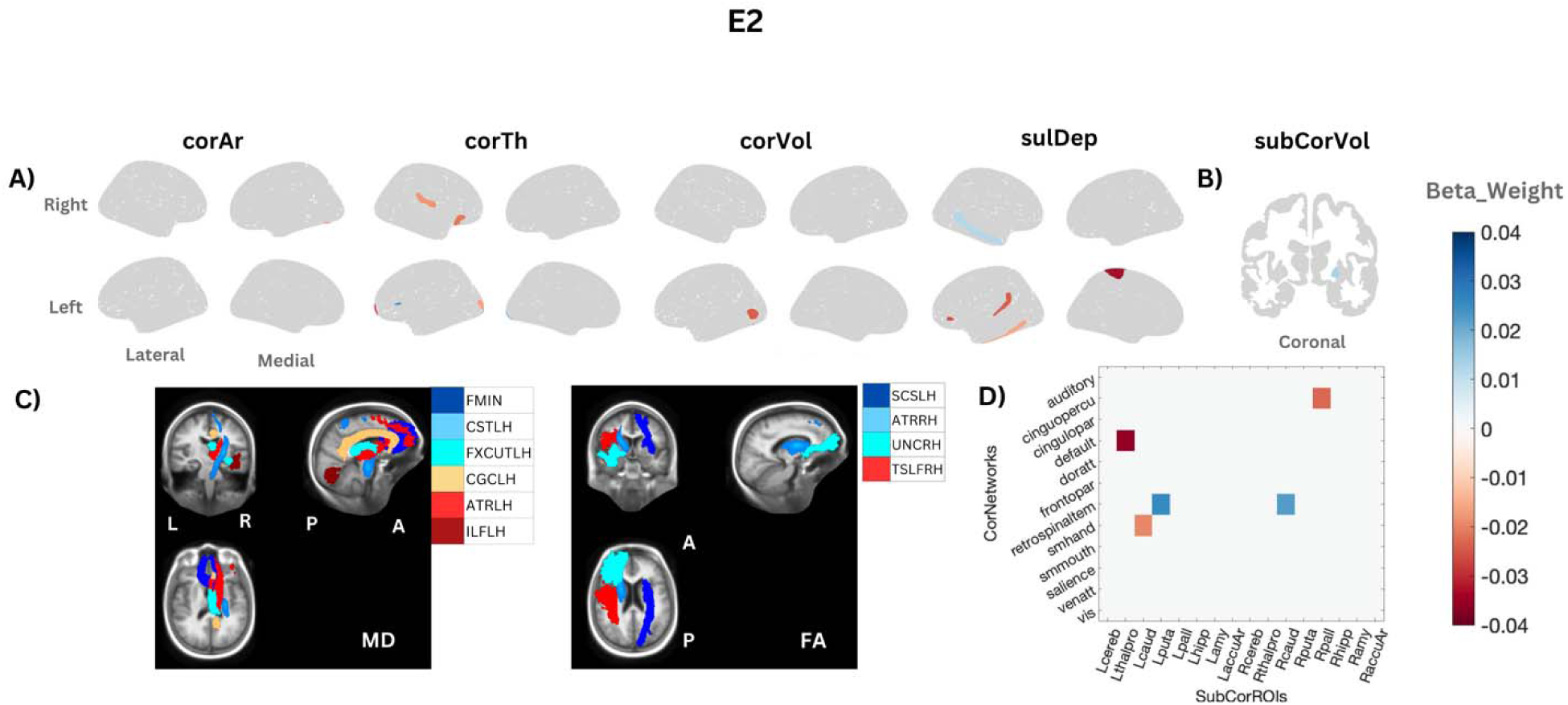
Association of E2 with different brain regions across different modalities in the multimodal analysis. The colour bar represents the beta weight of the fitted model; from blue (positive) to red (negative). Panel A) E2’s association with different structural features. Panel B) association with subcortical volume in pallidum. Panel C) associations with MD and FA within different tracts; legends are arranged from positive highest weight to negative highest weight such that positive weights are shown by blue shades, and negative weights are shown by red shades). Panel D) association with resting-state connectivity matrix. Abbreviations: corAr: cortical area, corTh: cortical thickness, corVol: cortical volume, sulDep: sulcal depth, subCorVol: subcortical volume, L & LH: Left, R & RH: Right, A: Anterior, P: Posterior; FMIN: forceps minor, CST: corticospinal tracts, FXCUT: Fornix cut; CGC: cingulum cingulate, ATR: anterior thalamic radiations, ILF: inferior longitudinal fasciculus, SCS: superior corticostriate, UNC: uncinate fasciculus, TSLF: temporal superior longitudinal fasciculus.

Functionally, E2 was associated with resting-state connectivity between cortical networks and subcortical regions only. Among all, the highest weighted functional features were associations with reduced resting-state connectivity between default mode network and left thalamus as well as greater resting-state connectivity between the retrosplenial network and left putamen.

#### 3.3.2 Tes

Multimodal elastic-net regression revealed that 71 out of 74 features explained variance in Tes (R2 = 0.14, r = 0.41). Tes showed diverse associations with the structure of the cingulate, frontal, temporal, and occipital cortices (Table 6; Figure 5). In terms of the highest weighted features, there was a positive association with cortical thickness of the left occipital lobe and thickness of left precentral and volume of the left hippocampus. Tes showed a strong negative association with thickness in the right calcarine, left subparietal, and right posterior cingulate, and volume of the left precuneus. Additionally, Tes showed associations with WM across the whole brain. The strongly associated WM features were higher MD in FMIN and IFOLH, and higher FA in left SCS, right UNC, left SIFCLH, and FMAJ, while lower MD in the CC, PSLFLH, and FSCFLH, and with lower FA in the PSCSRH and CC.

**Figure 5:**
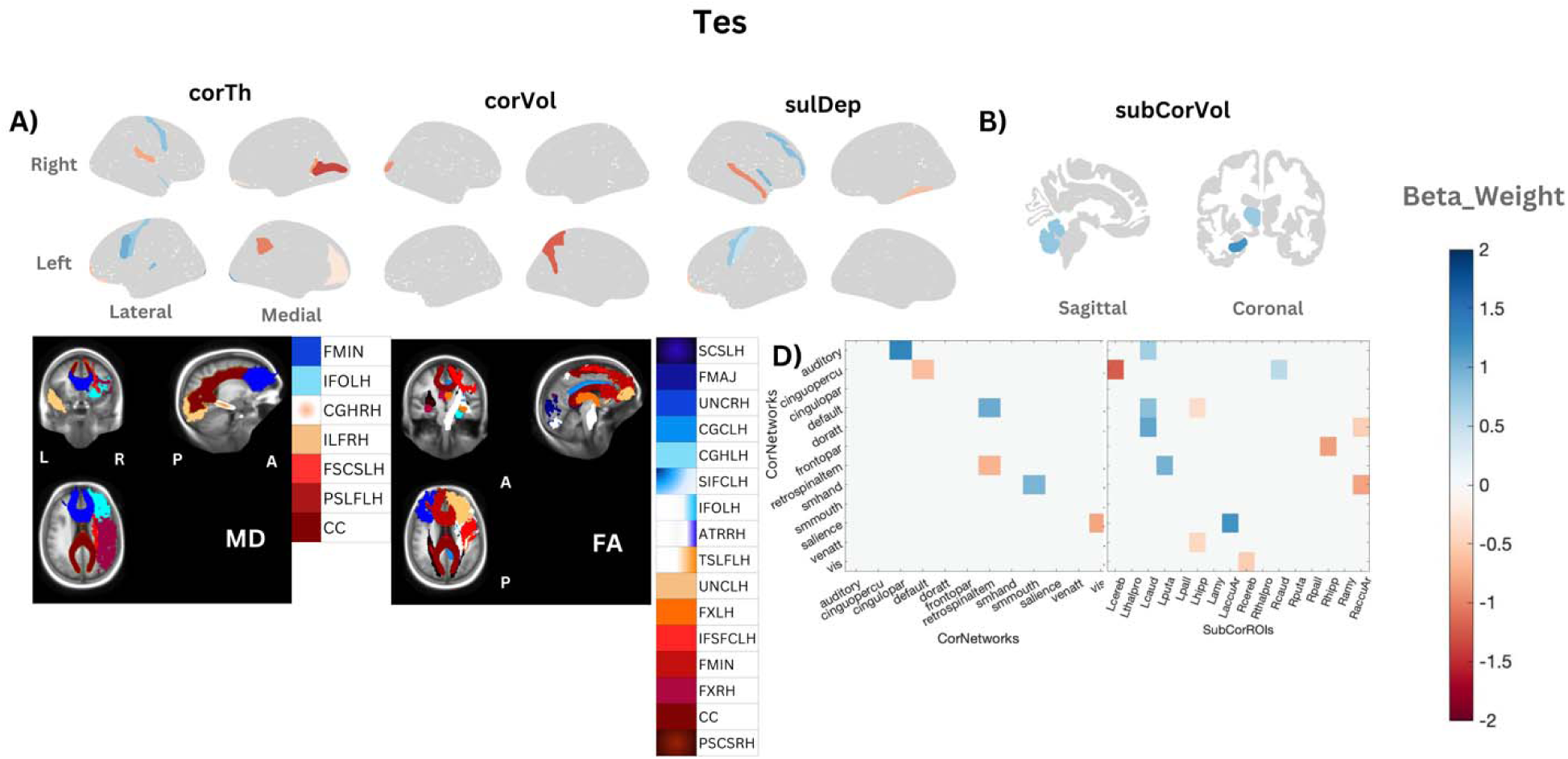
Association of Tes with different brain regions across different modalities in a multimodal analyses, the colour bar represents the beta weight of the fitted model; from blue (positive) to red (negative). Panel A) Tes’s association with different cortical structural features and with subcortical volume (Panel B)). Panel C) associations with MD and FA within different tracts (legends are arranged from positive highest weight to negative highest weight such that positive weights are shown by blue shades and negative weights are shown by red shades). Panel D) associations with resting-state connectivity matrix. Abbreviations: corAr: cortical area, corTh: cortical thickness, corVol: cortical volume, sulDep: sulcal depth, subCorVol: subcortical volume, L & LH: Left, R & RH: Right, A: Anterior, P: Posterior; FMIN: forceps minor, IFO: inferior fronto-occipital fasciculus, CGH: cingulum (parahippocampal), ILF: inferior lomgitudinal fasciculus, PSCS: parietal superior corticostriate, FSCS: frontal superior cortico striate, CC: corpus callosum, SCS: superior corticostriate, FMAJ: forceps major, UNC: uncinate fasciculus, SIF: striatal inferior frontal cortex, TSLF: temporal superior longitudinal fasciculus, FX: Fornix; CGC: cingulum cingulate, ATR: anterior thalamic radiations , IFSFC: inferior frontal superior frontal cortex.

**Table 6:**
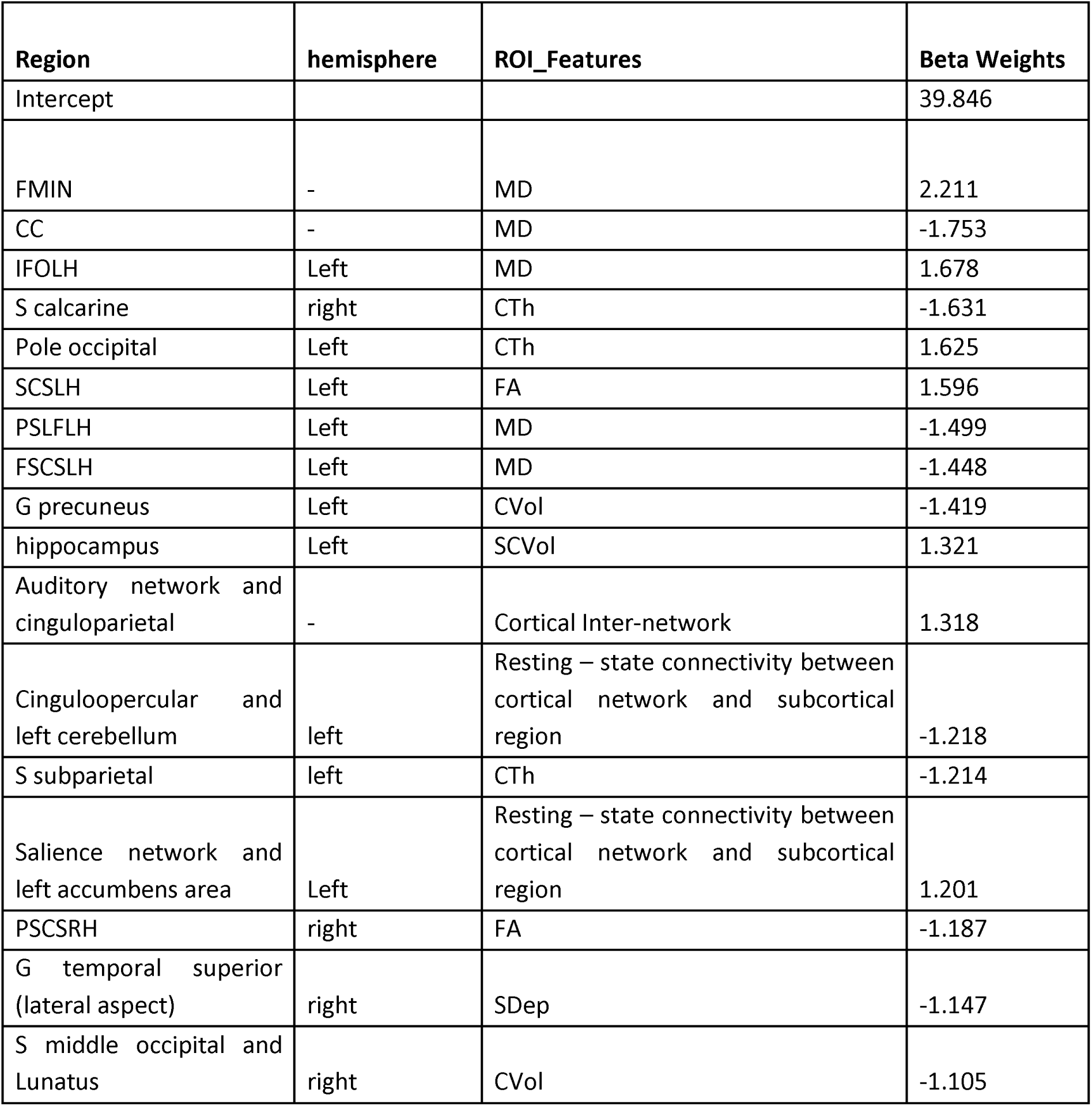
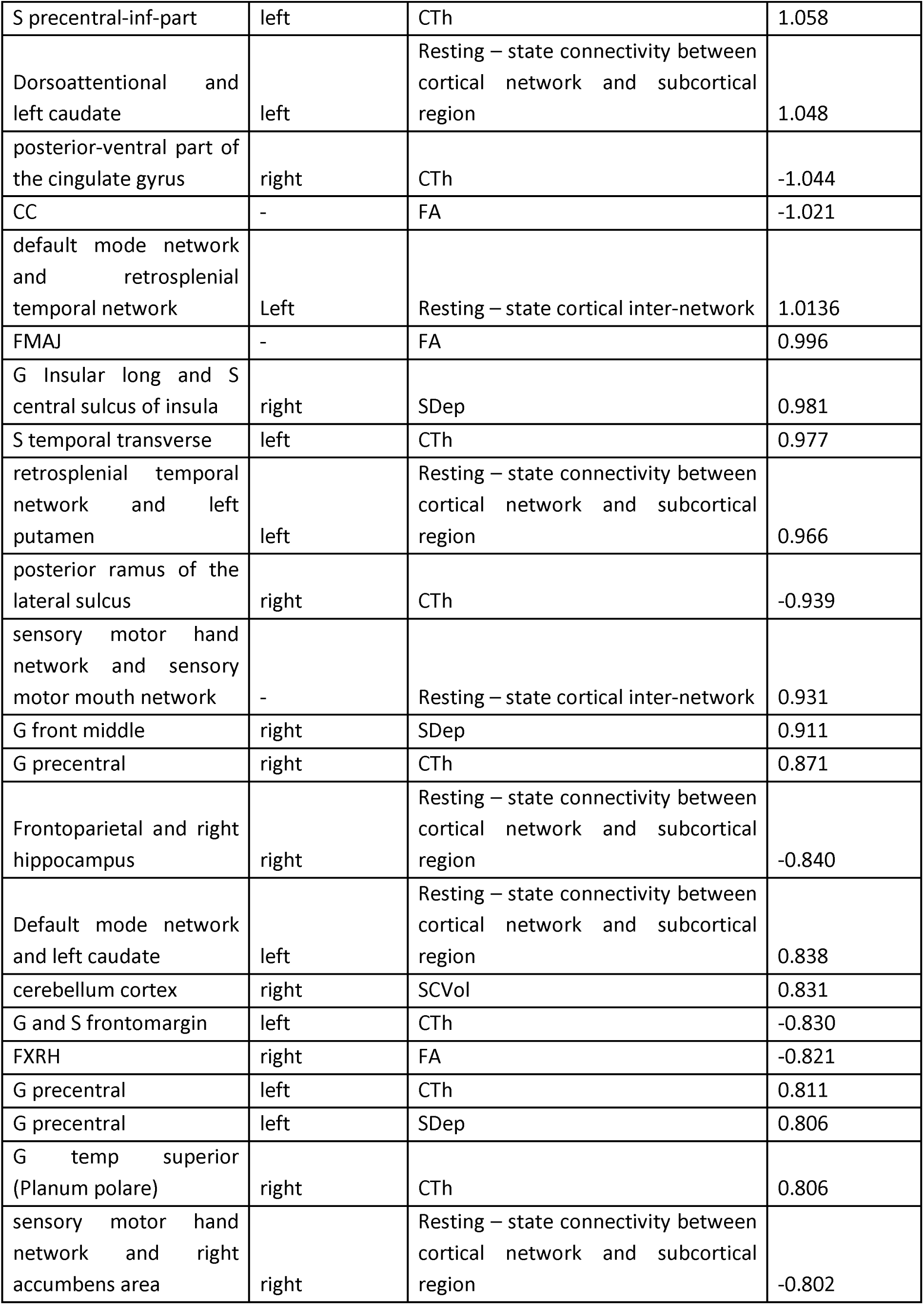
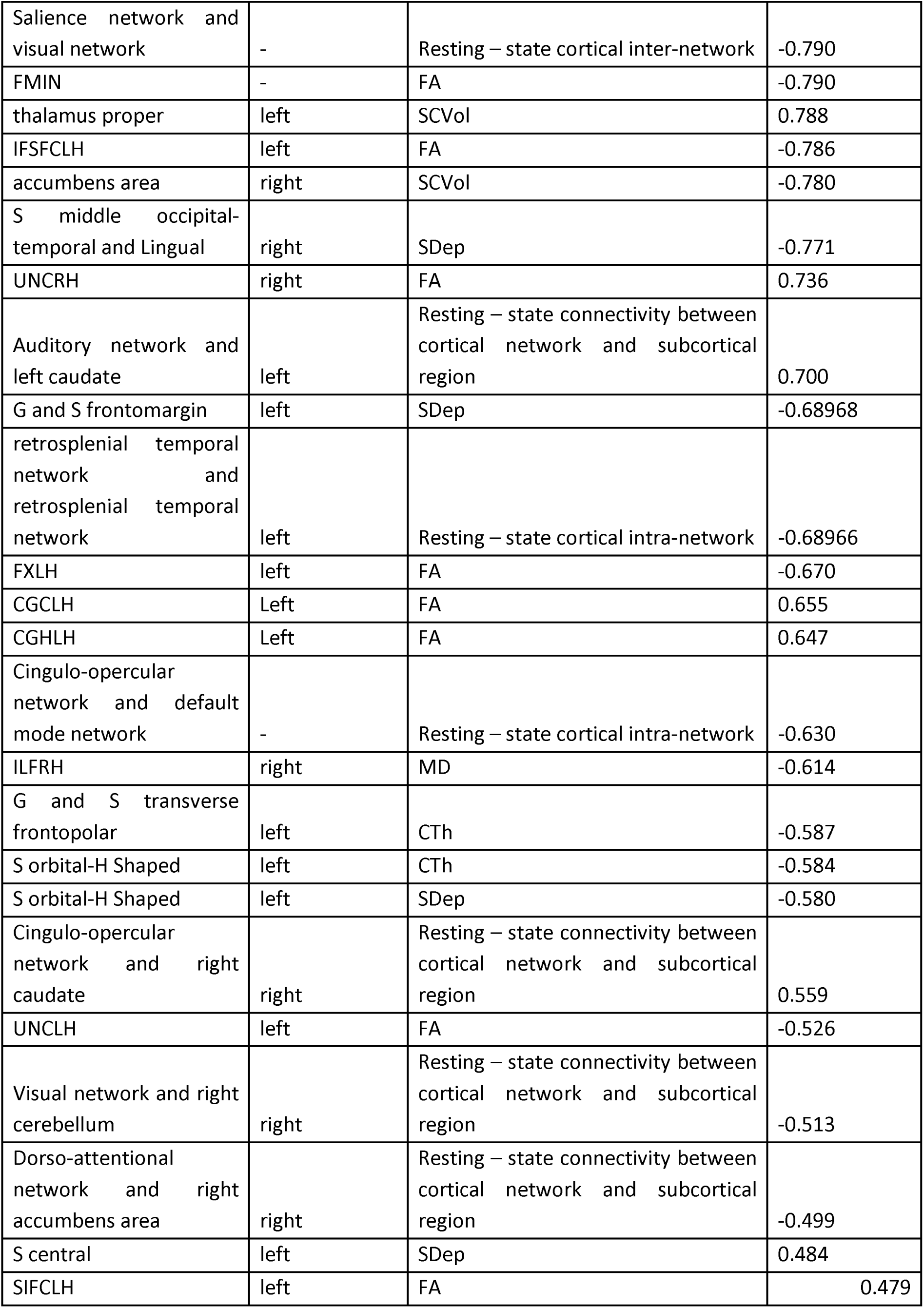
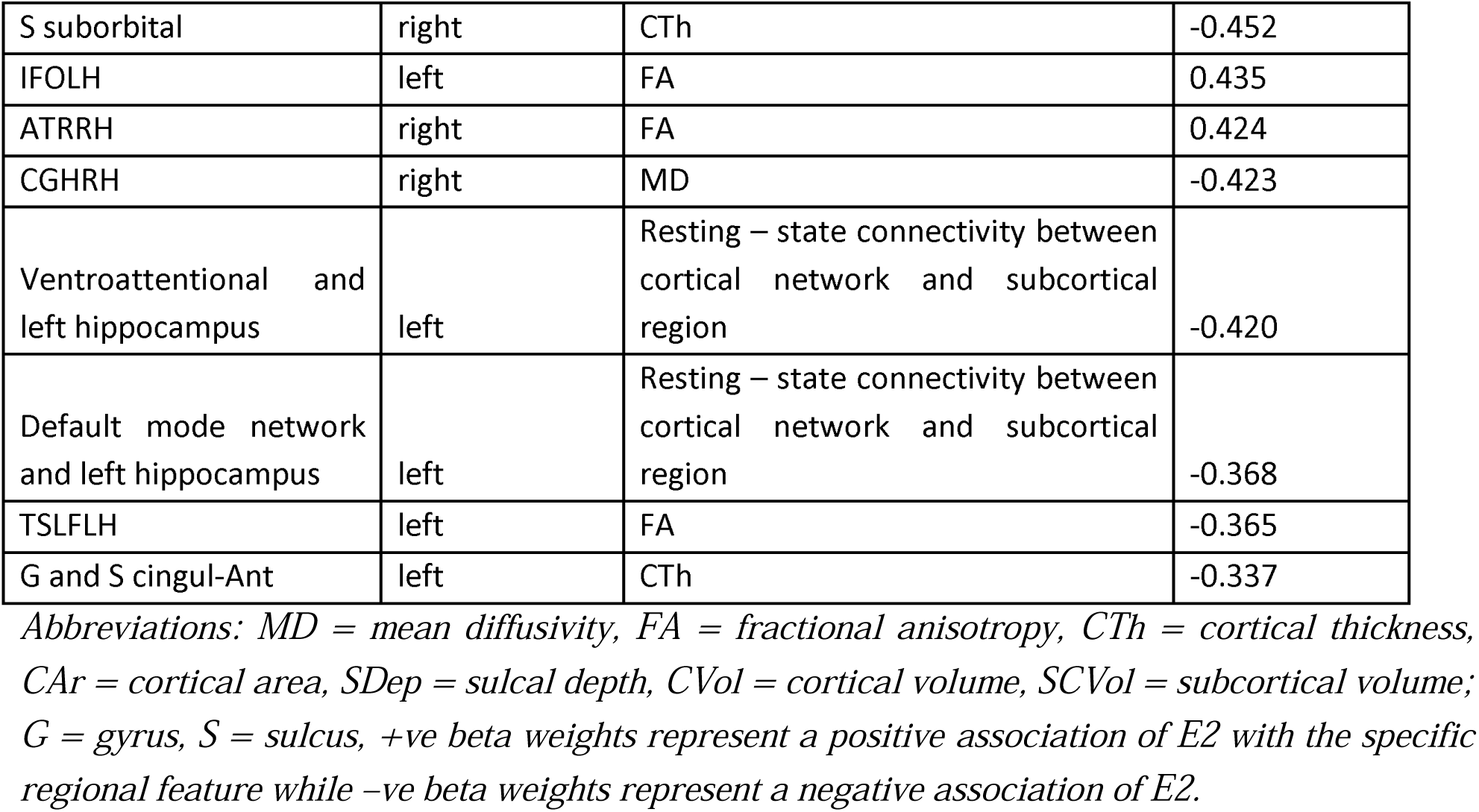
Selected Features from multimodal ENet regression for Tes

Functionally, task-based activity was not associated with Tes levels. Interestingly, associations with resting state connectivity were in diverse networks; the majority of the associations were within cortical-subcortical networks. Tes showed the strongest positive relationship with different inter-cortical network connections (between auditory and cingulo parietal networks; between default mode and retrosplenial networks; and between somato motor hand and somato motor mouth networks), and between cortical networks and subcortical regions (such as left caudate with dorso-attentional networks, and left accumbens area with the salience network), while there were strong negative associations with connectivity between cortical networks and subcortical regions (cingulo-opercular with left cerebellum).

#### 3.3.3 DHEA

Multimodal elastic-net regression revealed that 62 out of 79 features explained variance in DHEA levels (R2 = 0.07, r = 0.32) (Table 7; Figure 6). DHEA showed diverse associations with occipital, frontal, temporal and parietal cortices. In terms of the highest weighted features, structurally, DHEA showed positive associations with cortical thickness in the left occipital lobe, left precentral region, sulcal depth in left middle frontal and right insula, and volume of the right pallidum and left thalamus proper. Strong negative associations were found with volume in left precuneus, right superior frontal, and right lateral occipital, surface area in left inferior occipital, cortical thickness in left lateral orbital, right calcarine, and sulcal depth in right superior temporal. Additionally, DHEA showed wider associations with WM tracts across the whole brain. The highest positive association between DHEA and MD in IFOLH and FMIN, and FA in CGHLH, SCSLH, ILFRH, and FMAJ. However, the strongest negative association between DHEA and FA in PSCSRH and FXLH. (Table 5)

**Figure 6:**
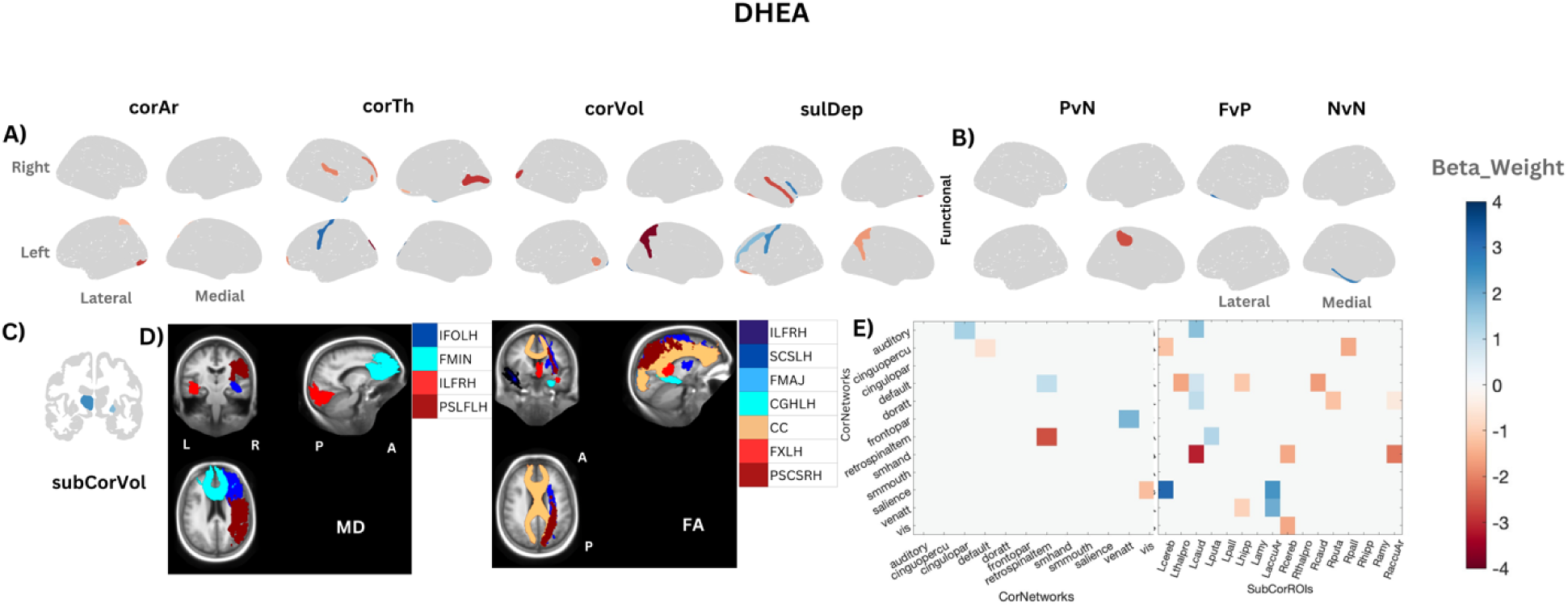
Association of DHEA with different brain regions across different modalities in multimodal analyses, the colour bar represents the beta weight of the fitted model; from blue (positive) to red (negative). Panel A) DHEA’s association with different cortical structural features. Panel B) associations with the activated functional regions in response to positive face vs neutral face, face vs place, and negative face vs neutral face. Panel C) association with subcortical volume in pallidum and thalamus proper. Panel D) associations with white matter microstructure; MD and FA (legends are arranged from positive highest weight to negative highest weight such that positive weights are shown by blue shades, and negative weights are shown by red shades). Panel E) associations with resting-state connectivity matrix. Abbreviations: corAr: cortical area, corTh: cortical thickness, corVol: cortical volume, sulDep: sulcal depth, subCorVol: subcortical volume, PvN: positive vs neutral faces, FvP: face vs places, NvN: negative vs neutral faces; L & LH: Left, R & RH: Right, A: Anterior, P: Posterior; FMIN: forceps minor, IFO: inferior fronto-occipital fasciculus, ILF: inferior longitudinal fasciculus, PSLF: parietal superior longitudinal fasciculus, SCS: superior cortico striate, FMAJ: forceps major, CGH: cingulum (parahippocampal), CC: corpus callosum, FX: fornix, PSCS: parietal superior corticostraiate

**Table 7:**
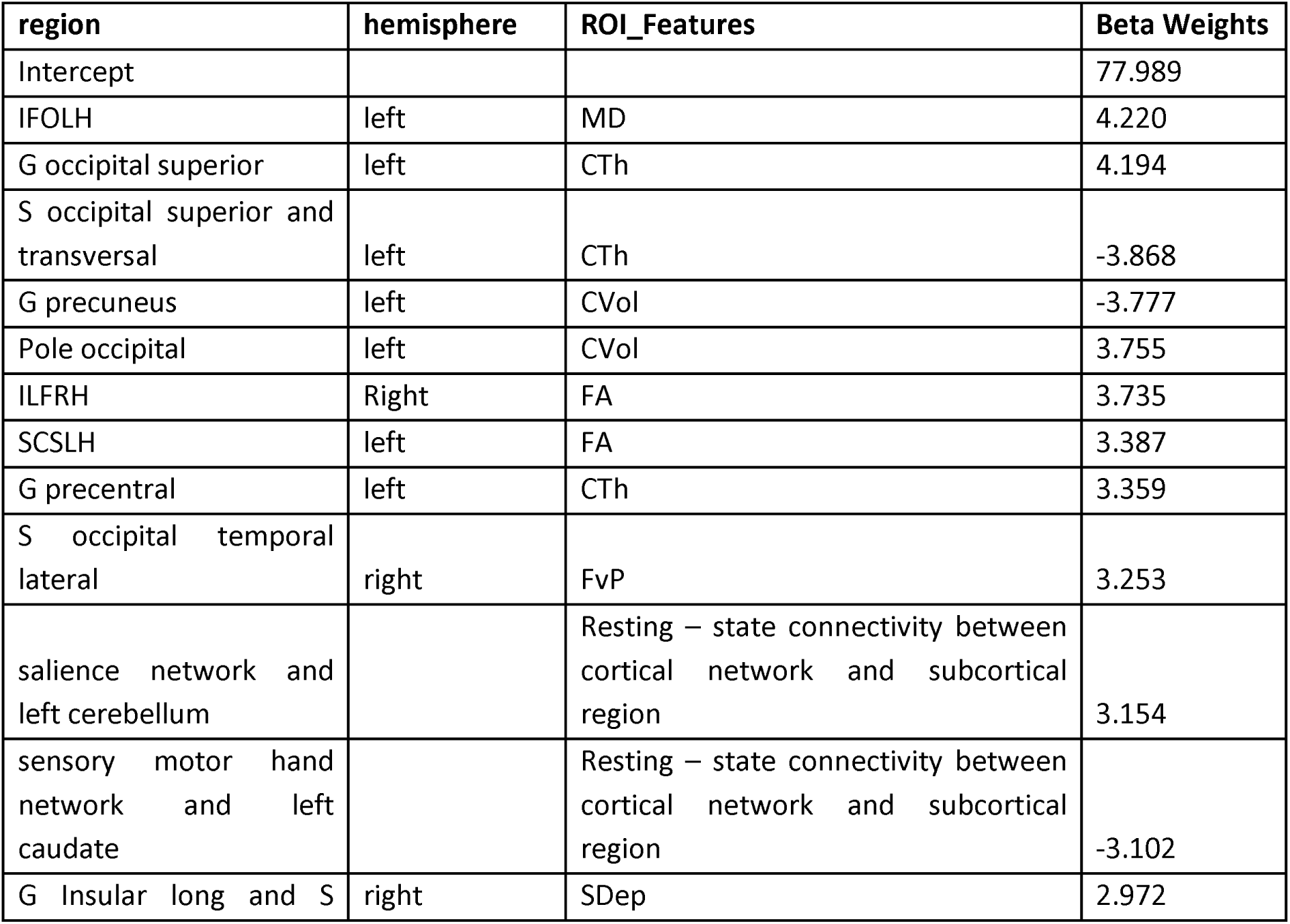
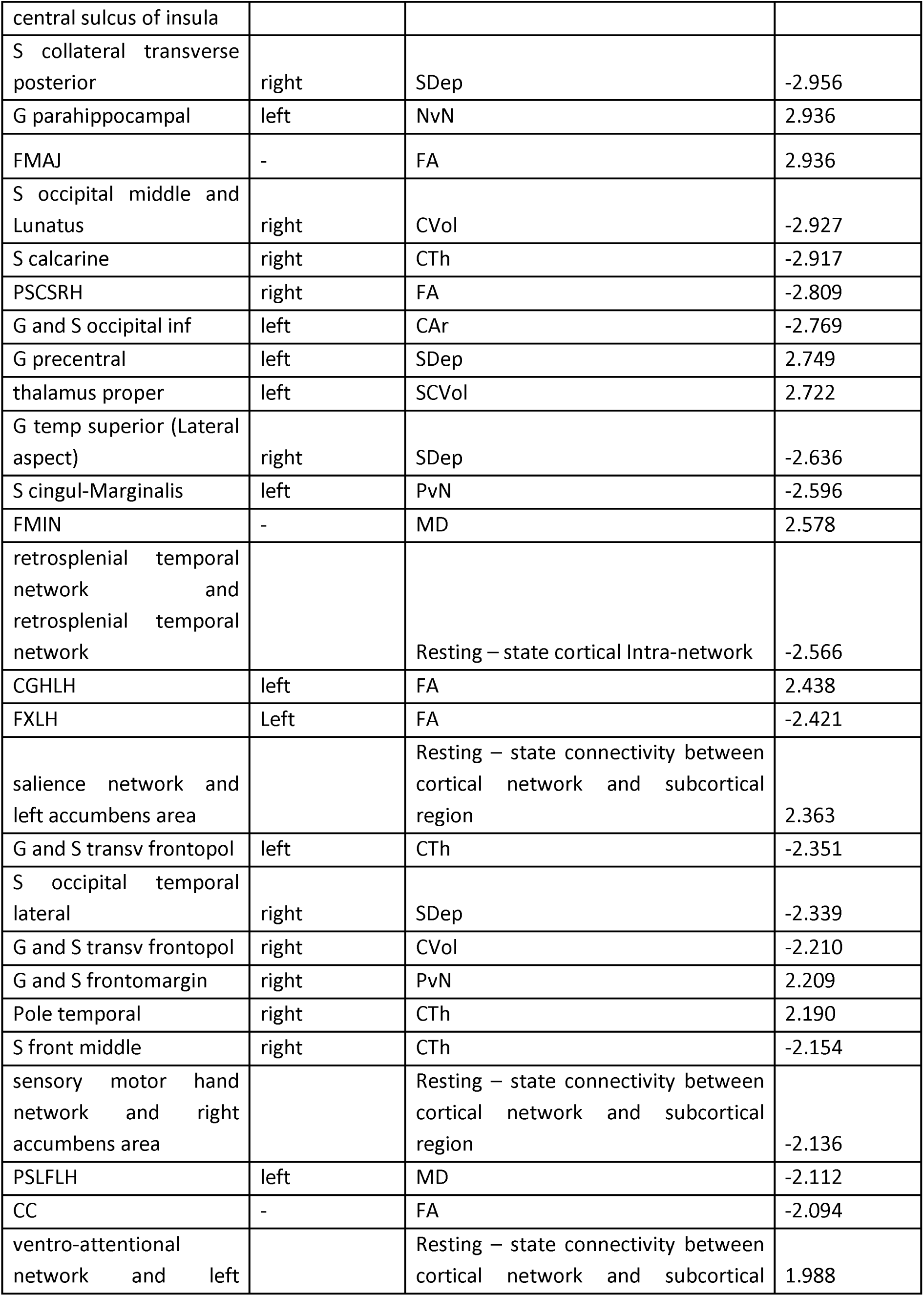
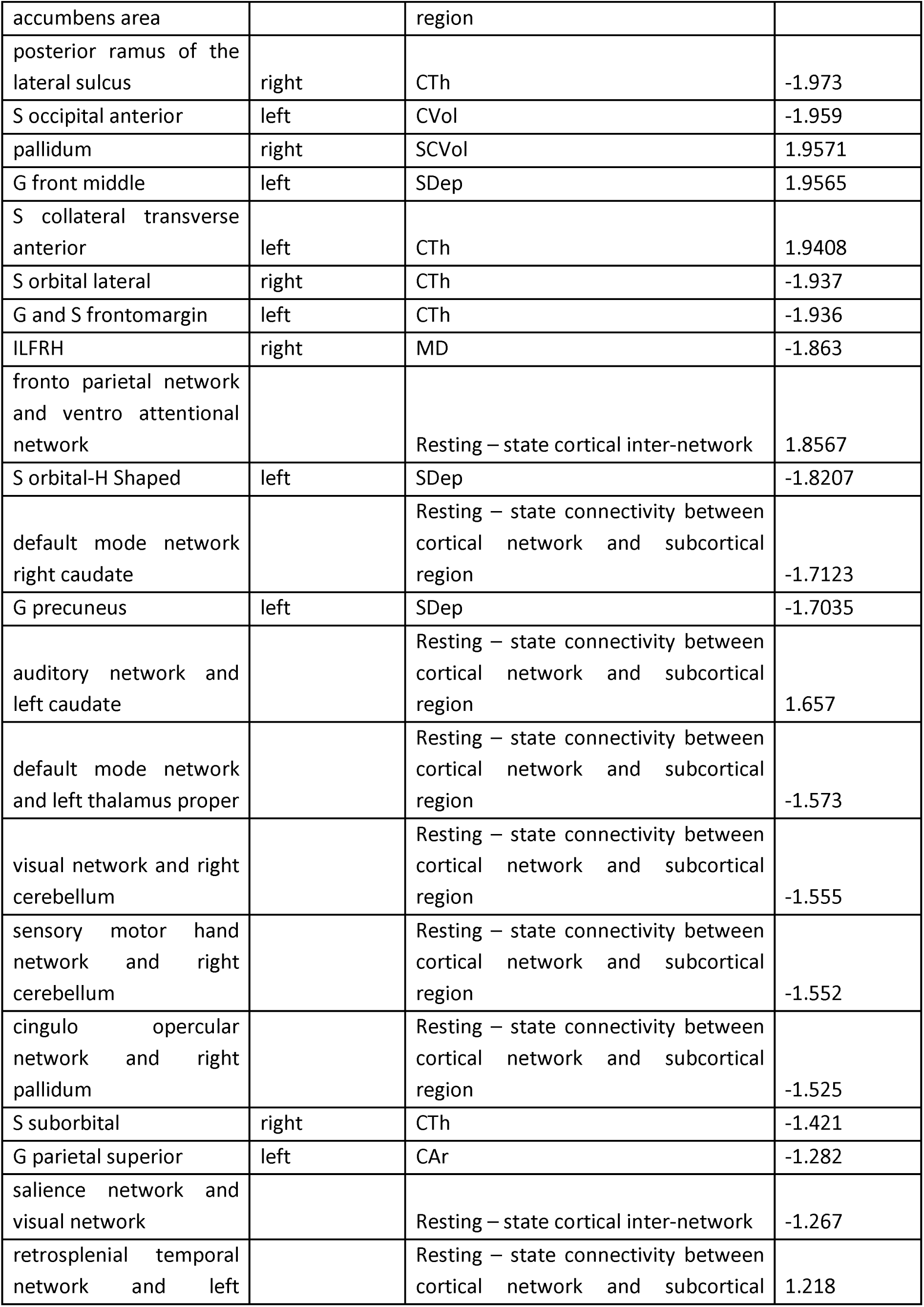
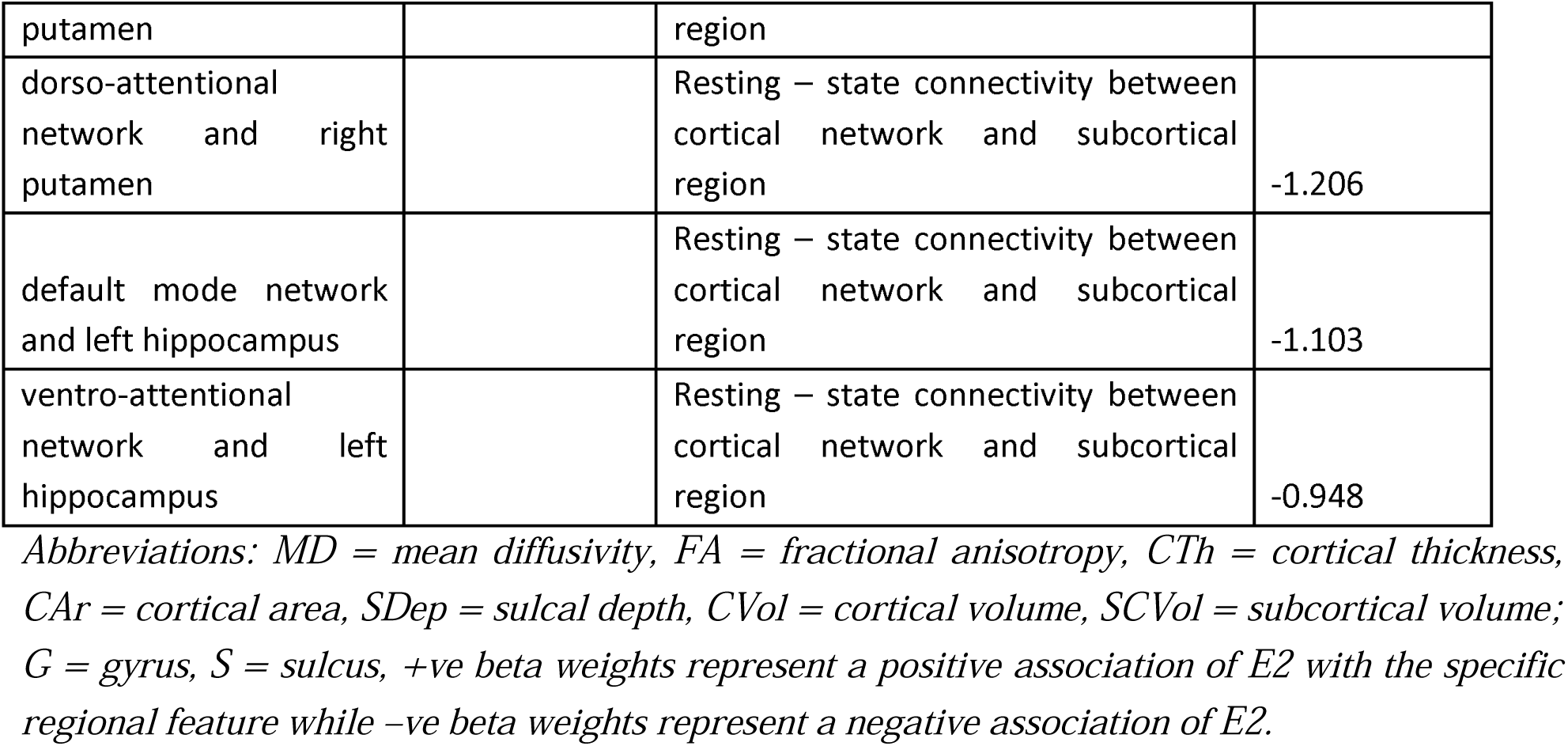
Selected Features from multimodal ENet regression for DHEA

Functionally, DHEA showed associations with both task-based activity and resting state connectivity. The highest weighted functional features were increased activity in left parahippocampal in response to negative emotion stimuli (NvN), and with right lateral occipital temporal activity in response to face stimuli (FvP), and a decreased activity in left cingulate in response to positive face stimuli (PvN). During rest, DHEA was strongly associated with greater connectivity between cortical networks and subcortical regions (salience network with left accumbens area and left cerebellum), and reduced intra retrosplenial network connectivity and subcortical to cortical network connectivity (between the somato motor hand network and left caudate).

### 3.4 Sensitivity Analyses

There were some changes in the findings after controlling age, such as all three hormones were no longer associated with white matter microstructures and task-based fMRI features. However, the strongest features were still implicated for all the hormones, although with different effect sizes. Results from the unimodality and multimodality of age-regressed models are presented in Supplementary Section 3A - 3.1.2. & 3.2.2, respectively. In addition, accounting for caffeine intake and collection time did not have a large impact on findings. Almost 90% of features survived after removing variance associated with caffeine intake (Refer Supplementary Section 3C and 3D). Interestingly, after removing hormone outliers, the strongest features were still in the brain regions and with the same direction of effects (Refer Supplementary Section 3 B).

## 4. DISCUSSION

In this study, we explored the association of steroid hormones with brain structure and function in a large, demographically diverse group of 9-13-year-old females from the ABCD Study, using a multimodality approach with cross-validation. Although brain features explained a small proportion of variance in hormones, our approach ensured findings were robust. Across brain imaging modalities, E2 levels showed the strongest associations with morphological features (including GM structure and WM microstructures) in the prefrontal and motor cortices. Tes and DHEA showed strongest associations with the structure and connectivity of limbic (e.g., hippocampus, striatum), occipital and parietal regions. In addition, all three hormones were associated with some overlapping structural and functional features, although these effects were weaker than the unique (i.e., non-overlapping) associations identified for each hormone. Of note, DHEA was the only hormone that showed associations with task-based brain function.

### 4.1 Structural and functional brain features associated with E2

Among all the implicated brain regions, E2 levels showed the strongest associations with the motor cortex and prefrontal cortex, whereby higher E2 levels were associated with reduced sulcal depth in the left paracentral and thickness in frontal pole, and a positive association with MD in tracts connecting prefrontal regions (forceps minor) and FA in tracts connecting superior cortex and striatum (left SCS) and motor cortex and spinal cord (left CST). E2 also had a strong negative association with resting state connectivity between the default mode network and the left thalamus. Prior studies of adolescent females have similarly found E2 to be associated with alterations in the prefrontal (Peper et al., 2009; Brouwer et al, 2015) and primary motor cortical gray matter (Stoica et al., 2019), although implicating different structural metrics. Further, animal studies (Lenz et al., 2010; Campbell et al., 2014; Arevalo et al., 2015) have shown that E2 plays a role in synaptogenesis and myelination in frontal areas. The prefrontal and somatosensory cortices have been implicated in emotion regulation and other executive functions, such as working memory, in adolescents (Vijayakumar et al., 2014) (Kropf et al., 2018; Satterthwaite et al., 2013). Higher FA in fronto-striatal tracts (Darki and Klingberg, 2015) and tracts connecting the bilateral prefrontal cortex have also been associated with working memory among adolescents (see review by Ribeiro et al., 2024). As such, our findings suggest a neurobiological mechanism by which E2 influences certain emotion and memory processes (Chung et al., 2019; Hampson, 2018; Toffoletto et al., 2014). Furthermore, recent work (Harrison et al., 22) has suggested that DMN-thalamus connectivity may mediate DMN-controlled behaviors, such as social cognition and memory (Menon et al., 2023), via an excitatory influence of the thalamus on DMN activity. Therefore, our findings of reduced DMN-thalamus connectivity with increased E2 levels might support a role for E2 in alterations in DMN-related social cognitive and memory functions.

Interestingly, most of the brain regions and white matter pathways most strongly associated with E2 were in the left hemisphere, which is in line with prior findings in adults (e.g., (Weis et al., 2008; Bibawi et al., 1995)). Left lateralisation of function among women during the high E2 phase of the menstrual cycle is a consistent finding in adult studies (Rode et al., 1995; Chiarello et al., 1989), including those investigating cognitive and memory functions (Volle et al., 2008; Fletcher et al., 1998; Frith et al., 1996), and it has been suggested that this reflects a role of E2 in brain lateralisation. Thus, our findings may further suggest a role of E2 levels in the lateralisation of these functions during adolescence.

### 4.2 Structural and functional brain features associated with androgens (Tes and DHEA)

There was a wide overlap in the structural correlates of Tes and DHEA, which might be due to the fact that they are both androgens, highly correlated (r = 0.77) and act on the same ARs in the brain (Sisk et al., 2005; Nunez et al., 2003). The strongest associations for both Tes and DHEA were with the structure of occipital and parietal regions. Higher hormone levels were associated with reduced thickness and volume in occipital-parietal areas, including the calcarine sulcus and precuneus, aligning with previous research (Vijayakumar et al., 2019; Bramen et al., 2012; Nguyen et al., 2013a; Neufang et al., 2009). In addition, both of these hormones were positively associated with precentral thickness partially supporting Nguyen et al.’s (2013) findings. Of subcortical regions, only higher Tes levels were strongly associated with higher hippocampus volume, consistent with prior work (Vijayakumar et al., 2021; Wierenga et al., 2018). All these regions (occipital, parietal, precentral, and hippocampus) have been implicated in visuospatial functions (Clements-Stephens et al., 2009; Whittingstall et al., 2014; Casey et al., 2005; Gilchrist et al., 2018). For instance, Nguyen and colleagues (2016) found that higher DHEA levels were associated with higher visual awareness, mediated by occipital-limbic structural covariance. Adult studies have also shown that higher androgen levels were associated with visuospatial memory, cognitive performance, and verbal fluency in pre- and post-menopausal women (Hausmann et al., 2000; Ryan et al., 2012; Stangl et al., 2011), and in women with polycystic ovary syndrome (Barry et al., 2013). Therefore, it can be speculated that androgens (Tes and DHEA) may exert influences on these cognitive functions in female adolescents via effects on these brain structures.

Additionally, associations between androgen levels and WM microstructure may reflect their role in cognitive functioning. For example, both Tes and DHEA showed positive associations with MD in the IFO (tract connecting frontal and occipital regions), while Tes alone showed a negative association with MD in the corpus callosum (CC). Alterations in WM microstructure within these tracts have been implicated in visuospatial working memory and across a wide range of cognitive functions (Krogsrud et al., 2018; review by Goddings et al., 2021). Previous studies reported similar findings, with higher DHEA and Tes associated with reduced WM connectivity in the IFO (Barendse et al., 2018, 2020) and lower frontal WM volume (Klauser et al., 2015), though these focused on late childhood. Mixed results were found in early- to mid-adolescents, with some studies showing no Tes associations with FA in IFO (Ho et al., 2020) and others finding associations in males but not females (Herting et al., 2012). However, Ho et al. (2020) reported that higher Tes correlated with increased FA in the CC, aligning with our findings. These results suggest that androgen-related WM changes and their cognitive effects may vary by developmental stage, requiring further investigation.

Interestingly, Tes and DHEA both showed similar and different associations with striatal structural and functional connectivity. Both hormones were positively associated with FA in the left SCS (tract connecting superior cortex and striatum) while only Tes was negatively associated with FA in the left FSCS (tract connecting parietal cortex and striatum). Tes was also positively associated with resting-state connectivity between the salience network and nucleus accumbens, while DHEA was negatively associated with connectivity between somato-motor hand network and caudate. These findings partially align with prior work relating Tes with striatal connectivity in adolescent females (Ladouceur et al., 2019), suggesting a role in reward processing (Somerville et al., 2011). In contrast, DHEA’s association with parietal-striatal connectivity may relate more to motor functions (Rodriguez-Sabate et al., 2016; Fiore et al., 2003) rather than reward processing (Ladouceur et al., 2019). Studies have also found antagonistic and interactive effects of Tes and DHEA on the brain (Nguyen et al., 2018; Barendse et al., 2018), suggesting these hormones may influence the brain through different androgen pathways. Our findings support the idea that these hormones may contribute to shaping the neural mechanisms underlying both similar and different domains of behavioural functioning.

Of note, DHEA was the only hormone associated with task-based functional activity, aligns with previous research showing DHEA’s involvement in affective processing (Whittle et al., 2015). Specifically, DHEA levels were positively associated with activity in the lateral occipital-temporal region (fusiform gyrus) in response to face vs. place stimuli and negative vs. neutral faces, suggesting a role in facial affect processing. In contrast to our null findings, some prior research has identified associations between E2 or Tes and affective processing (Cservenka et al., 2015; Chung et al., 2019; Vijayakumar er al., 2019). One explanation for our lack of findings could be that E2 and Tes are not associated with the specific affective processes captured by the task utilised in this study. Alternatively, null findings may be driven by certain study limitations (discussed below).

### 4.3 Overlapping findings for all hormones (E2, Tes, DHEA)

Although not necessarily among the strongest contributing features, of note was that all three hormones showed associations with the structure of the right temporoparietal junction/SMG, right insula, and bilateral orbitofrontal and frontopolar regions. There was also a positive association of all hormones with MD in the forceps minor (connecting bilateral prefrontal regions). These regions have widely been implicated in cognitive and social functions (such as attention, theory of mind), and emotion processing, experience and regulation (Doricchi et al., 2022; LaVarco et al., 2022; Rolls et al., 2020) (Nagai et al., 2007). Previous studies also have identified associations between specific hormone levels and the structure of these regions (DHEA: Nguyen et al., 2013; Tes: Bramen et al., 2012; Koolschijn et al., 2014; Nguyen et al 2016; E2: Brouwer et al., 2015). Hormone levels have also been associated with the function of the insula and OFC, specifically in adolescent females (Whittle et al., 2015; Op de Macks et al., 2016; Op de Macks et al., 2017). Although the implications of our findings for brain function are unclear, together with prior work, they suggest a common role for all three hormones in shaping the development of regions important for a range of adolescent behaviours.

### 4.4 Limitations and future directions

There are some limitations of the current study that should be noted. Firstly, our study design is cross-sectional. While the ABCD study has longitudinal data, we intentionally examined cross-sectional associations given the lack of prior research investigating hormonal (particularly E2) associations with brain structure and function in adolescent females. However, it would be of interest for future research to track associations between hormone levels and brain structure and function over time. Another important area of future research is to explore the unique and interactive influence of these hormones on the brain, to understand the likely complexity of hormone-brain associations.

In addition, we did not account for within-person variability in hormone levels (particularly relevant for E2 levels, which fluctuate across time, particularly in females who have reached menarche [∼300 in our sample]). The ABCD study lacks comprehensive information regarding cycle length and phase of the cycle the saliva sample was collected. Further, samples were collected at different times of the day, which introduces variability given the hormones investigated have diurnal variation (Matchock et al., 2007; Mitamura et al., 2000). Although we ran some sensitivity analysis indicated that time of day did not influence the strongest findings. Resulting variability in hormone levels may still affect our ability to detect effects.

Finally, we did not include participant age as a covariate in primary analyses, given that doing so would remove meaningful variance in hormone levels (due to the significant correlation between age and hormone levels). However, we ran supplementary multimodality models after regressing the age from each of the hormones. Some of the hormone effects remained (see Supplementary Materials), however, many (particularly involving WM) weakened. In addition, potentially confounding demographic variables were not included in any of the analysis models. Further research is needed to further dissociate the impact of hormones, age and other demographic variables on adolescent brain structure and function.

## 5. CONCLUSION

In conclusion, this study provides a comprehensive characterisation of the association between hormones and brain structure and function in adolescent females, which had not been explored previously in a sample of this size. Using a multimodal model with cross-validation provided us with robust findings while addressing the multidimensionality and multicollinearity issues with neuroimaging data. Our findings showed E2 to be associated with the structure and connectivity of brain regions involved in working memory and emotion processing, while Tes and DHEA were both associated with regions involved in a wide range of cognitive functions. In addition, all three hormones showed a majority of significant associations with resting-state functional connectivity between cortical and subcortical regions, highlighting their role in top-down/bottom-up regulation processes. Further studies are now needed to explore the behavioural implications of these hormone-brain associations.

### DECLARATIONS

## Acknowledgements

MK is supported by the Melbourne Research Scholarship at the University of Melbourne. The authors have declared that they have no competing or potential conflicts of interest and have agreed to the submitted version.

## Data Sharing

This study utilised data from the Adolescent Brain Cognitive Development (ABCD) Study (https://abcdstudy.org), available in the NIMH Data Archive (NDA). The ABCD Study is a large-scale, longitudinal project that began with roughly 11,000 children aged 9-10, tracking them over 10 years. Details on supporting organisations can be found at https://abcdstudy.org/federal-partners.html, and a full list of participating sites and study investigators is available at https://abcdstudy.org/scientists/workgroups/. Although ABCD consortium investigators were responsible for the study’s design, implementation, and data collection, they did not take part in the analysis or writing of this report. The views expressed in this manuscript are those of the authors and may not reflect those of the NIH or ABCD consortium investigators.

