## supplementary materials for "Pubertal hormones and the early adolescent female brain: a multimodality brain MRI study"

### 2.Methods

#### 2.1 Correlations among the hormones:

| **Hormone pairs** | **Correlation value** |
| --- | --- |
| E2-Tes | 0.55 |
| E2-DHEA | 0.56 |
| Tes-DHEA | 0.78 |

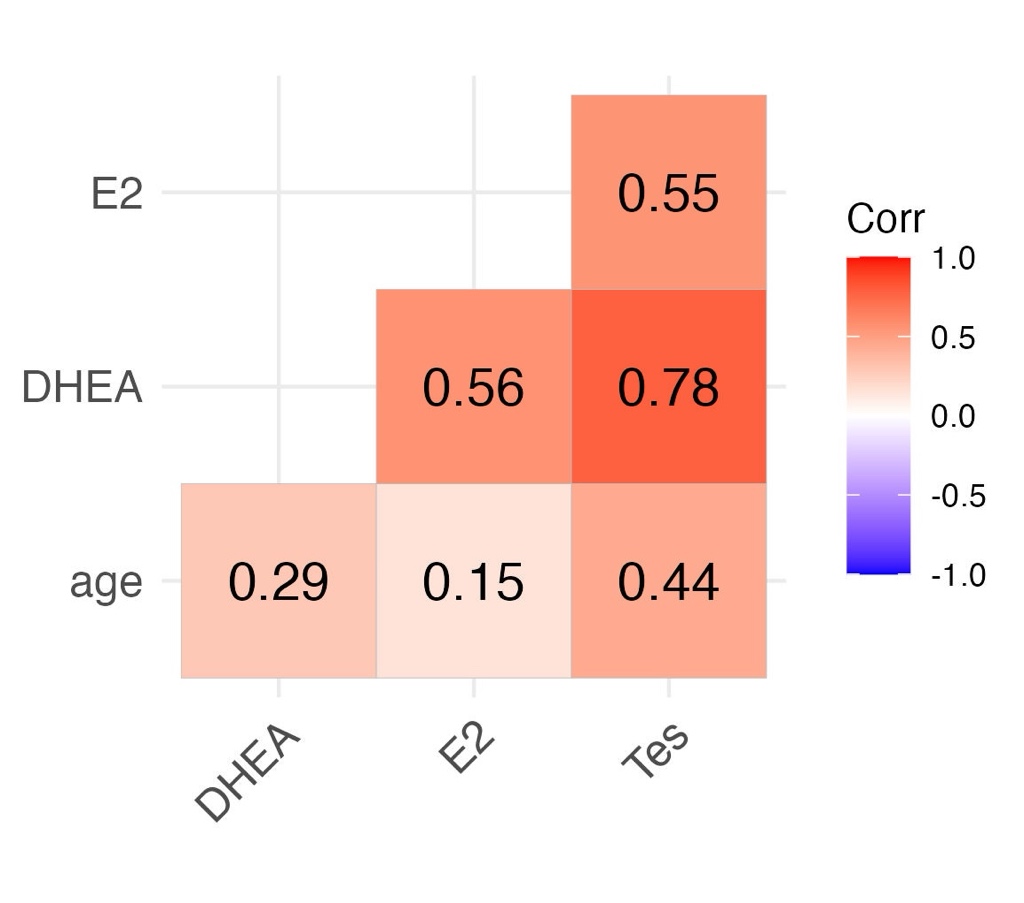

*Figure 1: Correlations of all three hormones and age with each other*

#### 2.2 Correlations of hormone levels with confounding factors at each wave

With raw hormone levels

|  | **N = 3025** | | | | | | |
| --- | --- | --- | --- | --- | --- | --- | --- |
|  | **E2** | | **Tes** | | | **DHEA** | |
| **Age** | 0.15 | p <0.00 | | 0.44 | p <0.00 | 0.29 | p <0.00 |
| **caffeine intake (yes/no)** | 0.129 | p<0.000 | | 0.1 | p<0.000 | 0.055 | p = 0.0021 |
| **physical activity (yes/no)** | 0.055 | p = 0.002 | | -0.002 | p = 0.907 | 0.0075 | p = 0.6789 |
| **Collection duration (in minutes)** | 0.024 | p = 0.18 | | -0.014 | p = 0.45 | 0.007 | p = 0.7 |
| **Collection time from midnight (in minutes)** | 0.136 | 4.579e-14 | | 0.00009 | 0.99 | -0.012 | 0.48 |
| **time from collection to freeze** | -0.011 | p = 0.54 | | -0.006 | p = 0.714 | 0.0097 | p = 0.592 |

Correlation of caffeine intake and physical activity with hormone levels after regressing age

|  | **N = 3025** | | | | | | |
| --- | --- | --- | --- | --- | --- | --- | --- |
|  | **E2** | | **Tes** | | | **DHEA** | |
| **caffeine intake (yes/no)** | 0.017 | p=0.3286 | | 0.067 | p=0.00019 | 0.0347 | p=0.056 |
| **physical activity (yes/no)** | 0.0084 | p=0.6427 | | -0.002 | p = 0.907 | 0.0075 | p = 0.6789 |

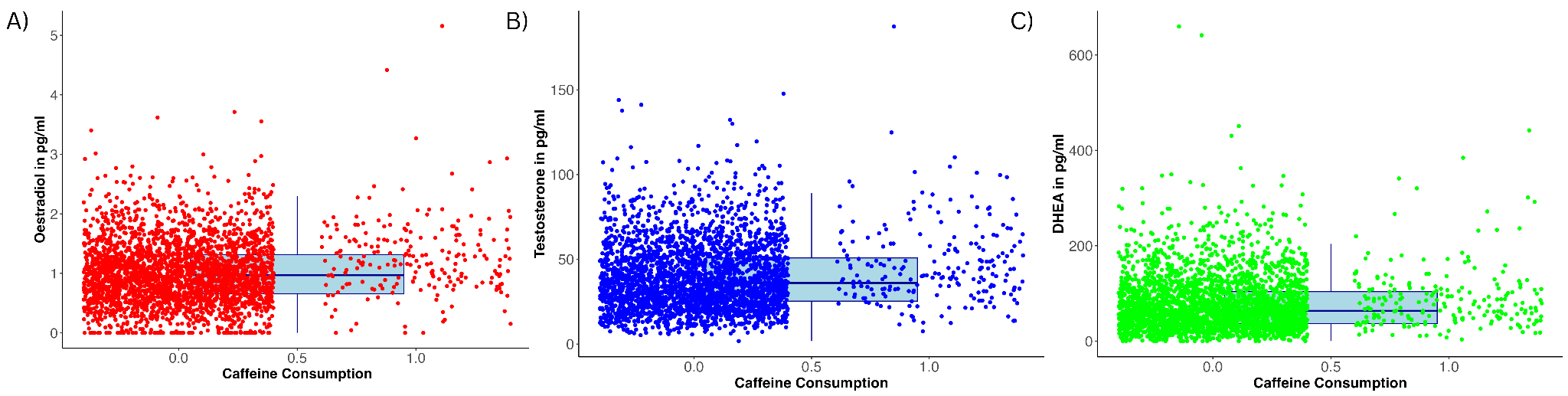

*Figure 2: Relation between steroid hormone levels and caffeine consumption A) E2 concentration in pg/ml B) Tes concentration in pg/ml C) Dhea concentration in pg/ml*

#### 2.3 Fibre tracts from AtlasTrack

Except for Fmaj, Fmin, and CC, all tracts are separately labelled for the left and right hemispheres.

| **Abbrev** | **Fiber Name** | **Connected Brain Regions** |
| --- | --- | --- |
| Fx | fornix | hippocampus & mammillary nuclei of hypothalamus |
| CgC | cingulate cingulum | cingulate gyrus & entorhinal cortex (cingulate portion) |
| CgH | parahippocampal cingulum | cingulate gyrus & entorhinal cortex (parahippocampal portion) |
| CST | corticospinal tract (pyramidal tract) | motor cortex & spinal cord |
| ATR | anterior thalamic radiations | thalamus & frontal lobe |
| UNC | uncinate | inferior frontal lobe & anterior temporal lobe |
| ILF | inferior longitudinal fasciculus | occipital lobe & temporal lobe |
| IFO | inferior frontal occipital fasciculus | occipital lobe & frontal lobe |
| Fmaj | forceps major^a^ | left occipital cortex & right occipital cortex |
| Fmin | forceps minor^a^ | left prefrontal cortex & right prefrontal cortex |
| CC | corpus callosum | left cortex & right cortex |
| SLF | superior longitudinal fasciculus | temporal and parietal lobes & frontal lobe |
| tSLF | temporal superior longitudinal fasciculus (arcuate fasciculus)^b^ | temporal lobe & frontal lobe |
| pSLF | parietal superior longitudinal fasciculus^b^ | parietal lobe & frontal lobe |
| SCS | superior corticostriate | superior cortex & striatum |
| fSCS | frontal superior corticostriate^c^ | superior frontal cortex & striatum |
| pSCS | parietal superior corticostriate^c^ | superior parietal cortex & striatum |
| SIFC | striatal inferior frontal cortex tract | inferior frontal cortex & striatum |
| IFSFC | inferior frontal to superior frontal cortical tract | inferior frontal cortex & superior frontal cortex |
| FXCUT | Fornix excluding fimbria | N/A |

a. *sub-tracts of the CC*
b. *sub-tracts of the SLF*
c. *sub-tracts of the SCS*

#### 2.4 Min, Max and IQR for each hormone

| **Hormones** | **Min – Max (in pg/ml)** | **IQR** | **Mean Intra-assay Variance** |
| --- | --- | --- | --- |
| E2 | 0 – 5.15 | 0.66 | 0.0079 |
| Tes | 1.75 - 187.35 | 25.51 | 5.982 |
| DHEA | 0 – 659.78 | 67.12 | 33.62 |

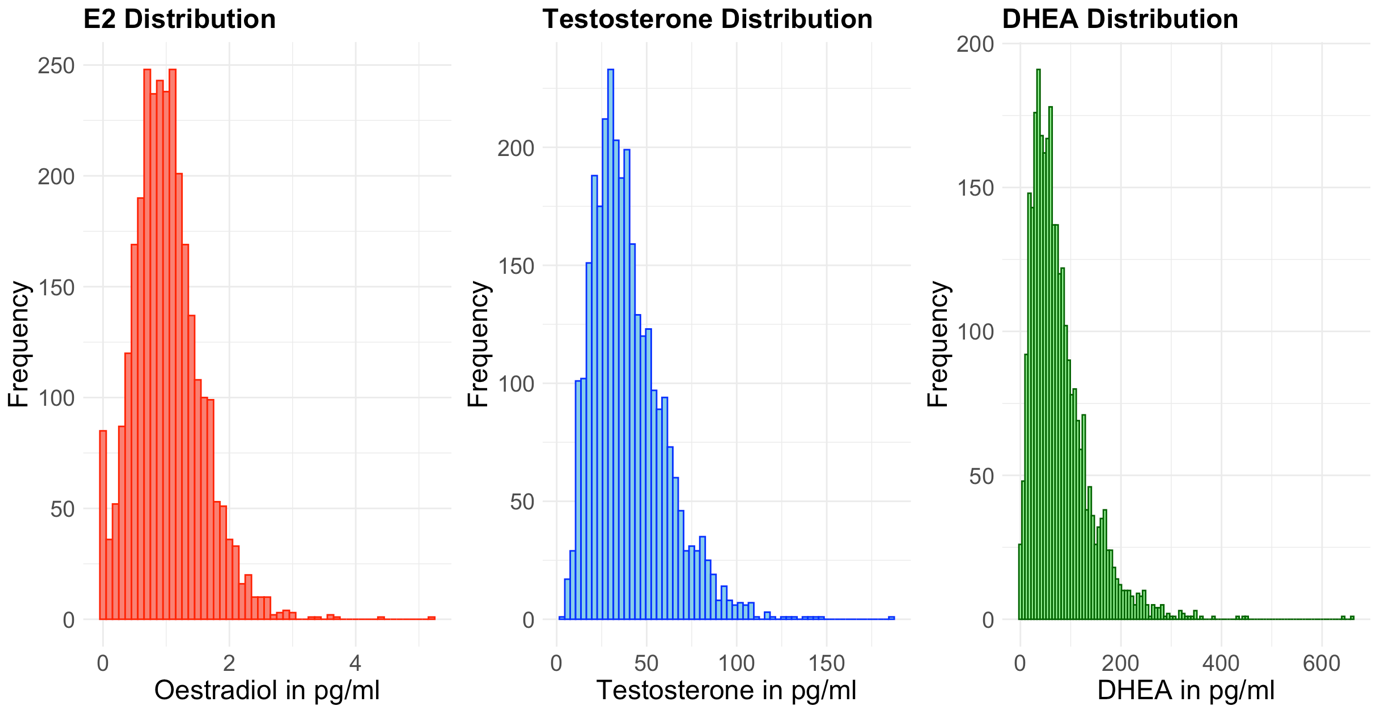

Figure 3: Distribution of the hormones

#### Table 2.5: demographic comparison between individuals at the baseline and 2-year-follow-up

|  | **T-value** | **p-value** |
| --- | --- | --- |
| Race/ethnicity | 0.70 | 0.48 |
| Parent combined income | 0.24 | 0.81 |
| Parent education | -1.4 | 0.16 |

#### Table 2.6: Hormone levels (pg/ml) in each Tanner stage

| **Tanner stage** | **n (%)** | **E2 (min-max; mean)** | **Tes (min-max; mean)** | **DHEA (min-max; mean)** |
| --- | --- | --- | --- | --- |
| 1: pre-puberty | 371 (12.27) | 0-2.92 (0.90) | 5.66 – 187.35 (30.29) | 0 – 268.99 (51.45) |
| 2: early puberty | 1128 (37.3) | 0-2.97 (0.95) | 5.21 – 101.46 (33.72) | 0 – 279.68 (64.91) |
| 3: mid puberty | 757 (25.03) | 0-3.27 (1.03) | 1.75 – 108.36 (41.48) | 4.95 – 341.11 (82.28) |
| 4: late puberty | 366 (12.10) | 0-5.15 (1.15) | 15.72 – 141.16 (52.27) | 2.46 – 641.46 (105.76) |
| 5: post-puberty | 208 (6.88) | 0-4.418 (1.32) | 18.58 – 147.73 (61.46) | (14.63 – 659.78) 132.43 |

### 3. Results:

#### A) With the full sample (including outliers)

3.1. Unimodality Analysis: Models running for each modality separately

###### 3.1.1 without Age-regress models:

| **Modality** | **Selected features from Unimodality (without age regress)** | | |
| --- | --- | --- | --- |
|  | **E2** | **Tes** | **DHEA** |
| sMRI | 18 | 30 | 32 |
| DTI | 14 | 25 | 19 |
| rs-fMRI | 5 | 19 | 24 |
| Task-based fMRI | 0 | 0 | 4 |
| Total | 37 | 74 | 79 |

###### 3.1.1.1 sMRI

| **Regions** | **Hemi** | **Features** | **Beta_coefficient** |
| --- | --- | --- | --- |
| **Estradiol (E2)** | | | |
| G and S frontomargin | Left | CTh | -0.0099128 |
| G and S transv frontopol | left | CTh | -0.0092838 |
| Lat Fis-ant-Vertical | left | CTh | 0.0057131 |
| Pole occipital | left | CTh | 0.0044942 |
| S oc middle and Lunatus | left | CTh | -0.0039915 |
| S oc sup and transversal | left | CTh | -0.0041765 |
| G and S frontomargin | right | CTh | -0.00438 |
| Lat Fis-post | right | CTh | -0.0052087 |
| S circular insula ant | right | CTh | -0.0066465 |
| S front middle | right | CTh | -0.0040605 |
| G and S paracentral | left | SDep | -0.0075984 |
| G temp sup-Plan tempo | Left | SDep | -0.0059958 |
| G temporal inf | Left | SDep | -0.0048858 |
| Lat Fis-ant-Horizont | Left | SDep | -0.0064826 |
| G temporal middle | Right | SDep | 0.0032257 |
| S collat transv post | Right | CAr | -0.0032783 |
| S occipital ant | Left | CVol | -0.0043258 |
| Pallidum | Right | Vol | 0.0042786 |
| **Testosterone (Tes)** | | | |
| Intercept |  |  | 39.8461 |
| G and S frontomargin | Left | CTh | -0.54936 |
| G and S transv frontopol | Left | CTh | -0.578945 |
| G and S cingul-Ant | Left | CTh | -0.62059 |
| G precentral | Left | CTh | 0.725588 |
| Pole occipital | Left | CTh | 0.593369 |
| S orbital-H Shaped | Left | CTh | -0.470509 |
| S precentral-inf-part | Left | CTh | 0.744805 |
| S subparietal | Left | CTh | -0.972735 |
| S temporal transverse | Left | CTh | 0.418503 |
| G and S transv frontopol | Right | CTh | -0.36725 |
| G cingul-Post-ventral | Right | CTh | -0.897626 |
| G precentral | Right | CTh | 0.915038 |
| G temp sup-Plan polar | Right | CTh | 0.610657 |
| Lat Fis-post | Right | CTh | -0.722308 |
| S calcarine | Right | CTh | -0.875186 |
| S suborbital | Right | CTh | -0.590426 |
| G and S frontomargin | Left | SDep | -0.34183 |
| G precentral | Left | SDep | 0.445486 |
| S central | Left | SDep | 0.364322 |
| S orbital-H Shaped | Left | SDep | -0.521855 |
| G front middle | Right | SDep | 0.420666 |
| G Ins lg and S cent ins | Right | SDep | 0.520994 |
| G temp sup-Lateral | Right | SDep | -0.799262 |
| S oc-temp med and Lingual | Right | SDep | -0.569058 |
| G precuneus | Left | CVol | -0.658246 |
| S oc middle and Lunatus | Right | CVol | -0.457678 |
| thalamus proper | Left | Vol | 0.476338 |
| Hippocampus | left | Vol | 1.03453 |
| cerebellum cortex | right | Vol | 0.556109 |
| accumbens area | right | Vol | -0.354618 |
| **DHEA** | | | |
| Intercept |  |  | 78.0399 |
| G precuneus | left | CVol | -2.07635 |
| S front middle | right | CTh | -2.00426 |
| S oc sup and transversal | Left | CTh | -1.78438 |
| G temp sup-Lateral | right | SDep | -1.57996 |
| S oc-temp lat | right | SDep | -1.55927 |
| G and S transv frontopol | left | CTh | -1.55106 |
| S orbital-H Shaped | left | SDep | -1.2412 |
| S calcarine | right | CTh | -1.23715 |
| S collat transv post | right | SDep | -1.23254 |
| Lat Fis-post | right | CTh | -1.18189 |
| S orbital lateral | right | CTh | -1.16335 |
| G and S frontomargin | Left | CTh | -1.14378 |
| G and S occipital inf | Left | CAr | -1.04344 |
| S oc middle and Lunatus | right | CVol | -0.959615 |
| S precentral-sup-part | right | CTh | -0.954621 |
| G and S transv frontopol | right | CVol | -0.904069 |
| G precuneus | left | SDep | -0.83679 |
| G parietal sup | left | CAr | -0.739335 |
| G occipital middle | Left | CVol | -0.736648 |
| S occipital ant | Left | CVol | -0.708319 |
| G and S cingul-Ant | Left | CTh | -0.686427 |
| G precentral | Left | CTh | 1.78876 |
| Thalamus proper | Left | Vol | 1.71105 |
| Pole occipital | Left | CVol | 1.55955 |
| G precentral | Left | SDep | 1.51242 |
| G Ins lg and S cent ins | right | SDep | 1.25514 |
| Pallidum | right | Vol | 1.17321 |
| G occipital sup | Left | CTh | 1.13417 |
| G precentral | Right | CTh | 1.00832 |
| S collat transv ant | left | CTh | 1.00515 |
| G front middle | left | SDep | 0.922084 |
| Pole temporal | right | CTh | 0.756934 |

###### 3.1.1.2 dMRI

| **Regions** | **Hemi** | **Features** | **Beta coefficient** |
| --- | --- | --- | --- |
| **Estradiol (E2)** | | | |
| Intercept | - | - | 1.0163 |
| FMIN | - | MD | 0.018954 |
| SCSLH | Left | FA | 0.01262 |
| CSTLH | Left | MD | 0.012376 |
| ATRRH | Right | FA | 0.012162 |
| FXCUTLH | Left | MD | 0.0091361 |
| FXCUTRH | Right | MD | 0.0087639 |
| UNCRH | Right | FA | 0.0086736 |
| SIFCLH | Left | FA | 0.007141 |
| CGCLH | Left | FA | 0.0063074 |
| CGCLH | Left | MD | -0.015674 |
| ATRLH | Left | MD | -0.013501 |
| ILFLH | Left | MD | -0.01157 |
| TSLFRH | Right | FA | -0.0092331 |
| FXCUTLH | Left | FA | -0.0059733 |
| **Testosterone (Tes)** | | | |
| Intercept | - | - | 39.8461 |
| IFOLH | Left | MD | 2.19376 |
| FMIN | - | MD | 1.77395 |
| SCSLH | Left | FA | 1.75995 |
| FMAJ | - | FA | 1.50559 |
| CGCLH | Left | FA | 1.35978 |
| UNCRH | Right | FA | 1.00817 |
| CGHLH | Left | FA | 0.916063 |
| ATRRH | Right | FA | 0.878993 |
| IFOLH | Left | FA | 0.814455 |
| SIFCLH | Left | FA | 0.680406 |
| PSLFLH | Left | MD | -1.63394 |
| CC | - | MD | -1.52461 |
| FMIN | - | FA | -1.49317 |
| CC | - | FA | -1.31304 |
| FSCSLH | Left | MD | -1.1635 |
| ILFRH | Right | MD | -1.0572 |
| PSCSRH | Right | FA | -1.02469 |
| IFSFCLH | Left | FA | -1.0222 |
| TSLFLH | Left | FA | -0.98132 |
| UNCLH | Left | FA | -0.911489 |
| FXRH | Right | FA | -0.869364 |
| CGHLH | Left | MD | -0.511567 |
| FXLH | Left | FA | -0.503042 |
| CGHRH | Right | MD | -0.451971 |
| IFSFCRH | Right | FA | -0.406353 |
| **DHEA** | | | |
| Intercept | - | - | 78.0376 |
| IFOLH | Left | MD | 3.11348 |
| FMAJ | - | FA | 2.25691 |
| FMIN | - | MD | 2.20467 |
| SCSLH | Left | FA | 1.88415 |
| CGHLH | Left | FA | 1.82493 |
| ILFRH | Right | FA | 1.64595 |
| CGCLH | Left | FA | 1.62553 |
| UNCRH | right | FA | 1.27492 |
| ILFRH | Right | MD | -1.83005 |
| PSLFLH | Left | MD | -1.64155 |
| ATRLH | Left | MD | -1.61543 |
| FXLH | Left | FA | -1.58069 |
| PSCSRH | Right | FA | -1.54671 |
| CC | - | FA | -1.46887 |
| UNCLH | Left | FA | -1.31711 |
| FSCSLH | Left | MD | -1.29629 |
| IFSFCRH | Right | FA | -1.29305 |
| FXCUTLH | Left | FA | -1.09094 |
| SLFLH | Left | FA | -1.02037 |

###### 3.1.1.3 fMRI

| **Regions** | **Hemi** | **Features** | **Beta_coefficient** |
| --- | --- | --- | --- |
| **DHEA** | | | |
| Intercept | - | - | 78.0399 |
| S oc-temp lat | right | FvP | 1.72801 |
| G and S frontomargin | right | PvN | 0.9847 |
| G oc-temp med-Parahip | Left | NvN | 0.826506 |
| S cingul-Marginalis | Left | PvN | -0.833702 |

###### 3.1.1.4 rsfMRI

| **Regions** | **Hemi** | **Features** | **Beta_coefficient** |
| --- | --- | --- | --- |
| **Estradiol (E2)** | | | |
| Intercept | - | - | 1.0163 |
| Retrospenial temporal network and right caudate | Right | Connectivity between cortical network and subcortical region | 0.0067894 |
| Retrospenial temporal network and left putamen | Left | Connectivity between cortical network and subcortical region | 0.00656 |
| Default mode network and left thalamus proper | Left | Connectivity between cortical network and subcortical region | -0.0099786 |
| Sensorymotor hand network and left caudate | Left | Connectivity between cortical network and subcortical region | -0.0050461 |
| Cingulo-opercular network and right pallidum | Right | Connectivity between cortical network and subcortical region | -0.0042359 |
| **Testosterone (Tes)** | | | |
| Intercept | - | - | 39.8461 |
| salience network and left accumbens area | Left | Connectivity between cortical network and subcortical region | 1.3334 |
| Sensory motor hand network and sensory motor mouth network | - | Inter cortical network | 0.928765 |
| retrosplenial temporal network and left putamen | left | Connectivity between cortical network and subcortical region | 0.881467 |
| dorsoattentional network and left caudate | Left | Connectivity between cortical network and subcortical region | 0.860504 |
| auditory network and cingulo-parietal network |  | Inter cortical network | 0.701936 |
| cingulo-opercular network and right caudate | Right | Connectivity between cortical network and subcortical region | 0.561755 |
| auditory network and left caudate | Left | Connectivity between cortical network and subcortical region | 0.492812 |
| default mode network and left caudate | Left | Connectivity between cortical network and subcortical region | 0.462185 |
| default mode network and retrosplenial temporal network |  | Inter cortical network | 0.42912 |
| sensorymotor hand network and right accumbens area | Right | Connectivity between cortical network and subcortical region | -1.03108 |
| default mode network and left hippocampus | Left | Connectivity between cortical network and subcortical region | -0.711736 |
| ventro-attentional network and left hippocampus | Left | Connectivity between cortical network and subcortical region | -0.688711 |
| retrosplenial temporal network and retrosplenial temporal network |  | Intra cortical network | -0.528833 |
| fronto-parietal network and right hippocampus | Right | Connectivity between cortical network and subcortical region | -0.517136 |
| cingulo-parietal network and default mode network |  | Inter cortical network | -0.503389 |
| cingulo-opercular and left cerebellum | Left | Connectivity between cortical network and subcortical region | -0.489716 |
| dorso-attentional network and right accumbens area | Right | Connectivity between cortical network and subcortical region | -0.447982 |
| visual network and right cerebellum | Right | Connectivity between cortical network and subcortical region | -0.370579 |
| salience network and visual network |  | Inter cortical network | -0.333476 |
| **DHEA** | | | |
| Intercept | - | - | 78.0394 |
| salience network and left accumbens area | Left | Connectivity between cortical network and subcortical region | 1.06036 |
| retrosplenial temporal network and left putamen | Left | Connectivity between cortical network and subcortical region | 0.781834 |
| auditory network and left caudate | Left | Connectivity between cortical network and subcortical region | 0.720094 |
| salience network and left cerebellum | Left | Connectivity between cortical network and subcortical region | 0.660471 |
| ventro-attentional network and left accumbens area | Left | Connectivity between cortical network and subcortical region | 0.618279 |
| cingulo-opercular network and right caudate | Right | Connectivity between cortical network and subcortical region | 0.609629 |
| sensorymotor hand network and sensory motor mouth network | - | Inter cortical network | 0.590746 |
| sensorymotor mouth network and sensorymotor hand network | - | Inter cortical network | 0.537112 |
| fronto-parietal network and ventro-attentional network | - | Inter cortical network | 0.422483 |
| cingulo-parietal network and cingulo-parietal network | - | Intra cortical network | 0.409021 |
| sensorymotor hand network and left caudate | Left | Connectivity between cortical network and subcortical region | -1.17233 |
| ventro-attentional network and left hippocampus | Left | Connectivity between cortical network and subcortical region | -0.961943 |
| visual network and right cerebellum | Right | Connectivity between cortical network and subcortical region | -0.954782 |
| default mode network and left hippocampus | Left | Connectivity between cortical network and subcortical region | -0.885434 |
| sensorymotor hand network and right accumbens area | Right | Connectivity between cortical network and subcortical region | -0.880152 |
| retrosplenial temporal network and retrosplenial temporal network |  | Intra cortical network | -0.82905 |
| default mode network and right caudate | Right | Connectivity between cortical network and subcortical region | -0.777956 |
| cingulo-opercular network and right pallidum | Right | Connectivity between cortical network and subcortical region | -0.731066 |
| default mode network and left thalamus proper | Left | Connectivity between cortical network and subcortical region | -0.70541 |
| dorso-attentional network and right putamen | Right | Connectivity between cortical network and subcortical region | -0.626737 |
| fronto-parietal network and right hippocampus | Right | Connectivity between cortical network and subcortical region | -0.620054 |
| retrosplenial temporal network and ventro-attentional network |  | Inter cortical network | -0.54432 |
| sensorymotor hand network and right cerebellum | Right | Connectivity between cortical network and subcortical region | -0.505056 |
| salience network and visual network |  | Inter cortical network | -0.487376 |

###### 3.1.2 Age-regress models:

| **Modality** | **Selected features from Unimodality (with age regress)** | | |
| --- | --- | --- | --- |
|  | **E2** | **Tes** | **DHEA** |
| sMRI | 5 | 61 | 30 |
| DTI | 0 | 0 | 0 |
| rs-fMRI | 2 | 3 | 3 |
| Task-based fMRI | 0 | 0 | 0 |
| Total | 7 | 64 | 33 |

###### 3.1.2.1 sMRI:

| **Region** | **Features name** | **Hemi** | **Beta_coefficient** |
| --- | --- | --- | --- |
| **Estradiol (E2)** | | | |
| Intercept | - | - | -1.565e-10 |
| G and S frontomargin | left | CTh | -0.007902 |
| G and S transv frontopol | Left | CTh | -0.008028 |
| S circular insula ant | right | CTh | -0.0056536 |
| G and S paracentral | left | SDep | -0.0071114 |
| G temp sup-Plan tempo | left | SDep | -0.0059398 |
| **Testosterone (Tes)** | | | |
| Intercept | - | - | 6.2753e-10 |
| G and S frontomargin | left | CTh | -0.054487 |
| G and S transv frontopol | Left | CTh | -0.07065 |
| G front inf-Triangul | Left | CTh | 0.071397 |
| G occipital sup | Left | CTh | 0.057402 |
| Lat Fis-ant-Vertical | Left | CTh | 0.067609 |
| Pole occipital | Left | CTh | 0.067693 |
| S subparietal | Left | CTh | -0.060714 |
| S temporal transverse | Left | CTh | 0.04524 |
| G and S transv frontopol | right | CTh | -0.059345 |
| G front inf-Opercular | right | CTh | 0.044427 |
| G front inf-Triangul | right | CTh | 0.043458 |
| G occipital sup | right | CTh | 0.0431 |
| G precentral | right | CTh | 0.046307 |
| G precuneus | right | CTh | 0.059496 |
| G temp sup-Lateral | right | CTh | 0.034495 |
| G temp sup-Plan polar | right | CTh | 0.053823 |
| Lat Fis-post | right | CTh | -0.06409 |
| S calcarine | right | CTh | -0.072794 |
| S circular insula ant | right | CTh | -0.075817 |
| S front middle | right | CTh | -0.046327 |
| S suborbital | right | CTh | -0.059562 |
| G and S frontomargin | left | SDep | -0.047694 |
| G and S transv frontopol | left | SDep | 0.047324 |
| G occipital sup | left | SDep | -0.056231 |
| G oc-temp lat-fusifor | left | SDep | 0.04471 |
| G precentral | left | SDep | 0.042906 |
| G precuneus | left | SDep | -0.080602 |
| S calcarine | left | SDep | -0.030436 |
| S intrapariet and P trans | left | SDep | 0.052136 |
| G front middle | right | SDep | 0.055257 |
| G Ins lg and S cent ins | right | SDep | 0.085978 |
| G temp sup-Lateral | right | SDep | -0.08188 |
| S precentral-sup-part | right | SDep | -0.066339 |
| G and S occipital inf | left | CAr | -0.086051 |
| G occipital middle | left | CAr | -0.070421 |
| G parietal sup | left | CAr | -0.045532 |
| G postcentral | left | CAr | -0.036173 |
| G precuneus | left | CAr | -0.055457 |
| S circular insula inf | left | CAr | 0.055174 |
| S collat transv post | left | CAr | -0.050188 |
| S oc middle and Lunatus | left | CAr | -0.052291 |
| S oc sup and transversal | left | CAr | -0.063157 |
| G and S occipital inf | right | CAr | -0.05769 |
| G front inf-Opercular | right | CAr | -0.04282 |
| G occipital sup | right | CAr | -0.03317 |
| S cingul-Marginalis | right | CAr | -0.04717 |
| S collat transv post | right | CAr | -0.046937 |
| S oc middle and Lunatus | right | CAr | -0.041152 |
| S oc sup and transversal | right | CAr | -0.047737 |
| G and S occipital inf | left | CVol | -0.069636 |
| G occipital middle | left | CVol | -0.065853 |
| G parietal sup | left | CVol | -0.03216 |
| G precuneus | left | CVol | -0.060542 |
| S oc middle and Lunatus | left | CVol | -0.055016 |
| S oc sup and transversal | left | CVol | -0.071066 |
| G and S occipital inf | right | CVol | -0.046224 |
| G and S transv frontopol | right | CVol | -0.02399 |
| S calcarine | right | CVol | -0.027154 |
| S collat transv post | right | CVol | -0.032531 |
| S front sup | right | CVol | -0.028524 |
| S oc middle and Lunatus | right | CVol | -0.05201 |
| **DHEA** | | | |
| Intercept | - | - | 2.3316e-08 |
| G and S frontomargin | Left | CTh | -0.29347 |
| G and S transv frontopol | Left | CTh | -0.41941 |
| G front inf-Triangul | Left | CTh | 0.21966 |
| G occipital sup | Left | CTh | 0.28776 |
| G temporal inf | Left | CTh | 0.25866 |
| S collat transv ant | Left | CTh | 0.31495 |
| S oc middle and Lunatus | Left | CTh | -0.31706 |
| S oc sup and transversal | Left | CTh | -0.36421 |
| G front inf-Triangul | right | CTh | 0.28295 |
| Lat Fis-post | right | CTh | -0.25383 |
| S calcarine | right | CTh | -0.23703 |
| S collat transv post | right | CTh | -0.29488 |
| S front middle | right | CTh | -0.51333 |
| G and S paracentral | left | SDep | -0.25117 |
| G front middle | left | SDep | 0.26313 |
| G precentral | left | SDep | 0.32718 |
| G precuneus | left | SDep | -0.42399 |
| G temp sup-Plan tempo | left | SDep | -0.22809 |
| G and S cingul-Mid-Post | right | SDep | -0.32199 |
| G front inf-Opercular | right | SDep | 0.29899 |
| G Ins lg and S cent ins | right | SDep | 0.38639 |
| G temp sup-Lateral | right | SDep | -0.33435 |
| S collat transv post | right | SDep | -0.40581 |
| S oc-temp lat | right | SDep | -0.44503 |
| G parietal sup | left | CAr | -0.30445 |
| G precuneus | left | CAr | -0.34869 |
| S collat transv post | right | CAr | -0.24695 |
| G occipital middle | left | CVol | -0.27003 |
| G precuneus | left | CVol | -0.38146 |
| S oc-temp lat | left | CVol | -0.22745 |

###### 3.1.2.2 rsfMRI:

| **Regions** | **Hemi** | **Features** | **Beta_coefficient** |
| --- | --- | --- | --- |
| **Estradiol (E2)** | | | |
| Intercept |  |  | -1.434e-10 |
| Default mode network and left thalamus proper | Left | Connectivity between cortical network and subcortical region | -0.00653 |
| Sensorymotor hand network and left caudate | Left | Connectivity between cortical network and subcortical region | -0.0030411 |
| **Testosterone (Tes)** | | | |
| Intercept | - | - | -3.2469e-10 |
| Cingulo-opercular network and left cerebellum | Left | Connectivity between cortical network and subcortical region | -0.18805 |
| Default mode network and left hippocampus | Left | Connectivity between cortical network and subcortical region | -0.11592 |
| Fronto parietal network and right hippocampus | Right | Connectivity between cortical network and subcortical region | -0.16134 |
| **DHEA** | | | |
| Intercept | - | - | 2.9097e-08 |
| Cingulo-opercular network and left cerebellum | Left | Connectivity between cortical network and subcortical region | -0.41599 |
| Cingulo-opercular network and left pallidum | Left | Connectivity between cortical network and subcortical region | -0.47596 |
| Sensorymotor hand network and left caudate | Left | Connectivity between cortical network and subcortical region | -0.9033 |

#### *3.2 Multimodality models:*

###### 3.2.1 without Age-regress models:

##### 3.2.1.1 E2

| **Regions** | **hemi** | **Features** | **Beta_coefficient** |
| --- | --- | --- | --- |
| Intercept | - | - | 1.0153 |
| FMIN | - | MD | 0.044345 |
| Default mode network and left thalamus proper | left | Connectivity between cortical network and subcortical region | -0.03565 |
| G and S paracentral | left | SDep | -0.033411 |
| G and S frontomargin | left | CTh | -0.027745 |
| SCSLH | left | FA | 0.027373 |
| ILFLH | left | MD | -0.027282 |
| CSTLH | left | MD | 0.027257 |
| Pole occipital | left | CTh | 0.026804 |
| Lat Fis-ant-Vertical | left | CTh | 0.026438 |
| Retrosplenial temporal network and left putamen | left | Connectivity between cortical network and subcortical region | 0.025613 |
| TSLFRH | right | FA | -0.025533 |
| ATRLH | left | MD | -0.025338 |
| Lat Fis-ant-Horizont | left | SDep | -0.025128 |
| G temp sup-Plan tempo | left | SDep | -0.023407 |
| S occipital ant | left | CVol | -0.023282 |
| cinguloopercular and right pallidum | right | Connectivity between cortical network and subcortical region | -0.02285 |
| Retrosplenial temporal network and right caudate | right | Connectivity between cortical network and subcortical region | 0.022328 |
| G and S transv frontopol | left | CTh | -0.021184 |
| CGCLH | left | MD | -0.020213 |
| S circular insula ant | right | CTh | -0.019891 |
| Somatomotor hand network and left caudate | left | Connectivity between cortical network and subcortical region | -0.019665 |
| S collat transv post | right | CAr | -0.019082 |
| ATRRH | right | FA | 0.01743 |
| Lat Fis-post | right | CTh | -0.016621 |
| pallidum | right | SubCVol | 0.015829 |
| S oc middle and Lunatus | left | CTh | -0.015086 |
| G temporal inf | left | SDep | -0.014511 |
| G temporal middle | right | SDep | 0.014326 |
| UNCRH | right | FA | 0.00995 |
| FXCUTLH | left | MD | 0.008963 |
| G and S frontomargin | right | CTh | -0.0071951 |

###### 3.2.1.2 Tes

| **Region** | **hemi** | **Features** | **Beta_coefficient** |
| --- | --- | --- | --- |
| Intercept | Intercept | Intercept | 39.8469 |
| FMIN | - | MD | 2.21121 |
| CC | - | MD | -1.75372 |
| IFOLH | Left | MD | 1.67885 |
| S calcarine | right | CTh | -1.63107 |
| Pole occipital | Left | CTh | 1.62599 |
| SCSLH | Left | FA | 1.59622 |
| PSLFLH | Left | MD | -1.49902 |
| FSCSLH | Left | MD | -1.44841 |
| G precuneus | Left | CVol | -1.41907 |
| hippocampus | Left | SubCVol | 1.32162 |
| Auditory network and cinguloparietal | - | Inter cortical network | 1.31864 |
| Cinguloopercular and left cerebellum | left | Connectivity between cortical network and subcortical region | -1.21865 |
| S subparietal | left | CTh | -1.21461 |
| Salience network and left accumbens area | Left | Connectivity between cortical network and subcortical region | 1.20132 |
| PSCSRH | right | FA | -1.18729 |
| G temp sup-Lateral | right | SDep | -1.14772 |
| S oc middle and Lunatus | right | CVol | -1.10507 |
| S precentral-inf-part | left | CTh | 1.05878 |
| Dorsoattentional and left caudate | left | Connectivity between cortical network and subcortical region | 1.04804 |
| G cingul-Post-ventral | right | CTh | -1.04477 |
| CC | - | FA | -1.02113 |
| Default mode network and retrosplenial temporal network | Left | Inter cortical network | 1.01363 |
| FMAJ | - | FA | 0.996282 |
| G Ins lg and S cent ins | right | SDep | 0.981812 |
| S temporal transverse | left | CTh | 0.977776 |
| Retrosplenial temporal network and left putamen | left | Connectivity between cortical network and subcortical region | 0.966155 |
| Lat Fis-post | right | CTh | -0.939898 |
| Sensorymotor mouth network and sensorymotor hand network | - | Inter cortical network | 0.93134 |
| G front middle | right | SDep | 0.911601 |
| G precentral | right | CTh | 0.871096 |
| Frontoparietal and right hippocampus | right | Connectivity between cortical network and subcortical region | -0.840471 |
| Default mode network and left caudate | left | Connectivity between cortical network and subcortical region | 0.838354 |
| cerebellum cortex | right | SubCVol | 0.831497 |
| G and S frontomargin | left | CTh | -0.830469 |
| FXRH | right | FA | -0.821209 |
| G precentral | left | CTh | 0.811658 |
| G precentral | left | SDep | 0.806511 |
| G temp sup-Plan polar | right | CTh | 0.806091 |
| Sensorymotor hand network and right accumbens area | right | Connectivity between cortical network and subcortical region | -0.802984 |
| Salience network and visual network | - | Inter cortical network | -0.790741 |
| FMIN | - | FA | -0.790726 |
| thalamus proper | left | SubCVol | 0.788643 |
| IFSFCLH | left | FA | -0.786519 |
| accumbens area | right | SubCVol | -0.780159 |
| S oc-temp med and Lingual | right | SDep | -0.771568 |
| UNCRH | right | FA | 0.736662 |
| Auditory and left caudate | left | Connectivity between cortical network and subcortical region | 0.700354 |
| G and S frontomargin | left | SDep | -0.68968 |
| Retrosplenial temporal network and retrosplenial temporal network | left | Intra cortical network | -0.689666 |
| FXLH | left | FA | -0.670864 |
| CGCLH | Left | FA | 0.655005 |
| CGHLH | Left | FA | 0.647778 |
| Cinguloopercular network and default mode network | - | Inter cortical network | -0.630611 |
| ILFRH | right | MD | -0.614166 |
| G and S transv frontopol | left | CTh | -0.587042 |
| S orbital-H Shaped | left | CTh | -0.584608 |
| S orbital-H Shaped | left | SDep | -0.580141 |
| Cinguloopercular network and right caudate | right | Connectivity between cortical network and subcortical region | 0.559716 |
| UNCLH | left | FA | -0.526138 |
| Visual network and right cerebellum | right | Connectivity between cortical network and subcortical region | -0.513981 |
| Dorsoattentional network and right accumbens area | right | Connectivity between cortical network and subcortical region | -0.499596 |
| S central | left | SDep | 0.48497 |
| SIFCLH | left | FA | 0.479996 |
| S suborbital | right | CTh | -0.45271 |
| IFOLH | left | FA | 0.43506 |
| ATRRH | right | FA | 0.424838 |
| CGHRH | right | MD | -0.423732 |
| Ventro-attentional and left hippocampus | left | Connectivity between cortical network and subcortical region | -0.420987 |
| Default mode network and left hippocampus | left | Connectivity between cortical network and subcortical region | -0.36884 |
| TSLFLH | left | FA | -0.365449 |
| G and S cingul-Ant | left | CTh | -0.337822 |

###### 3.2.1.3 DHEA

| **Region** | **hemi** | **Features** | **Beta_coefficient** |
| --- | --- | --- | --- |
| Intercept | Intercept | Intercept | 77.9898 |
| DMRI_DTIMD_FIBERAT_IFOLH | left | MD | 4.22404 |
| G occipital sup | left | CTh | 4.19485 |
| S oc sup and transversal | left | CTh | -3.86104 |
| G precuneus | left | CVol | -3.77746 |
| Pole occipital | left | CVol | 3.75556 |
| DMRI_DTIFA_FIBERAT_ILFRH | Right | FA | 3.73525 |
| DMRI_DTIFA_FIBERAT_SCSLH | left | FA | 3.3876 |
| G precentral | left | CTh | 3.35905 |
| S oc-temp lat | right | FvP | 3.25347 |
| Salience network and left cerebellum | left | Connectivity between cortical network and subcortical region | 3.15403 |
| Sensorymotor hand network left caudate | left | Connectivity between cortical network and subcortical region | -3.10267 |
| G Ins lg and S cent ins | right | SDep | 2.97238 |
| S collat transv post | right | SDep | -2.95619 |
| G oc-temp med-Parahip | left | NvN | 2.9367 |
| FMAJ | - | FA | 2.93634 |
| S oc middle and Lunatus | right | CVol | -2.92744 |
| S calcarine | right | CTh | -2.91791 |
| PSCSRH | right | FA | -2.80977 |
| G and S occipital inf | left | CAr | -2.769 |
| G precentral | left | SDep | 2.74906 |
| thalamus proper | left | SubCVol | 2.72236 |
| G temp sup-Lateral | right | SDep | -2.6363 |
| S cingul-Marginalis | left | PvN | -2.5964 |
| FMIN | - | MD | 2.57803 |
| retrosplenial temporal network and retrosplenial temporal network |  | Intra cortical network | -2.5668 |
| CGHLH | left | FA | 2.43834 |
| FXLH | Left | FA | -2.42129 |
| salience network and left accumbens area | left | Connectivity between cortical network and subcortical region | 2.36391 |
| G and S transv frontopol | left | CTh | -2.35115 |
| S oc-temp lat | right | SDep | -2.3398 |
| G and S transv frontopol | right | CVol | -2.21094 |
| G and S frontomargin | right | PvN | 2.20908 |
| Pole temporal | right | CTh | 2.19021 |
| S front middle | right | CTh | -2.1544 |
| sensorymotor hand network and right accumbens area | right | Connectivity between cortical network and subcortical region | -2.13698 |
| PSLFLH | left | MD | -2.11247 |
| CC | - | FA | -2.09407 |
| ventro-attentional network and left accumbens area | left | Connectivity between cortical network and subcortical region | 1.98863 |
| Lat Fis-post | right | CTh | -1.97366 |
| S occipital ant | left | CVol | -1.95992 |
| pallidum | right | SubCVol | 1.95719 |
| G front middle | left | SDep | 1.95653 |
| S collat transv ant | left | CTh | 1.94087 |
| S orbital lateral | right | CTh | -1.93704 |
| G and S frontomargin | left | CTh | -1.9361 |
| ILFRH | right | MD | -1.86363 |
| fronto-parietal network and ventro-attentional network |  | Inter cortical network | 1.85679 |
| S orbital-H Shaped | left | SDep | -1.82079 |
| default mode network and right caudate | right | Connectivity between cortical network and subcortical region | -1.7123 |
| G precuneus | left | SDep | -1.70357 |
| auditory network and left caudate | left | Connectivity between cortical network and subcortical region | 1.65703 |
| default mode network and left thalamus proper | left | Connectivity between cortical network and subcortical region | -1.57371 |
| visual network and right cerebellum | right | Connectivity between cortical network and subcortical region | -1.55539 |
| sensorymotor hand network and right cerebellum | right | Connectivity between cortical network and subcortical region | -1.55252 |
| cingulo-opercular network and right pallidum | right | Connectivity between cortical network and subcortical region | -1.5251 |
| S suborbital | right | CTh | -1.42183 |
| G parietal sup | left | CAr | -1.28278 |
| salience network and visual network |  | Inter cortical network | -1.26742 |
| retrosplenial temporal network and left putamen | left | Connectivity between cortical network and subcortical region | 1.21815 |
| dorso-attentional network and right putamen | right | Connectivity between cortical network and subcortical region | -1.2067 |
| default mode network and left hippocampus | left | Connectivity between cortical network and subcortical region | -1.10378 |
| ventro-attentional network and left hippocampus | left | Connectivity between cortical network and subcortical region | -0.94854 |

###### 3.2.2 Age-regress models:

| **Hormones** | **Selected Features from Multimodality** | **Selected Features from Unimodality** |
| --- | --- | --- |
| E2 | 7 | 7 |
| Tes | 43 | 64 |
| DHEA | 33 | 33 |

##### 3.2.2.1 E2

| **region** | **hemi** | **Features** | **B_ENet** |
| --- | --- | --- | --- |
| Intercept |  |  | 1.6942e-09 |
| Default mode network and left thalamus proper |  | Connectivity between cortical network and subcortical region | -0.036435 |
| G and S paracentral | left | SDep | -0.027315 |
| Sensory motor hand network and left caudate |  | Connectivity between cortical network and subcortical region | -0.026533 |
| G temp sup-Plan tempo | left | SDep | -0.024241 |
| G and S frontomargin | left | CTh | -0.020486 |
| G and S transv frontopol | left | CTh | -0.020196 |
| S circular insula ant | right | CTh | -0.019817 |

###### 3.2.2.2 Tes

| **region** | **hemi** | **Features** | **B_ENet** |
| --- | --- | --- | --- |
| Intercept |  |  | 9.498e-09 |
| Cingulo-opercular and left cerebellum |  | Connectivity between cortical network and subcortical region | -0.70486 |
| S circular insula inf | left | CAr | 0.56637 |
| G Ins lg and S cent ins | right | SDep | 0.53891 |
| Frontoparietal and right hippocampus |  | Connectivity between cortical network and subcortical region | -0.51214 |
| G front inf-Triangul | left | CTh | 0.5087 |
| S calcarine | right | CTh | -0.501 |
| Lat Fis-post | right | CTh | -0.4948 |
| G precuneus | left | SDep | -0.47455 |
| G and S transv frontopol | left | CTh | -0.45348 |
| Pole occipital | left | CTh | 0.44925 |
| Lat Fis-ant-Vertical | left | CTh | 0.43979 |
| S circular insula ant | right | CTh | -0.43277 |
| G oc-temp lat-fusifor | left | SDep | 0.4235 |
| S subparietal | left | CTh | -0.42152 |
| Default mode network and left hippocampus |  | Connectivity between cortical network and subcortical region | -0.41338 |
| G precuneus | right | CTh | 0.41256 |
| G and S frontomargin | left | CTh | -0.40833 |
| G and S transv frontopol | right | CTh | -0.4075 |
| S precentral-sup-part | right | SDep | -0.39021 |
| S suborbital | right | CTh | -0.35806 |
| S intrapariet and P trans | left | SDep | 0.34866 |
| G temp sup-Plan polar | right | CTh | 0.34519 |
| G temp sup-Lateral | right | SDep | -0.3365 |
| G occipital sup | left | CTh | 0.33578 |
| G occipital sup | right | CTh | 0.33231 |
| S temporal transverse | left | CTh | 0.33207 |
| G and S transv frontopol | left | SDep | 0.33095 |
| G front middle | right | SDep | 0.32822 |
| G and S frontomargin | left | SDep | -0.31266 |
| S cingul-Marginalis | right | CAr | -0.30704 |
| G precentral | right | CTh | 0.30294 |
| G occipital sup | left | SDep | -0.28439 |
| G front inf-Triangul | right | CTh | 0.25583 |
| S front middle | right | CTh | -0.23627 |
| G and S occipital inf | left | CAr | -0.21844 |
| G precentral | left | SDep | 0.21384 |
| G precuneus | left | CVol | -0.20095 |
| S collat transv post | right | CAr | -0.19839 |
| G front inf-Opercular | right | CTh | 0.19516 |
| S oc middle and Lunatus | right | CVol | -0.19432 |
| S oc sup and transversal | left | CVol | -0.17743 |
| S oc sup and transversal | right | CAr | -0.16133 |
| S oc middle and Lunatus | left | CVol | -0.15896 |

###### 3.2.2.3 DHEA

| **region** | **hemi** | **Features** | **B_ENet** |
| --- | --- | --- | --- |
| Intercept |  |  | -1.4821e-07 |
| G occipital sup | left | CTh | 3.6758 |
| Sensorymotor hand network and left caudate |  | Connectivity between cortical network and subcortical region | -3.5047 |
| S front middle | right | CTh | -2.7901 |
| S oc-temp lat | right | SDep | -2.7758 |
| G and S transv frontopol | left | CTh | -2.5472 |
| S collat transv post | right | SDep | -2.3924 |
| G front inf-Triangul | right | CTh | 2.3365 |
| S collat transv ant | left | CTh | 2.2571 |
| G precuneus | left | SDep | -2.2252 |
| Cingulo-opercular and left cerebellum |  | Connectivity between cortical network and subcortical region | -2.1747 |
| G and S cingul-Mid-Post | right | SDep | -2.0791 |
| Cinguloopercular and right pallidum |  | Connectivity between cortical network and subcortical region | -1.994 |
| S oc sup and transversal | left | CTh | -1.9785 |
| G temporal inf | left | CTh | 1.9605 |
| G front inf-Triangul | left | CTh | 1.9568 |
| S oc middle and Lunatus | left | CTh | -1.8892 |
| G Ins lg and S cent ins | right | SDep | 1.8267 |
| G temp sup-Lateral | right | SDep | -1.8043 |
| G front inf-Opercular | right | SDep | 1.7694 |
| G precentral | left | SDep | 1.7443 |
| G and S frontomargin | left | CTh | -1.7424 |
| G front middle | left | SDep | 1.6247 |
| G precuneus | left | CVol | -1.5216 |
| S collat transv post | right | CTh | -1.507 |
| G and S paracentral | left | SDep | -1.4179 |
| G temp sup-Plan tempo | left | SDep | -1.4047 |
| G occipital middle | left | CVol | -1.3914 |
| Lat Fis-post | right | CTh | -1.27 |
| S oc-temp lat | left | CVol | -1.2267 |
| S collat transv post | right | CAr | -1.1851 |
| G parietal sup | left | CAr | -0.91114 |
| S calcarine | right | CTh | -0.846 |

#### B) With the selected sample (excluded outliers)

##### 3.1. Unimodality Analysis:

###### 3.1.1 without Age-regress models:

| **Modality** | **Selected features from Unimodality** | | |
| --- | --- | --- | --- |
|  | **E2** | **Tes** | **DHEA** |
| sMRI | 19 | 33 | 31 |
| DTI | 0 | 1 | 1 |
| rs-fMRI | 4 | 22 | 49 |
| Task-based fMRI | 0 | 0 | 2 |
| Total | 26 | 85 | 100 |

###### 3.1.1.1 sMRI

| **Regions** | **Hemi** | **Features** | **Beta_coefficient** |
| --- | --- | --- | --- |
| **Estradiol (E2)** | | | |
| Intercept | - | - | 0.99073 |
| Pallidum | right | SCVol | 0.0036995 |
| S collat transv ant | right | CAr | 0.0036438 |
| Lat Fis-ant-Vertical | left | CTh | 0.003275 |
| G and S frontomargin | left | CTh | -0.0073598 |
| G and S transv frontopol | left | CTh | -0.0069951 |
| G temp sup-Plan tempo | left | SDep | -0.0048272 |
| G front inf-Orbital | left | SDep | -0.0039791 |
| G temporal inf | left | SDep | -0.0039178 |
| G and S paracentral | left | SDep | -0.003655 |
| S circular insula ant | right | CTh | -0.003454 |
| G and S frontomargin | right | CTh | -0.0033796 |
| G precentral | left | CAr | -0.0033476 |
| Lat Fis-post | right | CTh | -0.0032855 |
| S oc-temp med and Lingual | right | CTh | -0.0031277 |
| G oc-temp lat-fusifor | left | CVol | -0.0030188 |
| G and S occipital inf | left | CAr | -0.0028192 |
| G and S cingul-Mid-Post | right | SDep | -0.0028145 |
| G and S cingul-Mid-Post | left | CAr | -0.0022774 |
| S oc-temp med and Lingual | right | CVol | -0.0019987 |
| **Testosterone (Tes)** | | | |
| Intercept | - | - | 38.8757 |
| Hippocampus | left | SCVol | 0.967431 |
| Pole occipital | left | CTh | 0.772019 |
| G precentral | right | CTh | 0.730934 |
| cerebellum cortex | right | SCVol | 0.502079 |
| Pallidum | right | SCVol | 0.490419 |
| S temporal transverse | left | CTh | 0.46471 |
| G temp sup-Plan polar | right | CTh | 0.463026 |
| thalamus proper | left | SCVol | 0.438107 |
| G Ins lg and S cent ins | right | SDep | 0.430153 |
| S precentral-sup-part | right | CTh | 0.423826 |
| G front middle | left | SDep | 0.357282 |
| S calcarine | right | CTh | -0.804736 |
| G cingul-Post-ventral | right | CTh | -0.786947 |
| S subparietal | left | CTh | -0.734204 |
| G temp sup-Lateral | right | SDep | -0.707271 |
| S precentral-inf-part | right | SDep | -0.665838 |
| S oc-temp med and Lingual | right | SDep | -0.529328 |
| G precuneus | left | CVol | -0.52346 |
| Lat Fis-post | right | CTh | -0.513084 |
| G and S transv frontopol | left | CTh | -0.499631 |
| G and S cingul-Ant | left | CTh | -0.488952 |
| G and S frontomargin | left | CTh | -0.463353 |
| G postcentral | left | CAr | -0.458097 |
| G and S transv frontopol | right | CVol | -0.445891 |
| S suborbital | right | CTh | -0.445721 |
| Lat Fis-post | left | CTh | -0.439762 |
| G and S occipital inf | left | CAr | -0.400719 |
| G subcallosal | left | CTh | -0.387845 |
| G front inf-Orbital | left | SDep | -0.382542 |
| S orbital-H Shaped | left | SDep | -0.338632 |
| G oc-temp med-Lingual | right | CTh | -0.332028 |
| G oc-temp lat-fusifor | left | CTh | -0.277406 |
| G and S cingul-Ant | right | SDep | -0.257884 |
| **DHEA** | | | |
| Intercept | - | - | 74.1861 |
| pallidum | right | SCVol | 1.5231 |
| Pole occipital | left | CTh | 1.45157 |
| G precentral | left | SDep | 1.15631 |
| G front middle | left | SDep | 1.02629 |
| G precentral | left | CTh | 0.772617 |
| G Ins lg and S cent ins | right | SDep | 0.759853 |
| G precentral | right | CTh | 0.561245 |
| G oc-temp med-Parahip | right | CAr | 0.463391 |
| G precuneus | left | CVol | -1.70633 |
| G and S transv frontopol | left | CTh | -1.44709 |
| Lat Fis-post | right | CTh | -1.36858 |
| G and S cingul-Mid-Post | right | SDep | -1.20698 |
| G temp sup-Lateral | right | SDep | -1.17066 |
| S front middle | right | CTh | -1.06989 |
| S temporal inf | right | CVol | -1.0317 |
| S oc middle and Lunatus | right | CVol | -1.00849 |
| S calcarine | right | CTh | -0.942255 |
| G and S transv frontopol | right | CVol | -0.903418 |
| S precentral-inf-part | left | SDep | -0.890161 |
| G and S frontomargin | left | CTh | -0.859239 |
| G front inf-Triangul | right | SDep | -0.835207 |
| S suborbital | right | CTh | -0.806518 |
| S orbital-H Shaped | left | SDep | -0.799095 |
| S oc sup and transversal | left | CTh | -0.664681 |
| G and S occipital inf | left | CAr | -0.647996 |
| G postcentral | left | CAr | -0.617278 |
| G oc-temp med-Lingual | left | CTh | -0.573018 |
| S orbital-H Shaped | right | CTh | -0.566231 |
| S oc middle and Lunatus | left | CTh | -0.517402 |
| G temp sup-Lateral | right | CAr | -0.493995 |
| S collat transv post | right | SDep | -0.488303 |

###### 3.1.1.2 dMRI

| **Regions** | **Hemi** | **Features** | **Beta_coefficient** |
| --- | --- | --- | --- |
| **Estradiol (E2)** | | | |
| Intercept | - | - | 0.99078 |
| FMIN | - | MD | 0.016743 |
| SCSLH | left | FA | 0.013779 |
| CGCLH | left | MD | -0.012501 |
| **Testosterone (Tes)** | | | |
| Intercept | - | - | 38.8769 |
| IFOLH | Left | MD | 2.0378 |
| FMIN | - | MD | 1.65707 |
| SCSLH | Left | FA | 1.55686 |
| CGCLH | Left | FA | 1.15491 |
| FMAJ | - | FA | 1.08097 |
| ATRRH | Right | FA | 0.742776 |
| CGHLH | Left | FA | 0.738863 |
| IFOLH | Left | FA | 0.720825 |
| UNCRH | Right | FA | 0.393181 |
| SIFCLH | Left | FA | 0.379325 |
| CSTRH | Right | FA | 0.16846 |
| CC | - | MD | -1.43376 |
| PSLFLH | Left | MD | -1.31036 |
| CC | - | FA | -1.10741 |
| FMIN | - | FA | -1.04412 |
| TSLFLH | Left | FA | -0.907934 |
| IFSFCLH | Left | FA | -0.902114 |
| FSCSLH | Left | MD | -0.771508 |
| FXRH | Right | FA | -0.718207 |
| FXLH | Left | FA | -0.701401 |
| ILFRH | Right | MD | -0.679573 |
| IFSFCRH | Right | FA | -0.645712 |
| CGHRH | Right | MD | -0.540957 |
| UNCLH | Left | FA | -0.538928 |
| PSCSRH | Right | FA | -0.52936 |
| FSCSRH | Right | MD | -0.520955 |
| PSLFRH | Right | MD | -0.509278 |
| CGHLH | Left | MD | -0.507232 |
| SIFCRH | Right | MD | -0.33852 |
| FXCUTLH | Left | FA | -0.179332 |
| **DHEA** | | | |
| Intercept | - | - | 74.1861 |
| IFOLH | Left | MD | 2.31697 |
| FMIN | - | MD | 1.89895 |
| CGHLH | Left | FA | 1.71986 |
| CGCLH | Left | FA | 1.15607 |
| SCSLH | Left | FA | 1.07554 |
| FXCUTRH | Right | FA | 1.05014 |
| FMAJ | - | FA | 1.01401 |
| ATRRH | Right | FA | 0.862026 |
| FXCUTRH | Right | MD | 0.548129 |
| FXCUTLH | Left | MD | 0.414468 |
| FXLH | Left | FA | -2.07425 |
| ATRLH | Left | MD | -1.56166 |
| IFSFCRH | Right | FA | -1.43606 |
| CC |  | MD | -1.37741 |
| ILFRH | Right | MD | -1.16132 |
| TSLFLH | Left | FA | -1.04719 |
| PSLFRH | Right | MD | -0.835864 |
| CGHRH | right | MD | -0.65319 |

###### 3.1.1.3 fMRI

| **Regions** | **Hemi** | **Features** | **Beta_coefficient** |
| --- | --- | --- | --- |
| **DHEA** | | | |
| Intercept |  |  | 74.1861 |
| S oc-temp lat | Right | FvP | 1.08808 |
| S oc-temp lat | left | FvP | 0.6991 |

###### 3.1.1.4 rsfMRI

| **Regions** | **Hemi** | **Features** | | **Beta_coefficient** |
| --- | --- | --- | --- | --- |
| **Estradiol (E2)** | | | | |
| Intercept |  |  | | 0.99078 |
| retrosplenial temporal network and left putamen | left | Connectivity between cortical network and subcortical region | | 0.0056651 |
| default mode network and left thalamus proper | left | Connectivity between cortical network and subcortical region | | -0.010496 |
| cingulo-opercular and left cerebellum | left | Connectivity between cortical network and subcortical region | | -0.0065366 |
| sensorymotor hand network and left caudate | left | Connectivity between cortical network and subcortical region | | -0.0022522 |
| **Testosterone (Tes)** | | | | |
| Intercept | - | | - | 38.8769 |
| salience network and left accumbens area | left | | Connectivity between cortical network and subcortical region | 1.06456 |
| retrosplenial temporal network and left putamen | left | | Connectivity between cortical network and subcortical region | 0.810473 |
| sensorymotor hand network and sensory motor mouth network | - | | Inter cortical network | 0.724543 |
| dorsoattentional network and left caudate | left | | Connectivity between cortical network and subcortical region | 0.656379 |
| auditory network and cingulo-parietal network | - | | Inter cortical network | 0.585297 |
| cingulo-opercular network and right caudate | right | | Connectivity between cortical network and subcortical region | 0.522728 |
| default mode network and left hippocampus | left | | Connectivity between cortical network and subcortical region | -0.77906 |
| sensorymotor hand network and right accumbens area | right | | Connectivity between cortical network and subcortical region | -0.610073 |
| fronto-parietal network and right hippocampus | right | | Connectivity between cortical network and subcortical region | -0.561173 |
| ventro-attentional network and left hippocampus | left | | Connectivity between cortical network and subcortical region | -0.524961 |
| cingulo-parietal network and default mode network | - | | Inter cortical network | -0.432118 |
| cingulo-opercular and left cerebellum | left | | Connectivity between cortical network and subcortical region | -0.418948 |
| default mode network and ventro-attentional network | - | | Inter cortical network | -0.391779 |
| dorso-attentional network and left hippocampus | left | | Connectivity between cortical network and subcortical region | -0.341795 |
| retrosplenial temporal network and retrosplenial temporal network | - | | Intra cortical network | -0.332988 |
| salience network and visual network | - | | Inter cortical network | -0.276435 |
| dorso-attentional network and right accumbens area | right | | Connectivity between cortical network and subcortical region | -0.26928 |
| auditory network and dorso-attentional network | - | | Inter cortical network | -0.231156 |
| sensorymotor mouth network and visual network | - | | Inter cortical network | -0.140845 |
| default mode network and cingulo-opercular network | - | | Inter cortical network | -0.102461 |
| ventro-attentional network and default mode network | - | | Inter cortical network | -0.100582 |
| visual network and sensory motor mouth network | - | | Inter cortical network | -0.0237346 |
| **DHEA** | | | | |
| Intercept |  | |  | 74.1861 |
| salience network and left accumbens area | left | | Connectivity between cortical network and subcortical region | 1.26694 |
| retrosplenial temporal network and left putamen | left | | Connectivity between cortical network and subcortical region | 0.889809 |
| dorsoattentional network and left caudate | left | | Connectivity between cortical network and subcortical region | 0.604846 |
| cingulo-opercular network and right caudate | right | | Connectivity between cortical network and subcortical region | 0.581662 |
| sensorymotor hand network and sensory motor mouth network | - | | Inter cortical network | 0.554002 |
| auditory network and right caudate | right | | Connectivity between cortical network and subcortical region | 0.544118 |
| ventro-attentional network and left accumbens area | left | | Connectivity between cortical network and subcortical region | 0.517035 |
| sensorymotor mouth network and sensorymotor hand network | - | | Inter cortical network | 0.468898 |
| salience network and right pallidum | right | | Connectivity between cortical network and subcortical region | 0.42165 |
| sensorymotor hand network and salience network | - | | Inter cortical network | 0.372347 |
| auditory network and cingulo-parietal network | - | | Inter cortical network | 0.355227 |
| sensorymotor mouth network and left putamen | left | | Connectivity between cortical network and subcortical region | 0.347037 |
| default mode network and retrosplenial temporal network | - | | Inter cortical network | 0.337987 |
| salience network and sensorymotor hand network | - | | Inter cortical network | 0.303994 |
| cingulo-parietal network and auditory network | - | | Inter cortical network | 0.288032 |
| retrosplenial temporal network and default mode network | - | | Inter cortical network | 0.28029 |
| cingulo-parietal network and cingulo-parietal network | - | | Intra cortical network | 0.224125 |
| default mode network and left hippocampus | left | | Connectivity between cortical network and subcortical region | -1.02726 |
| fronto-parietal network and right hippocampus | right | | Connectivity between cortical network and subcortical region | -0.925244 |
| sensorymotor hand network and left caudate | left | | Connectivity between cortical network and subcortical region | -0.92225 |
| retrosplenial temporal network and retrosplenial temporal network | - | | Intra cortical network | -0.920082 |
| sensorymotor mouth network and right accumbens area | right | | Connectivity between cortical network and subcortical region | -0.788984 |
| cingulo-opercular and left cerebellum | left | | Connectivity between cortical network and subcortical region | -0.747221 |
| sensorymotor hand network and right accumbens area | right | | Connectivity between cortical network and subcortical region | -0.730223 |
| default mode network and right caudate | right | | Connectivity between cortical network and subcortical region | -0.697396 |
| ventro-attentional network and left hippocampus | left | | Connectivity between cortical network and subcortical region | -0.660758 |
| visual network and right cerebellum | right | | Connectivity between cortical network and subcortical region | -0.505851 |
| dorso-attentional network and left cerebellum | left | | Connectivity between cortical network and subcortical region | -0.495171 |
| dorso-attentional network and right accumbens area | right | | Connectivity between cortical network and subcortical region | -0.472652 |
| cingulo-opercular network and right putamen | right | | Connectivity between cortical network and subcortical region | -0.466176 |
| retrosplenial temporal network and right amygdala | right | | Connectivity between cortical network and subcortical region | -0.415155 |
| retrosplenial temporal network and ventro-attentional network | - | | Inter cortical network | -0.4108 |
| cingulo-parietal network and left accumbens area | left | | Connectivity between cortical network and subcortical region | -0.380655 |
| fronto-parietal network and left caudate | left | | Connectivity between cortical network and subcortical region | -0.379995 |
| ventro-attentional network and retrosplenial temporal network | - | | Inter cortical network | -0.37366 |
| salience network and visual network | - | | Inter cortical network | -0.368371 |
| default mode network and left thalamus proper | left | | Connectivity between cortical network and subcortical region | -0.366751 |
| auditory network and left pallidum | left | | Connectivity between cortical network and subcortical region | -0.36003 |
| salience network and right thalamus proper | right | | Connectivity between cortical network and subcortical region | -0.349518 |
| salience network and left caudate | left | | Connectivity between cortical network and subcortical region | -0.314192 |
| default mode network and right putamen | right | | Connectivity between cortical network and subcortical region | -0.313863 |
| visual network and salience network | - | | Inter cortical network | -0.298529 |
| auditory network and dorso-attentional network | - | | Inter cortical network | -0.294637 |
| default mode network and right thalamus proper | right | | Connectivity between cortical network and subcortical region | -0.281629 |
| default mode network and right putamen | right | | Connectivity between cortical network and subcortical region | -0.271005 |
| salience network and right amygdala | right | | Connectivity between cortical network and subcortical region | -0.260116 |
| retrosplenial temporal network and visual network | - | | Inter cortical network | -0.258 |
| dorso-attentional network and auditory network | - | | Inter cortical network | -0.249862 |
| visual network and retrosplenial temporal network | - | | Inter cortical network | -0.232756 |

###### 3.1.2 Age-regress models:

| **Modality** | **Selected features from Unimodality (with age regress)** | | |
| --- | --- | --- | --- |
|  | **E2** | **Tes** | **DHEA** |
| sMRI | 4 | 116 | 139 |
| DTI | 1 | 1 | 1 |
| Rs-fMRI | 4 | 23 | 7 |
| Task-based MRI | 0 | 0 | 0 |
| Total | 9 | 141 | 146 |

###### 3.1.2.1 sMRI:

| **Regions** | **Hemi** | **Features** | **Beta_coefficient** |
| --- | --- | --- | --- |
| **Estradiol (E2)** | | | |
| Intercept | - | - | -2.4967e-10 |
| G and S transv frontopol | left |  | -0.0068442 |
| G and S frontomargin | left |  | -0.0064122 |
| G temp sup-Plan tempo | left |  | -0.0040295 |
| G oc-temp lat-fusifor | left |  | -0.0019098 |
| **Testosterone (Tes)** | | | |
| Intercept | - | - | -1.6898e-08 |
| G Ins lg and S cent ins | right | SDep | 0.039244 |
| Pole occipital | left | CTh | 0.036094 |
| Lat Fis-ant-Vertical | left | CTh | 0.033949 |
| S intrapariet and P trans | left | SDep | 0.03084 |
| S pericallosal | right | SDep | 0.030668 |
| G precuneus | right | CTh | 0.029293 |
| G front middle | left | SDep | 0.028121 |
| S temporal transverse | left | CTh | 0.027426 |
| G front inf-Triangul | left | CTh | 0.02635 |
| G oc-temp med-Parahip | right | SDep | 0.02467 |
| S subparietal | right | CVol | 0.024122 |
| S circular insula inf | left | CAr | 0.023985 |
| G temp sup-G T transv | right | SDep | 0.022417 |
| G oc-temp med-Parahip | left | SDep | 0.022383 |
| G precentral | right | CTh | 0.021971 |
| G temp sup-Plan polar | right | CTh | 0.021847 |
| Lat Fis-ant-Horizont | left | CVol | 0.021666 |
| G temp sup-G T transv | left | SDep | 0.02131 |
| S circular insula ant | right | SDep | 0.020031 |
| Thalamus proper | right | SCVol | 0.019074 |
| S precentral-sup-part | right | CTh | 0.018444 |
| G front inf-Triangul | right | CTh | 0.018267 |
| Cerebellum cortex | right | SCVol | 0.018193 |
| G front inf-Opercular | right | SDep | 0.017804 |
| G occipital sup | left | CTh | 0.01683 |
| Lat Fis-ant-Horizont | left | CAr | 0.015812 |
| G and S transv frontopol | left | SDep | 0.015317 |
| G precentral | left | SDep | 0.015256 |
| G pariet inf-Angular | left | CTh | 0.014825 |
| G temp sup-Plan polar | left | CTh | 0.013547 |
| G precentral | right | SDep | 0.013498 |
| G temp sup-Lateral | right | CTh | 0.013303 |
| G and S occipital inf | left | CAr | -0.044222 |
| G temp sup-Lateral | right | SDep | -0.041971 |
| S calcarine | right | CTh | -0.040865 |
| G occipital middle | left | CAr | -0.039277 |
| G and S cingul-Ant | right | SDep | -0.03866 |
| S oc middle and Lunatus | right | CVol | -0.037726 |
| S precentral-inf-part | right | SDep | -0.037356 |
| G and S transv frontopol | left | CTh | -0.037109 |
| S oc middle and Lunatus | left | CVol | -0.036328 |
| G temp sup-Plan tempo | left | SDep | -0.035644 |
| S oc middle and Lunatus | left | CAr | -0.035548 |
| G occipital middle | left | CVol | -0.035083 |
| S oc middle and Lunatus | right | CAr | -0.034512 |
| G and S cingul-Mid-Post | right | SDep | -0.034235 |
| G oc-temp lat-fusifor | left | CTh | -0.034218 |
| G and S occipital inf | left | CVol | -0.033208 |
| Lat Fis-post | right | CTh | -0.032399 |
| S calcarine | left | SDep | -0.031978 |
| G and S occipital inf | right | CAr | -0.031442 |
| G precuneus | left | CAr | -0.031212 |
| G and S frontomargin | left | CTh | -0.030147 |
| S parieto occipital | left | CTh | -0.03006 |
| S circular insula ant | right | CTh | -0.029642 |
| Lat Fis-post | left | CTh | -0.028911 |
| G precuneus | left | SDep | -0.027707 |
| G occipital sup | left | SDep | -0.027704 |
| G precuneus | left | CVol | -0.027572 |
| S temporal inf | right | CVol | -0.027569 |
| G subcallosal | left | CTh | -0.026648 |
| S oc sup and transversal | right | CAr | -0.026624 |
| G and S transv frontopol | right | CTh | -0.026434 |
| S interm prim-Jensen | right | CTh | -0.025708 |
| G postcentral | left | CAr | -0.025484 |
| S collat transv post | right | CAr | -0.025036 |
| G cingul-Post-ventral | right | CTh | -0.024984 |
| G pariet inf-Angular | left | SDep | -0.024818 |
| S subparietal | left | CTh | -0.024468 |
| G and S transv frontopol | right | CVol | -0.023708 |
| G and S occipital inf | right | CVol | -0.023516 |
| G occipital sup | right | CAr | -0.022956 |
| S oc sup and transversal | left | CVol | -0.022363 |
| G front inf-Triangul | right | SDep | -0.022212 |
| S front inf | left | CVol | -0.022087 |
| G precentral | left | CAr | -0.02171 |
| G occipital sup | left | CAr | -0.021685 |
| G cuneus | right | CAr | -0.021245 |
| S suborbital | right | CTh | -0.020961 |
| S oc-temp lat | right | SDep | -0.020625 |
| S cingul-Marginalis | right | CAr | -0.020498 |
| G parietal sup | left | CAr | -0.020251 |
| S cingul-Marginalis | right | CVol | -0.020115 |
| S calcarine | left | CTh | -0.019733 |
| G front sup | right | CAr | -0.019722 |
| S collat transv post | left | CAr | -0.019266 |
| S oc sup and transversal | left | CAr | -0.019219 |
| G oc-temp lat-fusifor | left | CVol | -0.019145 |
| G front inf-Triangul | left | SDep | -0.019122 |
| G oc-temp lat-fusifor | right | CTh | -0.018994 |
| S circular insula ant | left | CAr | -0.018558 |
| S precentral-inf-part | left | SDep | -0.017945 |
| G front sup | right | CVol | -0.016638 |
| S temporal transverse | right | SDep | -0.016144 |
| S temporal inf | right | CAr | -0.015611 |
| G parietal sup | left | CVol | -0.015111 |
| G oc-temp med-Lingual | left | CTh | -0.014902 |
| S oc middle and Lunatus | left | CTh | -0.014665 |
| S occipital ant | left | CVol | -0.014433 |
| G and S subcentral | right | SDep | -0.014414 |
| S oc sup and transversal | right | CVol | -0.014359 |
| S precentral-inf-part | right | CAr | -0.014357 |
| S postcentral | left | CAr | -0.013485 |
| G cingul-Post-ventral | right | CVol | -0.013257 |
| G oc-temp med-Lingual | left | CVol | -0.012777 |
| G oc-temp med-Lingual | right | CVol | -0.012602 |
| S orbital-H Shaped | left | SDep | -0.012581 |
| S precentral-inf-part | right | CVol | -0.011925 |
| S collat transv post | right | CVol | -0.011797 |
| G front sup | left | CAr | -0.011499 |
| S front middle | right | CTh | -0.011386 |
| S calcarine | right | CVol | -0.01112 |
| G and S cingul-Mid-Post | left | CAr | -0.010035 |
| S intrapariet and P trans | left | CVol | -0.0096053 |
| G temp sup-Lateral | right | CAr | -0.0092356 |
| S temporal transverse | right | CTh | -0.0062906 |
| G and S paracentral | left | SDep | -0.0056593 |
| **DHEA** | | | |
| Intercept | - | - | 8.6577e-08 |
| Pole occipital | left | CTh | 0.24049 |
| G front inf-Opercular | right | SDep | 0.24023 |
| G front middle | left | SDep | 0.23196 |
| G precentral | left | SDep | 0.23131 |
| G Ins lg and S cent ins | right | SDep | 0.23071 |
| Pole occipital | left | CVol | 0.1703 |
| G front inf-Opercular | left | SDep | 0.16941 |
| S occipital ant | right | SDep | 0.16806 |
| S collat transv ant | left | CTh | 0.15805 |
| S occipital ant | left | SDep | 0.15675 |
| Lat Fis-ant-Vertical | left | CTh | 0.15121 |
| G temp sup-G T transv | left | SDep | 0.15018 |
| Pallidum | right | SCVol | 0.14871 |
| G pariet inf-Supramar | left | SDep | 0.12789 |
| Accumbens area | left | SCVol | 0.12479 |
| G temp sup-G T transv | right | SDep | 0.11663 |
| G front inf-Triangul | right | CTh | 0.11529 |
| G temporal inf | left | CTh | 0.10963 |
| Lat Fis-post | right | CAr | 0.10654 |
| Pallidum | left | SCVol | 0.1043 |
| G and S subcentral | left | CAr | 0.098242 |
| G and S frontomargin | right | SDep | 0.095841 |
| S circular insula ant | right | SDep | 0.093413 |
| G front inf-Triangul | left | CTh | 0.091568 |
| G rectus | left | SDep | 0.08702 |
| G oc-temp med-Parahip | right | CAr | 0.086975 |
| G occipital sup | left | CTh | 0.085743 |
| S circular insula inf | left | CVol | 0.084139 |
| S interm prim-Jensen | right | CAr | 0.073672 |
| S collat transv ant | right | CVol | 0.05973 |
| G and S cingul-Mid-Post | right | SDep | -0.30075 |
| G and S transv frontopol | left | CTh | -0.28878 |
| S precentral-inf-part | left | SDep | -0.2731 |
| G precuneus | left | CAr | -0.2423 |
| Lat Fis-post | right | CTh | -0.23136 |
| G front inf-Triangul | right | SDep | -0.21757 |
| S front middle | right | CTh | -0.2169 |
| S temporal inf | right | CVol | -0.21465 |
| G subcallosal | left | SDep | -0.21032 |
| G precuneus | left | SDep | -0.20737 |
| G precuneus | left | CVol | -0.20434 |
| S oc-temp lat | right | SDep | -0.20331 |
| G and S cingul-Ant | right | SDep | -0.1988 |
| G and S frontomargin | left | CTh | -0.19083 |
| S oc middle and Lunatus | left | CTh | -0.19017 |
| S oc middle and Lunatus | right | CVol | -0.18106 |
| S interm prim-Jensen | right | CTh | -0.17273 |
| S collat transv post | right | SDep | -0.17152 |
| G temp sup-Lateral | right | SDep | -0.17109 |
| G pariet inf-Angular | left | SDep | -0.16969 |
| G cuneus | right | SDep | -0.16404 |
| S oc-temp lat | left | CVol | -0.16403 |
| G occipital middle | left | CVol | -0.16346 |
| S precentral-inf-part | right | CTh | -0.16002 |
| G and S occipital inf | left | CAr | -0.15919 |
| G temp sup-Plan polar | left | CVol | -0.15688 |
| S orbital lateral | left | CTh | -0.15658 |
| G parietal sup | left | CAr | -0.15653 |
| G occipital middle | left | CAr | -0.15222 |
| G and S transv frontopol | right | CVol | -0.15199 |
| S orbital lateral | right | CTh | -0.14887 |
| S occipital ant | left | CVol | -0.14836 |
| S oc middle and Lunatus | left | CVol | -0.14806 |
| G temp sup-Plan tempo | left | SDep | -0.1451 |
| G temporal middle | right | CVol | -0.14397 |
| S subparietal | left | CVol | -0.13852 |
| S precentral-inf-part | right | SDep | -0.13567 |
| S orbital-H Shaped | left | SDep | -0.1348 |
| S front middle | left | CTh | -0.13454 |
| S oc middle and Lunatus | right | CAr | -0.13236 |
| S oc sup and transversal | left | CTh | -0.13225 |
| G occipital sup | left | SDep | -0.13059 |
| S oc middle and Lunatus | left | CAr | -0.13033 |
| G front sup | right | CVol | -0.13014 |
| S subparietal | left | CAr | -0.12695 |
| S suborbital | right | CTh | -0.12679 |
| G temp sup-Plan polar | left | CAr | -0.12547 |
| S calcarine | right | CTh | -0.12485 |
| S precentral-inf-part | right | CVol | -0.12433 |
| S occipital ant | right | CVol | -0.12043 |
| S temporal inf | right | CAr | -0.11877 |
| G temp sup-Lateral | right | CAr | -0.11601 |
| S oc-temp lat | left | CAr | -0.11491 |
| G postcentral | left | CAr | -0.11303 |
| G and S subcentral | right | CTh | -0.11172 |
| S temporal sup | right | CVol | -0.11158 |
| S oc sup and transversal | right | CAr | -0.11143 |
| G orbital | left | CTh | -0.11142 |
| G temp sup-Lateral | left | SDep | -0.10962 |
| G Ins lg and S cent ins | right | CVol | -0.10909 |
| S temporal sup | right | CTh | -0.10896 |
| G and S transv frontopol | right | CTh | -0.10865 |
| G front sup | right | CAr | -0.10809 |
| G oc-temp lat-fusifor | left | CVol | -0.10801 |
| G temp sup-Plan polar | right | CVol | -0.10783 |
| S circular insula ant | right | CTh | -0.1077 |
| G cuneus | right | CAr | -0.10649 |
| G oc-temp lat-fusifor | left | CAr | -0.10515 |
| S collat transv post | right | CAr | -0.099365 |
| G temporal middle | right | CAr | -0.098936 |
| G and S occipital inf | left | SDep | -0.09885 |
| G and S paracentral | right | SDep | -0.09825 |
| G oc-temp lat-fusifor | right | CTh | -0.094349 |
| G parietal sup | left | CVol | -0.094172 |
| G and S transv frontopol | left | CVol | -0.094059 |
| S precentral-inf-part | right | CAr | -0.093963 |
| G insular short | right | CTh | -0.093638 |
| S calcarine | left | SDep | -0.093614 |
| G oc-temp med-Lingual | right | SDep | -0.092852 |
| Lat Fis-post | left | CTh | -0.092754 |
| G temporal middle | left | CVol | -0.091428 |
| S orbital-H Shaped | right | SDep | -0.090305 |
| G temporal inf | left | SDep | -0.090294 |
| G precuneus | right | SDep | -0.089461 |
| S orbital lateral | right | SDep | -0.089205 |
| S parieto occipital | left | CTh | -0.087782 |
| S temporal transverse | right | CTh | -0.087517 |
| S occipital ant | left | CAr | -0.087043 |
| S orbital lateral | left | CVol | -0.08521 |
| Pole temporal | right | SDep | -0.084069 |
| S temporal transverse | right | SDep | -0.081302 |
| G front middle | right | CVol | -0.081066 |
| S cingul-Marginalis | right | CTh | -0.080111 |
| S front sup | right | CAr | -0.077715 |
| G and S occipital inf | right | CAr | -0.075224 |
| G and S occipital inf | left | CVol | -0.0737 |
| G temporal inf | left | CAr | -0.073015 |
| Pole temporal | left | SDep | -0.069792 |
| S front inf | right | CTh | -0.069596 |
| G oc-temp med-Lingual | left | CTh | -0.069432 |
| S precentral-sup-part | left | CVol | -0.068963 |
| S cingul-Marginalis | right | CVol | -0.068751 |
| S postcentral | left | CTh | -0.06078 |
| S orbital-H Shaped | right | CAr | -0.059293 |
| S cingul-Marginalis | right | CAr | -0.056764 |
| S oc sup and transversal | left | CVol | -0.051274 |
| S orbital-H Shaped | right | CVol | -0.049143 |
| G occipital sup | left | CAr | -0.03346 |

###### 3.1.2.2 rsfMRI:

| **Regions** | | **Hemi** | **Features** | | **Beta_coefficient** |
| --- | --- | --- | --- | --- | --- |
| **Estradiol (E2)** | | | | | |
| Intercept | |  |  | | -1.4863e-08 |
| Fronto-parietal network and right hippocampus | | Right | Connectivity between cortical network and subcortical region | | -0.32365 |
| Cingulo-opercular network and left cerebellum | | left | Connectivity between cortical network and subcortical region | | -0.25714 |
| Default mode network and left hippocampus | | left | Connectivity between cortical network and subcortical region | | -0.14719 |
| **Testosterone (Tes)** | | | | | |
| Intercept | - | | | - | 38.8769 |
| salience network and left accumbens area | left | | | Connectivity between cortical network and subcortical region | 1.06537 |
| retrosplenial temporal network and left putamen | left | | | Connectivity between cortical network and subcortical region | 0.810606 |
| sensorymotor hand network and sensory motor mouth network | - | | | Inter cortical network | 0.714474 |
| dorsoattentional network and left caudate | - | | | Connectivity between cortical network and subcortical region | 0.657177 |
| auditory network and cingulo-parietal network | - | | | Inter cortical network | 0.584948 |
| cingulo-opercular network and right caudate | right | | | Connectivity between cortical network and subcortical region | 0.523083 |
| sensorymotor mouth network and sensorymotor hand network | - | | | Inter cortical network | 0.202787 |
| default mode network and left hippocampus | left | | | Connectivity between cortical network and subcortical region | -0.781374 |
| sensorymotor hand network and right accumbens area | right | | | Connectivity between cortical network and subcortical region | -0.610923 |
| fronto-parietal network and right hippocampus | right | | | Connectivity between cortical network and subcortical region | -0.562499 |
| ventro-attentional network and left hippocampus | left | | | Connectivity between cortical network and subcortical region | -0.52458 |
| cingulo-parietal network and default mode network | - | | | Inter cortical network | -0.445936 |
| cingulo-opercular and left cerebellum | left | | | Connectivity between cortical network and subcortical region | -0.420209 |
| default mode network and ventro-attentional network | - | | | Inter cortical network | -0.393598 |
| dorso-attentional network and left hippocampus | left | | | Connectivity between cortical network and subcortical region | -0.341585 |
| retrosplenial temporal network and retrosplenial temporal network | - | | | Intra cortical network | -0.332742 |
| salience network and visual network | - | | | Inter cortical network | -0.276198 |
| dorso-attentional network and right accumbens area | right | | | Connectivity between cortical network and subcortical region | -0.268577 |
| auditory network and dorso-attentional network | - | | | Inter cortical network | -0.231274 |
| sensorymotor mouth network and visual network | - | | | Inter cortical network | -0.140874 |
| ventro-attentional network and default mode network | - | | | Inter cortical network | -0.0996058 |
| default mode network and cingulo-opercular network | - | | | Inter cortical network | -0.0887756 |
| visual network and sensory motor mouth network | - | | | Inter cortical network | -0.0237295 |
| **DHEA** | | | | | |
| Intercept | - | | | - | 8.5114e-08 |
| retrosplenial temporal network and left amygdala | left | | | Connectivity between cortical network and subcortical region | 0.39975 |
| cingulo-opercular and left cerebellum | left | | | Connectivity between cortical network and subcortical region | -0.74572 |
| fronto-parietal network and right hippocampus | right | | | Connectivity between cortical network and subcortical region | -0.71897 |
| sensorymotor hand network and left caudate | left | | | Connectivity between cortical network and subcortical region | -0.68702 |
| sensorymotor mouth network and right accumbens area | right | | | Connectivity between cortical network and subcortical region | -0.5363 |
| dorso-attentional network and left cerebellum | left | | | Connectivity between cortical network and subcortical region | -0.51043 |
| default mode network and left hippocampus | left | | | Connectivity between cortical network and subcortical region | -0.39356 |

###### 3.1.2.3 dMRI:

| **Regions** | **Hemi** | **Features** | **Beta_coefficient** |
| --- | --- | --- | --- |
| **Testosterone (Tes)** | | | |
| Intercept | - | - | -1.8782e-08 |
| FXLH | Left | FA | -0.42486 |
| **DHEA** | | | |
| Intercept |  |  | 9.3195e-08 |
| FXLH | Left | FA | -1.1895 |

##### 3.2 Multimodality models:

###### 3.2.1 without Age-regress models:

| **Hormones** | **Selected Features from Multimodality** | **Selected Features from Unimodality** |
| --- | --- | --- |
| E2 | 25 | 26 |
| Tes | 68 | 85 |
| DHEA | 52 | 100 |

##### 3.2.1.1 E2

| **region** | **hemi** | **Features** | **Beta_coefficient** |
| --- | --- | --- | --- |
| Intercept | - | - | 0.98971 |
| FMIN | - | MD | 0.032236 |
| SCSLH | left | FA | 0.02981 |
| default mode network and left thalamus proper | left | Connectivity between cortical network and subcortical region | -0.029072 |
| S collat transv ant | right | CAr | 0.027012 |
| pallidum | right | SubCVol | 0.026774 |
| G precentral | left | CAr | -0.025208 |
| retrosplenial temporal network and left putamen | left | Connectivity between cortical network and subcortical region | 0.025187 |
| G and S frontomargin | left | CTh | -0.024837 |
| G and S cingul-Mid-Post | left | CAr | -0.024449 |
| G and S transv frontopol | left | CTh | -0.023835 |
| Lat Fis-ant-Vertical | left | CTh | 0.023799 |
| G front inf-Orbital | left | SDep | -0.023576 |
| G temp sup-Plan tempo | left | SDep | -0.023249 |
| cingulo-opercular and left cerebellum | left | Connectivity between cortical network and subcortical region | -0.022429 |
| G and S paracentral | left | SDep | -0.021444 |
| G temporal inf | left | SDep | -0.020665 |
| DMRI_DTIMD_FIBERAT_CGCLH | left | Connectivity between cortical network and subcortical region | -0.016832 |
| G and S cingul-Mid-Post | right | SDep | -0.016594 |
| G and S occipital inf | left | CAr | -0.015779 |
| G oc-temp lat-fusifor | left | CVol | -0.015039 |
| sensorymotor hand network and left caudate | left | Connectivity between cortical network and subcortical region | -0.014941 |
| S circular insula ant | right | CTh | -0.014566 |
| S oc-temp med and Lingual | right | CTh | -0.01328 |
| G and S frontomargin | right | CTh | -0.008067 |
| Lat Fis-post | right | CTh | -0.0079107 |

###### 3.2.1.2 Tes

| **region** | **hemi** | **Features** | **Beta_coefficient** |
| --- | --- | --- | --- |
| Intercept | - | - | 38.8766 |
| FMIN | - | MD | 2.62132 |
| CC | - | MD | -2.2739 |
| IFOLH | Left | MD | 1.70981 |
| PSLFLH | Left | MD | -1.57141 |
| SCSLH | Left | FA | 1.43962 |
| Pole occipital | left | CTh | 1.38644 |
| Corpus callosum | - | FA | -1.38443 |
| S calcarine | right | CTh | -1.26238 |
| S precentral-inf-part | right | SDep | -1.24937 |
| G postcentral | left | CAr | -1.18257 |
| salience network and Left accumbens area | left | Connectivity between cortical network and subcortical region | 1.16133 |
| S temporal transverse | left | CTh | 1.11319 |
| frontoparietal network and right hippocampus | right | Connectivity between cortical network and subcortical region | -1.07549 |
| G precentral | right | CTh | 1.06292 |
| dorsoattentional network and left caudate | left | Connectivity between cortical network and subcortical region | 1.01588 |
| hippocampus | left | SubCVol | 0.957272 |
| cingulo-opercular network and left cerebellum | left | Connectivity between cortical network and subcortical region | -0.935823 |
| G cingul-Post-ventral | right | CTh | -0.92704 |
| S subparietal | left | CTh | -0.9151 |
| auditory network and cingulo-parietal network | - | Inter cortical network | 0.884952 |
| G temp sup-Lateral | right | SDep | -0.884649 |
| cerebellum cortex | right | SubCVol | 0.875675 |
| G and S occipital inf | left | CAr | -0.85131 |
| FMAJ | - | FA | 0.845868 |
| retrosplenial temporal network and left putamen | left | Connectivity between cortical network and subcortical region | 0.809896 |
| S precentral-sup-part | right | CTh | 0.805264 |
| G temp sup-Plan polar | right | CTh | 0.800723 |
| G Ins lg and S cent ins | right | SDep | 0.776916 |
| salience network and visual network | - | Inter cortical network | -0.773066 |
| G precuneus | left | CVol | -0.764576 |
| sensorymotor hand network and sensory motor mouth network | - | Inter cortical network | 0.757902 |
| FXLH | left | FA | -0.753454 |
| G and S transv frontopol | right | CVol | -0.731702 |
| default network and left hippocampus | left | Connectivity between cortical network and subcortical region | -0.731615 |
| G front inf-Orbital | left | SDep | -0.712321 |
| CGHLH | left | FA | 0.696944 |
| thalamus proper | left | SubCVol | 0.693191 |
| G subcallosal | left | CTh | -0.667971 |
| retrosplenial temporal network and retrosplenial temporal network | - | Intra cortical network | -0.654803 |
| FXRH | right | FA | -0.646693 |
| cingulo-parietal network and default mode network | - | Inter cortical network | -0.645821 |
|  | right | MD | -0.629097 |
| S oc-temp med and Lingual | right | SDep | -0.626376 |
|  | left | FA | 0.611495 |
| Lat Fis-post | left | CTh | -0.609425 |
| G oc-temp med-Lingual | right | CTh | -0.60176 |
| G and S cingul-Ant | right | SDep | -0.597709 |
| PSCSRH | right | FA | -0.583785 |
| IFSFCLH | left | FA | -0.573449 |
| Lat Fis-post | right | CTh | -0.558914 |
| G and S transv frontopol | left | CTh | -0.545581 |
| Pallidum | right | SubCVol | 0.543308 |
| G and S frontomargin | left | CTh | -0.531576 |
| cingulo-opercular network and right caudate | right | Connectivity between cortical network and subcortical region | 0.529893 |
| auditory network and dorso-attentional network |  | Inter cortical network | -0.528508 |
| CGHRH | right | MD | -0.524626 |
| sensorymotor hand network and right accumbens | right | Connectivity between cortical network and subcortical region | -0.507986 |
| default network and ventroattentional network |  | Inter cortical network | -0.504025 |
| S suborbital | right | CTh | -0.459564 |
| FXCUTLH | left | FA | -0.43549 |
| G front middle | left | SDep | 0.434733 |
| CGCLH | left | FA | 0.433582 |
| sensorymotor mouth network and visual network | - | Inter cortical network | -0.390717 |
| UNCLH | left | FA | -0.386637 |
| IFSFCRH | right | FA | -0.368554 |
| S orbital-H Shaped | left | SDep | -0.362471 |
| G oc-temp lat-fusifor | left | CTh | -0.358055 |
| SIFCLH | left | FA | 0.326733 |
| dorso-attentional network and right accumbens area | right | Connectivity between cortical network and subcortical region | -0.321757 |
| G and S cingul-Ant | left | CTh | -0.292062 |

###### 3.2.1.3 DHEA

| **region** | **hemi** | **Features** | **Beta_coefficient** |
| --- | --- | --- | --- |
| Intercept | - | - | -3.1846e-08 |
| G and S cingul-Mid-Post | right | SDep | -2.4852 |
| Pole occipital | left | CVol | 2.2324 |
| G occipital sup | left | CTh | 2.0587 |
| S collat transv ant | left | CTh | 1.9058 |
| S collat transv ant | right | CVol | 1.8601 |
| G and S transv frontopol | left | CTh | -1.8536 |
| G Ins lg and S cent ins | right | SDep | 1.8147 |
| cingulo-opercular network and left cerebellum | left | Connectivity between cortical network and subcortical region | -1.8113 |
| sensorymotor mouth network and right accumbens area | right | Connectivity between cortical network and subcortical region | -1.7901 |
| accumbens area | left | SubCVol | 1.7896 |
| S oc-temp lat | right | SDep | -1.7272 |
| G precuneus | left | CAr | -1.6992 |
| Fornix | left | FA | -1.6907 |
| frontoparietal network and right hippocampus | right | Connectivity between cortical network and subcortical region | -1.6692 |
| G and S subcentral | left | CAr | 1.6071 |
| Lat Fis-ant-Vertical | left | CTh | 1.5593 |
| S oc middle and Lunatus | left | CTh | -1.5575 |
| retrosplenial temporal network and left amygdala | left | Connectivity between cortical network and subcortical region | 1.5197 |
| G front middle | left | SDep | 1.426 |
| Lat Fis-post | right | CTh | -1.3535 |
| G oc-temp med-Lingual | right | SDep | -1.3512 |
| S temporal inf | right | CVol | -1.3484 |
| G front inf-Triangul | left | CTh | 1.2954 |
| G pariet inf-Angular | left | SDep | -1.2608 |
| S collat transv post | right | SDep | -1.2459 |
| G subcallosal | left | SDep | -1.227 |
| G temp sup-Plan polar | right | CVol | -1.2248 |
| sensorymotor hand network and left caudate | left | Connectivity between cortical network and subcortical region | -1.2187 |
| G front inf-Opercular | right | SDep | 1.2074 |
| G and S occipital inf | left | CAr | -1.0987 |
| S interm prim-Jensen | right | CTh | -1.0786 |
| G Ins lg and S cent ins | right | CVol | -1.0727 |
| G and S cingul-Ant | right | SDep | -1.0581 |
| G precuneus | left | SDep | -1.0518 |
| G front inf-Triangul | right | SDep | -1.0458 |
| S front middle | right | CTh | -1.0451 |
| G temporal inf | left | CTh | 1.0398 |
| S precentral-inf-part | left | SDep | -1.0367 |
| G front inf-Triangul | right | CTh | 1.0193 |
| S circular insula inf | left | CVol | 1.0179 |
| G temp sup-G T transv | left | SDep | 1.013 |
| S occipital ant | right | SDep | 0.96373 |
| G temp sup-G T transv | right | SDep | 0.89167 |
| dorsoattentional network and left cerebellum | left | Connectivity between cortical network and subcortical region | -0.8421 |
| G temp sup-Plan polar | left | CVol | -0.83798 |
| G cuneus | right | SDep | -0.75792 |
| S orbital lateral | left | CTh | -0.75305 |
| S occipital ant | left | SDep | 0.74026 |
| G and S frontomargin | left | CTh | -0.71199 |
| S suborbital | right | CTh | -0.6093 |
| S orbital lateral | right | SDep | -0.5443 |
| S precentral-inf-part | right | CVol | -0.53811 |

###### 3.2.2 Age-regress models:

| **Hormones** | **Selected Features from Multimodality** | **Selected Features from Unimodality** |
| --- | --- | --- |
| E2 | 9 | 9 |
| Tes | 53 | 141 |
| DHEA | 83 | 146 |

##### 3.2.2.1 E2

| **region** | **Hemi** | **Features** | **Beta_coefficient** |
| --- | --- | --- | --- |
| Intercept | - | - | -1.3119e-10 |
| cingulo-opercular and left cerebellum | left | Connectivity between cortical network and subcortical region | -0.026641 |
| G temp sup-Plan tempo | left | SDep | -0.019616 |
| G and S transv frontopol | left | CTh | -0.018556 |
| G and S frontomargin | left | CTh | -0.018421 |
| G oc-temp lat-fusifor | left | CVol | -0.017176 |
| fronto-parietal network and right hippocampus | right | Connectivity between cortical network and subcortical region | -0.012863 |
| default mode network and left hippocampus | left | Connectivity between cortical network and subcortical region | -0.010537 |
| cingulo-opercular and left cerebellum | left | Connectivity between cortical network and subcortical region | -0.020787 |
| FXCUTRH | right | MD | 0.018483 |

###### 3.2.2.2 Tes

| **region** | **hemi** | **Features** | **Beta_coefficient** |
| --- | --- | --- | --- |
| Intercept | - | - | -2.2598e-09 |
| cingulo-opercular and left cerebellum | left | Connectivity between cortical network and subcortical region | -0.66444 |
| fronto-parietal network and right hippocampus | right | Connectivity between cortical network and subcortical region | -0.60981 |
| FXLH | left | FA | -0.54595 |
| Pole occipital | left | CTh | 0.50056 |
| dorsoattentional network and left caudate | left | Connectivity between cortical network and subcortical region | 0.49023 |
| S subparietal | right | CVol | 0.46342 |
| salience network and left accumbens area | left | Connectivity between cortical network and subcortical region | 0.45299 |
| cerebellum cortex | right | SubCVol | 0.45189 |
| G Ins lg and S cent ins | right | SDep | 0.44666 |
| G precuneus | right | CTh | 0.42089 |
| G and S transv frontopol | left | CTh | -0.39176 |
| S precentral-inf-part | right | SDep | -0.38956 |
| G oc-temp lat-fusifor | left | CTh | -0.3835 |
| Lat Fis-ant-Vertical | left | CTh | 0.3827 |
| G and S cingul-Mid-Post | right | SDep | -0.37695 |
| G occipital sup | left | CTh | 0.37068 |
| G subcallosal | left | CTh | -0.33582 |
| G temp sup-Lateral | right | SDep | -0.32735 |
| thalamus proper | right | SubCVol | 0.32733 |
| G and S cingul-Ant | right | SDep | -0.32427 |
| S circular insula inf | left | CAr | 0.32389 |
| Lat Fis-ant-Horizont | left | CVol | 0.31585 |
| G pariet inf-Angular | left | CTh | 0.30988 |
| G temp sup-Plan polar | right | CTh | 0.30235 |
| S intrapariet and P trans | left | SDep | 0.30044 |
| S calcarine | right | CTh | -0.29223 |
| default mode network and left hippocampus | left | Connectivity between cortical network and subcortical region | -0.28943 |
| G front inf-Triangul | left | CTh | 0.28747 |
| retrosplenial temporal network and left putamen | left | Connectivity between cortical network and subcortical region | 0.28119 |
| G precuneus | left | SDep | -0.28045 |
| S interm prim-Jensen | right | CTh | -0.27964 |
| S oc-temp lat | right | SDep | -0.26419 |
| Lat Fis-post | left | CTh | -0.26318 |
| S subparietal | left | CTh | -0.25568 |
| G temp sup-G T transv | right | SDep | 0.25241 |
| S circular insula ant | right | CTh | -0.2397 |
| G front inf-Triangul | right | CTh | 0.22494 |
| G temp sup-Plan tempo | left | SDep | -0.22201 |
| G and S frontomargin | left | CTh | -0.21499 |
| G and S transv frontopol | right | CTh | -0.21365 |
| S circular insula ant | left | CAr | -0.20923 |
| G precuneus | left | CVol | -0.2002 |
| S pericallosal | right | SDep | 0.19586 |
| cingulo-opercular network and right caudate | right | Connectivity between cortical network and subcortical region | 0.1941 |
| G and S occipital inf | left | CAr | -0.19282 |
| G cingul-Post-ventral | right | CTh | -0.18876 |
| G front middle | left | SDep | 0.18667 |
| G pariet inf-Angular | left | SDep | -0.18409 |
| S oc sup and transversal | right | CAr | -0.1767 |
| S circular insula ant | right | SDep | 0.17065 |
| G front inf-Triangul | right | SDep | -0.16742 |
| retrosplenial temporal network and retrosplenial temporal network | - | Intra cortical network | -0.16333 |
| G and S cingul-Mid-Post | left | CAr | -0.14901 |

###### 3.2.2.3 DHEA

| **region** | **Hemi** | **Features** | **Beta_coefficient** |
| --- | --- | --- | --- |
| Intercept | - | - | -3.1846e-08 |
| G and S cingul-Mid-Post | right | SD | -2.4852 |
| Pole occipital | left | CVol | 2.2324 |
| G occipital sup | left | CTh | 2.0587 |
| S collat transv ant | left | CTh | 1.9058 |
| S collat transv ant | right | CVol | 1.8601 |
| G and S transv frontopol | left | CTh | -1.8536 |
| G Ins lg and S cent ins | right | SDep | 1.8147 |
| cingulo-opercular network and left cerebellum | left | Connectivity between cortical network and subcortical region | -1.8113 |
| somatomotor motor network and right accumbens area | right | Connectivity between cortical network and subcortical region | -1.7901 |
| accumbens area | left | SCVol | 1.7896 |
| S oc-temp lat | right | SDep | -1.7272 |
| G precuneus | left | CAr | -1.6992 |
| FXLH | left | FA | -1.6907 |
| frontoparietal network and right hippocampus | right | Connectivity between cortical network and subcortical region | -1.6692 |
| G and S subcentral | left | CAr | 1.6071 |
| Lat Fis-ant-Vertical | left | CTh | 1.5593 |
| S oc middle and Lunatus | left | CTh | -1.5575 |
| retrosplenial temporal network and left amygdala | left | Connectivity between cortical network and subcortical region | 1.5197 |
| G front middle | left | SDep | 1.426 |
| Lat Fis-post | right | CTh | -1.3535 |
| G oc-temp med-Lingual | right | SDep | -1.3512 |
| S temporal inf | right | CVol | -1.3484 |
| G front inf-Triangul | left | CTh | 1.2954 |
| G pariet inf-Angular | left | SDep | -1.2608 |
| S collat transv post | right | SDep | -1.2459 |
| G subcallosal | left | SDep | -1.227 |
| G temp sup-Plan polar | right | CVol | -1.2248 |
| somatomotor hand network and left caudate | left | Connectivity between cortical network and subcortical region | -1.2187 |
| G front inf-Opercular | right | SDep | 1.2074 |
| G and S occipital inf | left | CAr | -1.0987 |
| S interm prim-Jensen | right | CTh | -1.0786 |
| G Ins lg and S cent ins | right | CVol | -1.0727 |
| G and S cingul-Ant | right | SDep | -1.0581 |
| G precuneus | left | SDep | -1.0518 |
| G front inf-Triangul | right | SDep | -1.0458 |
| S front middle | right | CTh | -1.0451 |
| G temporal inf | left | CTh | 1.0398 |
| S precentral-inf-part | left | SDep | -1.0367 |
| G front inf-Triangul | right | CTh | 1.0193 |
| S circular insula inf | left | CVol | 1.0179 |
| G temp sup-G T transv | left | SDep | 1.013 |
| S occipital ant | right | SDep | 0.96373 |
| G temp sup-G T transv | right | SDep | 0.89167 |
| dorsoattentional network and left cerebellum | left | Connectivity between cortical network and subcortical region | -0.8421 |
| G temp sup-Plan polar | left | CVol | -0.83798 |
| G cuneus | right | SDep | -0.75792 |
| S orbital lateral | left | CTh | -0.75305 |
| S occipital ant | left | SDep | 0.74026 |
| G and S frontomargin | left | CTh | -0.71199 |
| S suborbital | right | CTh | -0.6093 |
| S orbital lateral | right | SDep | -0.5443 |
| S precentral-inf-part | right | CVol | -0.53811 |

#### C) Regressing caffeine from Tes hormone:

Only showing the different features

| **Features without regressing caffeine intake** | **Features after regressing caffeine intake** |
| --- | --- |
| cortical thickness in left transverse temporal sulcus | cortical volume in right superior parietal lobule |
| FA in FMIN | sulcal depth in left middle-anterior part of the cingulate gyrus and sulcus |
| Left thalamus proper volume | cortical thickness in right opercular part of the inferior frontal gyrus |
| Accumbens area volume | sulcal depth in left lateral occipito-temporal gyrus |
| Connectivity between auditory network and left caudate |  |
| FA in FXLH |  |
| Connectivity between visual attention network and left hippocampus |  |
| FA in TSLFLH |  |

#### D) Regressing collection time from E2 levels:

Only showing different features

| **Features without regressing collection time** | **Features after regressing collection time** |
| --- | --- |
| Cortical thickness in right Lat Fis-post | Cortical thickness in left subparietal |
| Subcortical volume in right pallidum | Cortical thickness in left S oc sup and transversal |
| Cortical thickness in left S oc middle and Lunatus | MD in right FXCUT |
| Sulcal depth in right G temporal middle | FA in left CGC |
| FA in right UNC |  |
| MD in left FXCUT |  |
